# Projecting spatiotemporal bioclimatic niche dynamics of endemic Pyrenean plant species under climate change: how much will we lose?

**DOI:** 10.1101/2025.03.19.644085

**Authors:** Noèmie Collette, Sébastien Pinel, Valérie Delorme-Hinoux, Joris A.M. Bertrand

## Abstract

Species distributions are shifting under global change, with mountain ecosystems among the most vulnerable. In such landscapes, ability to track changing conditions is limited, threatening narrowly distributed species. As a mountain biodiversity hotspot in southwestern Europe, the Pyrenees harbors many such species, making it a key case study for climate vulnerability assessments.

This study implements a bioclimatic niche modeling pipeline to evaluate climate change impact on endemic Pyrenean plant species by 2100. Objectives are to (i) map current bioclimatic niche suitability, (ii) forecast its future spatial dynamics, and (iii) identify potential climate refugia for conservation. Species occurrences were combined with 19 bioclimatic variables (1x1 km resolution) to characterize bioclimatic niche suitability, using an ensemble modeling approach integrating five algorithms (MaxEnt, Generalized Linear Model, Generalized Additive Model, Gradient Boosting Machine, and Random Forest). Their future spatiotemporal dynamics were projected under four climate scenarios (Shared Socioeconomic pathways 126, 245, 370, 585) for four successive periods spanning 2021 to 2100.

By 2100, 69% of endemic species are projected to lose over 75% of their bioclimatic niche, and half to face complete losses under high-emission scenarios. Only two species may gain suitable areas, highlighting the need for species-specific conservation strategies. Bioclimatic niches are projected to shift by ∼180 m upslope and ∼3 km in latitude on average, with areas of highest multi-species suitability, referred to as bioclimatic hotspots, becoming restricted to elevation above 2000 m. These trends intensify after 2041-2060 period, reflecting escalating climate pressures as the century progresses.

Our findings highlight the profound threat climate change may pose to endemic Pyrenean flora, with widespread bioclimatic niche losses projected by the century’s end and high elevation refugia emerging as key conservation priorities. Anticipating these shifts and integrating them into conservation planning will be crucial to mitigating high-elevation biodiversity loss in a rapidly changing world.

## 1 Introduction

Climate change represents one of the greatest environmental challenges of our time, with a significant and far-reaching impact on world’s ecosystems (Intergovernmental Panel On Climate Change, 2022 ; Li *et al*., 2018). Among these ecosystems, mountain regions are of special concern as they are experiencing some of the highest and fastest rates of warming, a phenomenon widely recognized as elevation-dependent warming (Pepin *et al*., 2015). This refers to the tendency for warming rates at higher altitudes to surpass those observed in lowland regions, a pattern identified across multiple mountainous regions worldwide (Pepin *et al*., 2022 ; Qixiang *et al*., 2018). Such warming raises significant concerns for mountain ecosystems, which are particularly sensitive to these rapid changes (Intergovernmental Panel on Climate Change (IPCC), 2023 ; Knight, 2022). A cascade of ecological consequences follows (Dainese *et al*., 2024 ; Rogora *et al*., 2018), including shifts in species distribution, a well-documented response to climate change (Lawlor *et al*., 2024 ; Pearse & Davies, 2019 ; Rubenstein *et al*., 2023).

Climate velocity describes the speed and direction in which a species needs to move to stay within its climatic niche as climate conditions change, which, in mountainous contexts, forces species to occupy progressively higher elevations (Chan *et al*., 2024), resulting in shifts upward in their core distribution (Freeman *et al*., 2018). However, these elevational shifts are unlikely to keep pace with the rapidly changing climate (Carroll *et al*., 2015 ; Goodwin *et al*., 2025 ; Kellner *et al*., 2023). Mountain plants, in particular, face even greater challenges than other organisms due to their limited dispersal capacity (Di Musciano *et al*., 2020 ; Morgan & Venn, 2017). Their migration is further constrained by the complex topography of mountainous landscapes and habitat fragmentation (Bueno de Mesquita *et al*., 2018 ; Elsen *et al*., 2020), which create substantial barriers to movement by breaking the spatial continuity of suitable areas. Even when they successfully reach climatically suitable habitat, or are already located there, establishment and persistence may be constrained by several factors. Biotic mismatches with pollinators (Dainese *et al*., 2024 ; Marshall *et al*., 2020 ; Trunschke *et al*., 2024 ; Vitasse *et al*., 2021), competition with resident or incoming species (Alexander *et al*., 2015 ; Schuchardt *et al*., 2023) or the absence of key mutualists can hinder recruitment. Abiotic constraints, including unsuitable edaphic conditions (Eichel *et al*., 2023 ; Ni & Vellend, 2024) or microtopography may also reduce habitat suitability. Finally, intrinsic species traits such as germination requirements (Alexander *et al*., 2018) can further restrict successful colonization. Moreover, if its establishment occurs within an isolated microclimatic refugia, prolonged isolation of these persistent populations may compromise their long-term survival by limiting genetic mixing, thereby reducing genetic diversity (Pinto *et al*., 2024 ; Turnock *et al*., 2024). This, in turn, limits their adaptability and resilience to environmental changes (Bijlsma & Loeschcke, 2012 ; Frankham, 2005 ; Mathur *et al*., 2023). Such prolonged isolation mirrors patterns observed on a broader scale among mountaintop species, which persist in isolated high-elevation habitats often referred to as “sky island”. Habitat fragmentation and progressive loss of physical space with elevation amplify extinction risk for these species (Elsen & Tingley, 2015), as they lack alternative areas to persist when facing unfavorable conditions or competition with other species. This scenario, referred to as “mountaintop extinction”, primarily threatens species with narrow distribution, residing in the upper limits of latitude gradients, or restricted to highest elevation.

The Pyrenees mountain chain, Europe’s second-largest biodiversity hotspot with 3,652 indigenous vascular plants, is particularly concerning due to the number of mountaintop species, including a panel of endemics and specialized flora (GÓmez *et al*., 2017). The region’s complex climate and geodiversity have historically favored this exceptional diversity (KÖrner & Hiltbrunner, 2021 ; Rahbek *et al*., 2019a) and served as both refuge and corridor for organisms during past climatic changes (GarcÍa *et al*., 2022 ; Schmitt, 2009). However, intensifying global pressures such as climate, land use, and socio-economic changes, including tourism, urban development and agricultural abandonment (OPCC, 2018 ; Zango-Palau *et al*., 2024), may compromise its capacity to buffer future environmental shifts (Rahbek *et al*., 2019b). The upslope shift of plants in the Pyrenees (Ameztegui *et al*., 2016 ; Lenoir *et al*., 2008 ; Pauli *et al*., 2012) appears to contribute to increasing species richness on summits (Camarero *et al*., 2006 ; Steinbauer *et al*., 2018). Yet, this dynamic often comes at the cost of endemic and range-restricted species.

Endemic species represent about 5 % of the Pyrenean flora and species at their upper distribution limits represent another 20 % (GÓmez *et al*., 2017). Although not all of them are strictly confined to high elevations, their limited distribution makes them particularly vulnerable to range contraction. Altogether, up to a quarter of the regional flora may face mountain top extinction scenario in the coming decades. Observations in the Alps, another European mountain range, support the potential mountaintop extinction risk for endemic and range- restricted plant species in the Pyrenees. Red-listed plants have experienced rapid contraction of their elevation margins, resulting in significant habitat loss in lowland areas (Geppert *et al*., 2023), and endemic ones undergoing substantial range reductions, with projections suggesting that 40 % of their current distribution could become climatically unsuitable by the end of the 21^st^ century (Dullinger *et al*., 2012). Similar dynamics in the Pyrenees could drive extinctions, altering summit biodiversity and ecosystems. Although alpine plants show traits promoting resilience, such as adaptation to harsh conditions, clonal growth, and access to diverse thermal microhabitats (KÖrner & Hiltbrunner, 2021), these ongoing dynamics raise concerns about their ability to track or adapt to climatic changes, with potential consequences for mountain biodiversity and ecological communities. In this context, identifying potential responses of species to these changes is essential for anticipating and guiding conservation efforts.

To assess the impact of climate change on plant distributions, the use of modeling tools is a widely adopted approach. Species distribution models (SDMs), most often implemented as correlative approaches, relate species occurrences to environmental variables to predict and describe habitat suitability or probability of presence of a species across spatial and temporal scales (Vasconcelos *et al*., 2024). The robust methodological foundations and predictive power of these models have led to their application across diverse fields, including agriculture, public health, and, notably, conservation biology (Franklin, 2023). In the Pyrenees, SDMs have primarily been used for conservation purposes. Research has explored species distribution drivers (Dalibard *et al*., 2021 ; Kouba *et al*., 2011 ; Williams-Tripp *et al*., 2012), projected climate- driven shifts (de Pous *et al*., 2015 ; Glad & Mallard, 2022 ; MartÍnez *et al*., 2012 ; PÉrez-GarcÍa *et al*., 2013; Russo *et al*., 2023 ; Salvado *et al*., 2025, 2022), and reconstructed distribution over geological time (Bidegaray-Batista *et al*., 2016 ; Charbonnel *et al*., 2016 ; Gutierrez *et al*., 2022). Together, these studies underscore the value of SDMs in informing management and conservation strategies for many species. By providing insights into species-environment relationships and distribution patterns for past, present and future, SDMs become relevant tools for anticipating and addressing climate change impacts, and, more generally, the impact of global change on biodiversity.

Open-access environmental and occurrence databases have expanded possibilities for studying species- environment relationships. Although various tools exist for each SDMs stage (Kass *et al*., 2024), fully integrated pipeline (i.e. framework), especially those suited for rare or range-restricted species, such as endemics, remain scarce. Default settings in existing frameworks are often not fully suited for these taxa and could require manual adjustments that become time-consuming when applied to many species. To address this, we proposed a fully integrated and reproducible SDMs pipeline specifically tailored to species with few occurrences. It automates algorithm-specific tuning, provides auxiliary scripts to guide key decisions (e.g. block size for spatial cross-validation) and used evaluation metrics suited for low-prevalence data (e.g. Boyce Index, AUC-PR). It also incorporates spatial visualization of uncertainties related to both algorithms and climate projections, addressing the frequent lack of uncertainty reporting in SDMs for rare and endemic species (Qazi *et al*., 2022). In addition to streamlining the modeling process, this pipeline produces harmonized outputs across species, enhancing understanding of spatial and temporal biodiversity dynamics and enabling consistent comparisons of climate-driven range shifts, making it well suited for multi-species assessments. Ultimately, it provides natural area managers practical tools to preserve floristic diversity and sustain the ecological resilience of mountain ecosystems in the face of accelerating climate change.

Endemic species, typically restricted to narrow geographical and climatic niches, serve as indicators of regional biodiversity (Lamoreux *et al*., 2006). Their concentration in mountainous areas highlights the key role these regions play in enhancing floristic richness and uniqueness. As irreplaceable components of the region’s natural heritage, their conservation is therefore crucial. In this context, the proposed framework, which enables parallel analysis of SDMs outputs across species, is particularly valuable for identifying areas where several endemic species may overlap and could be most at risk, thus supporting more effective and targeted management efforts. Consequently, using SDMs, this study aims to: (i) map current distribution of bioclimatic suitable areas for 59 endemic Pyrenean plant species, (ii) forecast future extent and dynamics of these areas by the end of the 21^st^ century under multiple climate scenarios, and (iii) identify where these species could find refuge by 2081-2100 period.

## 2 Material and methods

### Study area and species occurrences data

This study follows the mountain delimitation of Körner *et al*. (2011), updated by Snethlage *et al*. (2022). Within this framework, the Pyrenees span ∼40,000 km² along the France-Spain-Andorra border (Figure 1), with over 150 peaks above 3000 m (GÓmez *et al*., 2017). Their steep northern slopes contrast with broader southern valleys, featuring diverse landscapes. At the Atlantic-Mediterranean-continental interface, this mountain range displays diverse climatic conditions. Mean annual temperatures range from >10 °C in lowlands to <4 °C on peaks, with precipitation varying from >2000 mm (Atlantic slopes) to ∼1000 mm (sheltered areas) (Cuadrat *et al*., 2024).

**Figure 1.**
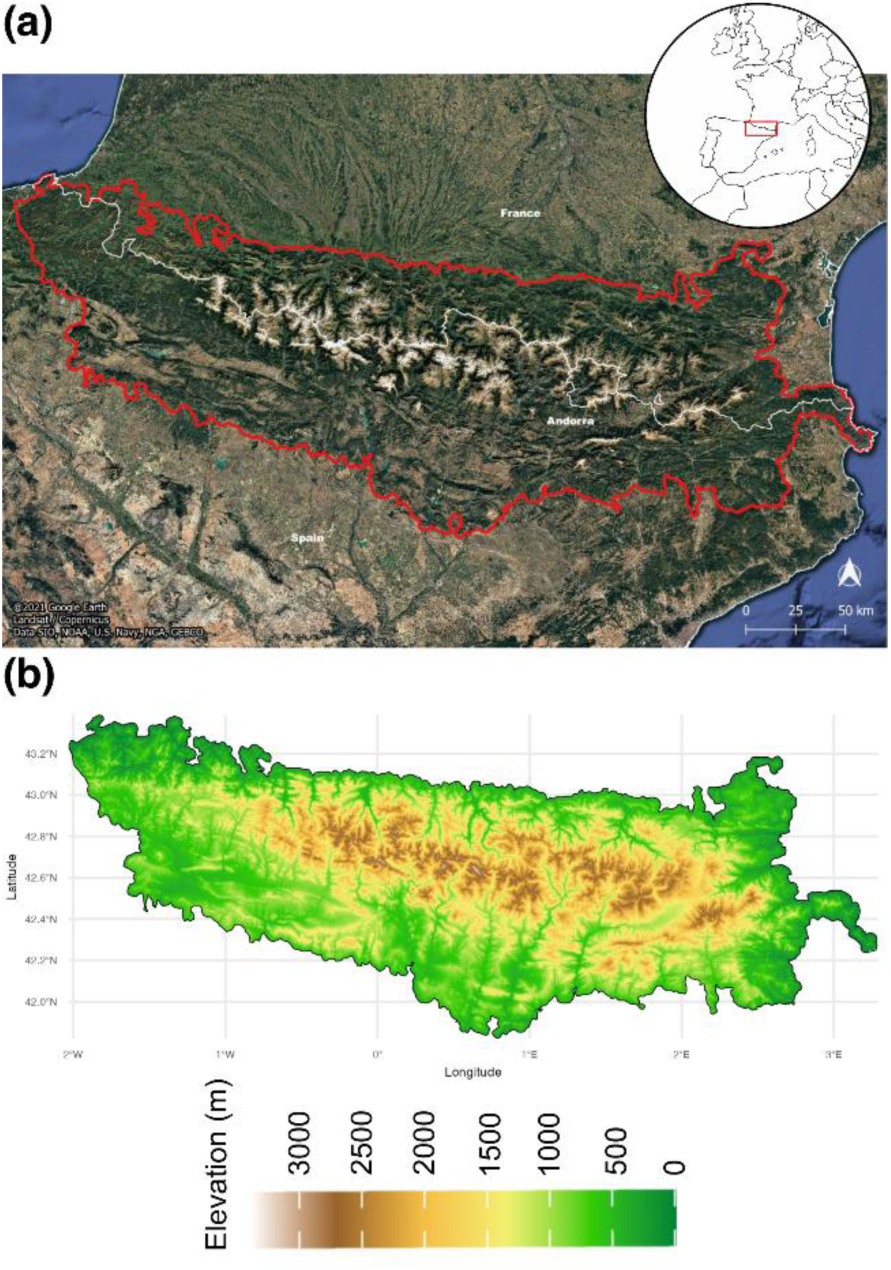
Geographic extent and topography of the Pyrenees. **(a)** The Pyrenees Mountain range (red outline) spans the border regions of France, Spain, and Andorra. Its delimitation follows the definition set out by Global Mountain Biodiversity Assessment (Körner *et al*., 2011, updated by Snethlage *et al*., 2022). The inset map shows the location of the Pyrenees in Europe. Satellite imagery: Google Earth (© Google, Image Landsat/Copernicus, Data SIO, NOAA, U.S. Navy, NGA, GEBCO). **(b)** Elevation gradient from a 90m- resolution digital elevation model (CGIAR-CSI, 2018), showing the region’s topographic variation from lowlands (green) to high peaks (brown to white).

Among the 2,682 species listed in the Atlas of Pyrenean Flora (Pironon *et al*., 2022), 84 were classified as endemic. Excluding endemic subspecies reduced the set to 78 species. Occurrence data were obtained from the Global Biodiversity Information Facility (GBIF.ORG, 2025), supplemented with research-grade data from iNaturalist (https://www.inaturalist.org). Occurrences since 1970 within the study area and ≤ 1 km coordinate uncertainty were retained, fitting the resolution of environmental variables. Spatial thinning retained one occurrence per grid cell. Species with <4 occurrences and/or highly restricted distributions were also removed, as selected SDMs assessment method requires presence of species in each spatial block during cross- validation, and therefore requires sufficient spatial representation. Ultimately, 59 species were included.

### Bioclimatic data

WorldClim 2.1 (Fick & Hijmans, 2017) provided 19 rasterized bioclimatic variables (see Supplemental Information, Appendix A) for the current (1970-2000) and future periods (2021-2040, 2041-2060, 2061-2080, 2081-2100 designated as 2030, 2050, 2070, 2090 in this study). Future datasets based on CMIP6 General Circulation Models (GCMs) at 0.5 arcs/min resolution (∼1x1 km at the equator) were combined into median raster to capture model variability (Carroll & Mahony, 2025 ; Song *et al*., 2024). Projections follow four Shared Socio-economic Pathways (SSP, 126, 245, 370, 585, best to worst-case, respectively), representing socio-economic trajectory and radiative forcing (W.m⁻²) expected in 2100 (O’Neill *et al*., 2017 ; Welch, 2024). Under these scenarios, mean annual temperature in the Pyrenees (9.4 °C) is projected to rise by +18.69 % (11.56°C) to +39.51 % (15.54 °C) in 2081-2100 period, while current annual precipitation (975.41 mm) may decrease by -2.18 % (971.85 mm) to -13.57 % (843 mm) (Figure 2).

**Figure 2.**
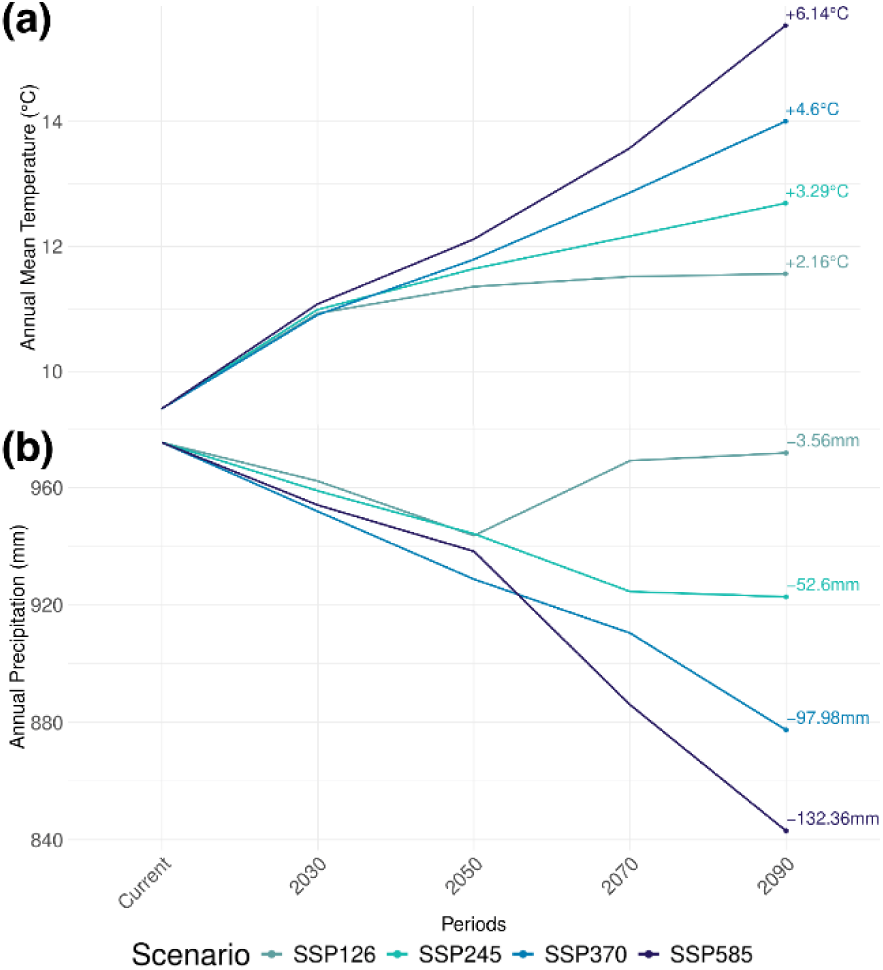
Projected changes in temperature and precipitation in the Pyrenees under Shared Socioeconomic Pathways (SSP) by 2081-2100 period from WorldClim 2.1. **(a)** Mean annual temperature (°C) and **(b)** annual precipitation (mm) projections for 2021-2100 under SSP126, SSP245, SSP370, and SSP585. Values (+/-) reflect differences from the 1970-2000 baseline (current period) for the 2081-2100 period. 2021-2040, 2041-2060, 2061-2080, 2081-2100 periods designated as 2030, 2050, 2070, 2090.

To reduce redundancy, highly positively correlated variables (r > 0.8) at presence locations for each species were clustered. Within each cluster, the most relevant variable per species was retained based on a predefined ecological priority list (Appendix A), which favors variables that are easier to interpret ecologically (e.g. mean annual temperature), followed by variables describing climatic extremes during cold and warm seasons, and finally more derived or complex indices (e.g. temperature seasonality, isothermality). Negatively correlated variables, reflecting complementary environmental gradients, were not excluded. Depending on species, models include between three and eight bioclimatic variables after the selection process. Selected variables for each species are listed in the Appendix A.

### Bioclimatic suitability modeling

#### Model development

A presence/pseudo-absence approach was employed to improve SDMs reliability when true absences are unavailable (Mateo *et al*., 2010). Following recommendations from Barbet-Massin et al., 2012 and Biomod2 team (Gueguen *et al*., 2025), we tested different pseudo-absence configurations they outlined and retained the one that, in our analyses, consistently improved model evaluation scores (Appendix B). The number of pseudo-absences was set equal to the number of presences and sampled within the environmental space, using environmental constraints to restrict sampling to unsuitable conditions (Broussin *et al*., 2024). In line with Biomod2 guidelines, each algorithm was run 10 times with distinct pseudo-absence sets generated separately for each species, thereby reducing background dependence. Each set is spatially thinned to retain a single occurrence per environmental grid cell.

As modeling algorithms significantly impact SDMs outcomes (Buisson *et al*., 2010 ; Thuiller *et al*., 2019) and predictive performance varying across approaches (Norberg *et al*., 2019 ; Valavi *et al*., 2022), bioclimatic suitability was modeled using five algorithms : Maxent (MXT) Generalized Linear Models (GLM) Generalized Additive Models (GAM), Gradient Boosting Machines (GBM) and RandomForest (RF). To ensure comparability across algorithms, probability outputs from GLM, GAM, GBM, and RF were converted into favorability values using the Fav function from the fuzzySim package (Barbosa, 2015), which corrects sample prevalence (presence/absence ratio). This transformation was not applied to MXT outputs, as it generates suitability scores rather than probabilities.

#### Parameters tuning, evaluation and ensemble modeling

An automated workflow tuned model parameters for each iteration and algorithm. For each case, a parameter combination was tested (see Appendix C), and the combination yielding the highest performance score was retained. This score was calculated as a weighted average of three evaluation metrics: the Boyce Index (weight 0.5), AUC-PR (0.25), and Sensitivity (0.25). Greater weight was given to the Boyce Index as it best captures how predicted suitability aligns with observed occurrences, ensuring more reliable projections from sparse data. Each metric captured distinct aspect of model performance : discrimination, classification, and calibration metrics (Sillero *et al*., 2021). Discrimination was assessed via Area Under the Precision-Recall Curve (AUC- PR, 0-1 scale, with 0.5 indicating random performance), balancing precision and recall, making it particularly relevant for rare species, as it focuses on correctly predicting presences without being inflated by abundance of absences like AUC (McDermott *et al*., 2025 ; Sofaer *et al*., 2019). Classification ability was estimated through Sensitivity, measuring true positive rates (0 for no true positives, 1 for perfect classification), crucial given limited occurrences, where any omission of known occurrences would disproportionately affect model reliability. Calibration was quantified using Boyce Index, assesses alignment between predictions and observed presences (-1 for counter-prediction, 0 for random, and 1 for optimal prediction) (Liu *et al*., 2025 ; Sillero *et al*., 2021). This metric relies solely on presence data, making it particularly advantageous in presence-pseudoabsence frameworks where absence information is artificially generated.

For each pseudo-absence set, block cross-validation was used to identify the best-performing parameter set, and the corresponding model was then run and its evaluation retained for subsequent analyses. A four-fold (k=4) spatial cross-validation was used, training models on k-1 blocks and testing on the remaining one. Spatial block size was derived from the range of environmental spatial autocorrelation, estimated using the cv_spatial_autocor() function from the blockCV package (Valavi *et al*., 2019), based on 65,496 randomly sampled cells (subsampled by the function from the 1,244,424 cells available in the study area). Given limited occurrence data, final block size was set to two thirds of the median autocorrelation range. This configuration of k and block size balances environmental independence and data sufficiency that ensure presence records in each fold. Larger blocks improve spatial independence but constrain k to avoid empty folds; conversely, higher k increases replication but may reduce independence. A script is available (Appendix C) to explore multiple k and block size combinations, aiming to retain as many species as possible. Once the optimal parameters were selected per species and algorithm, models were refitted, and only those with Boyce Index > 0.5 were retained for ensemble modeling. This threshold ensures minimal calibration quality.

Ensemble modeling enhances SDMs robustness by combining multiple model outputs, effectively improving predictive accuracy by better capturing species distribution patterns (Marmion *et al*., 2009 ; Zhu & Peterson, 2017). However, ensemble models do not always outperform individual ones in transferability (Zhu *et al*., 2021; Zhu & Peterson, 2017). For this reason, ensemble modeling was applied at two distinct stages: (i) for current conditions, based on model runs calibrated and validated (Boyce Index > 0.5, maximum 5 algorithms x 10 pseudo absence set = 50 models) under present-day climate; and (ii) for future projections, by projecting these same validated models to every combination of future periods and scenarios, and then creating an ensemble for each period-scenario combination. Consensus ensemble maps were computed as pixel-wise median across all algorithms and replicates, minimizing the influence of outliers and aligning with model central tendencies (Marmion *et al*., 2009 ; Zhu *et al*., 2021). The entire pipeline is independently applied to each species, resulting in 17 continuous distribution maps per species (1 under current conditions, 16 under future). Ensemble models assessments were calculated by averaging metrics from models used to build the ensemble (Boyce Index > 0.5). Algorithm performances were compared using Kruskal-Wallis, with Dunn’s post hoc test (Bonferroni correction) identifying pairwise differences between algorithms for each evaluation metric.

#### Assessing uncertainty and spatio-temporal dynamics

Uncertainty was quantified using standard deviation across predictions. Pixels with >5 % divergence, indicating inconsistent predictions across algorithms, were masked for visualization on the current distribution map to highlight most reliable regions. To assess SDMs reliability under novel conditions (i.e. conditions in future periods deviating from the calibration range), the Multivariate Environmental Similarity Surface (MESS) index (Elith *et al*., 2010) was applied. Negative MESS values, indicating extrapolation, were masked for visualization on future distribution maps to highlight most reliable projections.

To quantify extent changes in suitable bioclimatic areas, maps were binarized for each period and SSP using the MaxSSS threshold, appropriate for presence-only data (Liu *et al*., 2016, 2013). Spatial overlap between current and future suitable areas was also computed on binarized maps, yielding the proportion of the present distribution projected to persist under each SSP. Spatial shifts between the current period and 2081-2100 were assessed by overlaying binarized projections with elevation data (CGIAR-CSI, 2018, ∼1x1 km at the equator) and latitude. Mean latitude and elevation of suitable pixels were calculated for the current period and for each SSP in 2081-2100 period. Values were averaged across the four scenarios, and only species retaining suitable areas in all SSPs were used to estimate shifts between the present and future. Continuous suitability maps for the current and 2081-2100 periods and all SSPs were averaged across species, with persistent suitable pixels identified as bioclimatic hotspots (areas where the highest number of species could find suitable bioclimatic conditions). Results are measured on a continuous scale to emphasize relative hotspot importance.

Framework and model characteristics were documented following the ODMAP protocol (Zurell *et al*., 2020, Appendix C) and summarized in Figure 3.

**Figure 3:**
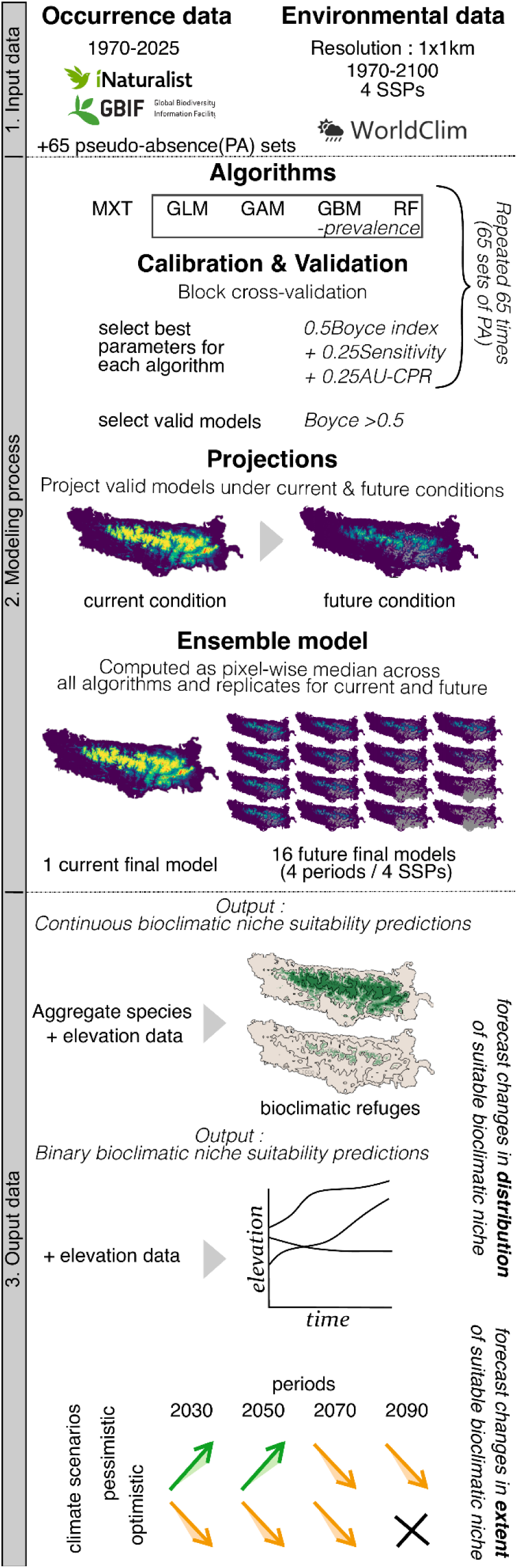
Overview of the species distribution modeling pipeline. The framework consists of three main stages: (**1**) Input data, including occurrence records, environmental data processing, and background sampling; (**2**) Modeling, encompassing model building, performance assessment, projection, and ensemble modeling; (**3**) Output processing, manipulating species projections to assess spatiotemporal dynamics of suitable bioclimatic niches.

## 3 Results

### Models performances

Between 19 and 50 models per species were retained for ensemble modeling based on the Boyce Index criterion (Boyce Index > 0.5). Significant differences in model performance across the five algorithms were detected for the Boyce Index and Sensitivity (Kruskal–Wallis, p < 0.001 and 0.01 respectively; see Figure 4a for p-values), while no significant difference was found for AUC-PR (p = 0.07). Dunn’s post hoc test revealed significant pairwise differences between algorithms. For the Boyce Index, MXT and GBM outperformed GAM, GLM and RF. Regarding Sensitivity, MXT outperformed GLM and GAM, while GBM and RF outperformed GLM. Other comparisons were not significant (Figure 4a). This highlights variation in predictive performance, with GBM and MXT generally performing best, and RF consistently performed well in identifying suitable areas where species are present. GLM exhibited the lowest Boyce Index values, falling under 0.5 for multiple species, indicating weaker prediction consistency.

**Figure 4:**
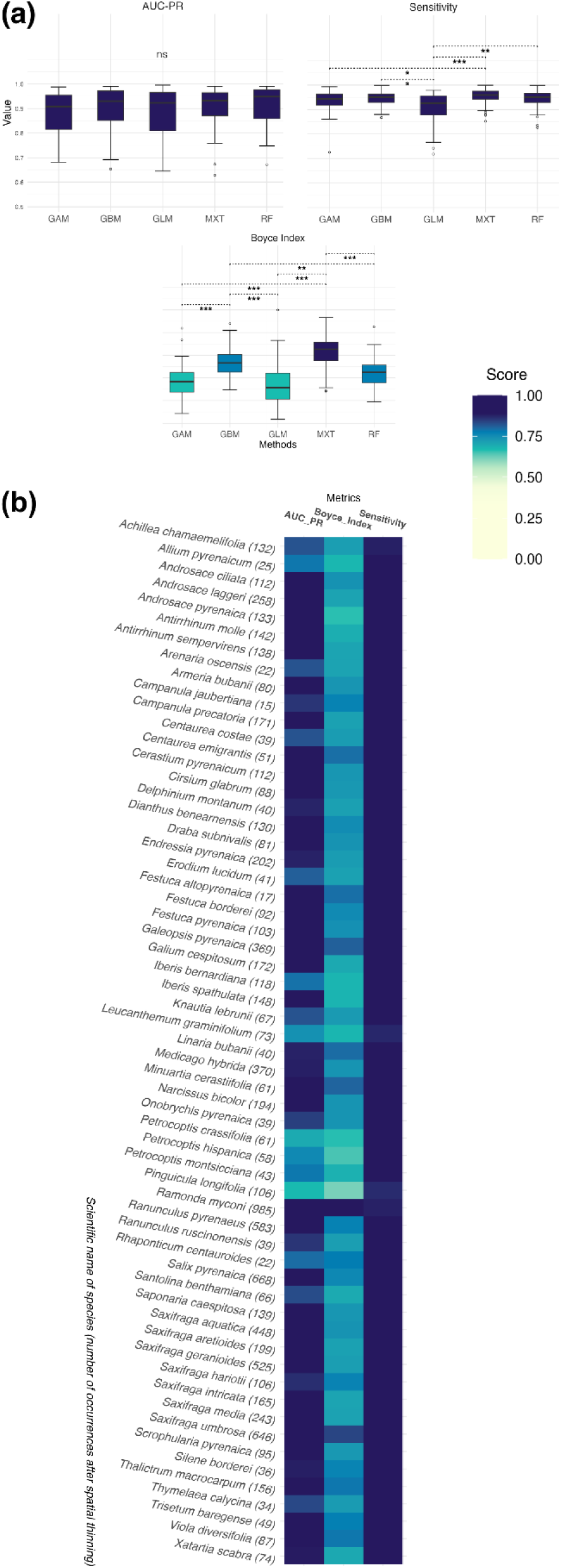
Endemic Pyrenean plant species model performances. **(a)** Boxplot of three performance metrics, AUC-PR (discrimination), Boyce Index (calibration), and Sensitivity (classification), for the five modeling algorithms (GAM, GBM, GLM, MAXENT, RF) for all species. Statistical significance is denoted as: *\*p* < 0.05*, **p* < 0.01*, ***p* < 0.001. **(b)** Performance of ensemble models across species, computed by averaging evaluation metrics from the validated models (Boyce Index > 0.5) across the five algorithms used to build the ensemble consensus. Darker shades indicate higher performance.

Ensemble model performance ranged from 0.67-0.98 for AUC-PR, 0.62-0.92 for the Boyce index and 0.87-0.98 for sensitivity (Figure 4b). Poor evaluation metrics were not consistently linked to species with fewer presence points, suggesting additional factors influencing performance.

### Projected bioclimatic niche loss and species vulnerability

The spatial extent of bioclimatic suitable areas for endemic Pyrenean plant species is projected to decline under climate change (Figure 5). By the end of the century, on average across SSPs, 57 out of the 59 species are projected to lose a significant part of their suitable bioclimatic niche, with 41 species losing >75 %, 11 losing 50-75 % and 7 experiencing a more moderate loss of <50 %. These 57 species are expected to lose on average between -62.35 % (min: +31.9 %, max: -100 %) and -95.21 % (min: -19.8 %, max: -100 %) of their suitable bioclimatic niche by 2081-2100 period under the most optimistic (SSP126) and the worst-case (SSP585) climate scenario, respectively. The severity of loss varies across species. More than half (32 species) are projected to experience a complete loss (100 %) of their suitable bioclimatic areas by the end of the century for at least one of the SSP, putting them at elevated risk of extinction. *Delphinium montanum*, *Endressia pyrenaica, Silene borderei* and *Onobrychis pyrenaica* appear particularly vulnerable, as models project a total disappearance of their bioclimatic suitable areas as early as the 2021-2040 period, under all SSPs, suggesting that these species are already present in unsuitable bioclimatic conditions. In contrast, *Pinguicula longifolia* and *Linaria Bubanii* emerged as exceptions, showing consistent increase in suitable areas with an average gain of +127.4 % (min: +95.5%, max: +181.63%) by 2081-2100 period. This suggests that shifting climatic conditions may benefit some species, including endemic and threatened ones. The analysis also highlights an accelerating trend in suitable area loss over time, particularly in more extreme scenarios such as SSP370 and SSP585. Most species experience sharp declines after the 2041-2060 period, reflecting the escalating impact of climate change as the century progresses (Figure 5). All suitability maps, covering both current and future projections for all species, are available in Appendix D.

**Figure 5:**
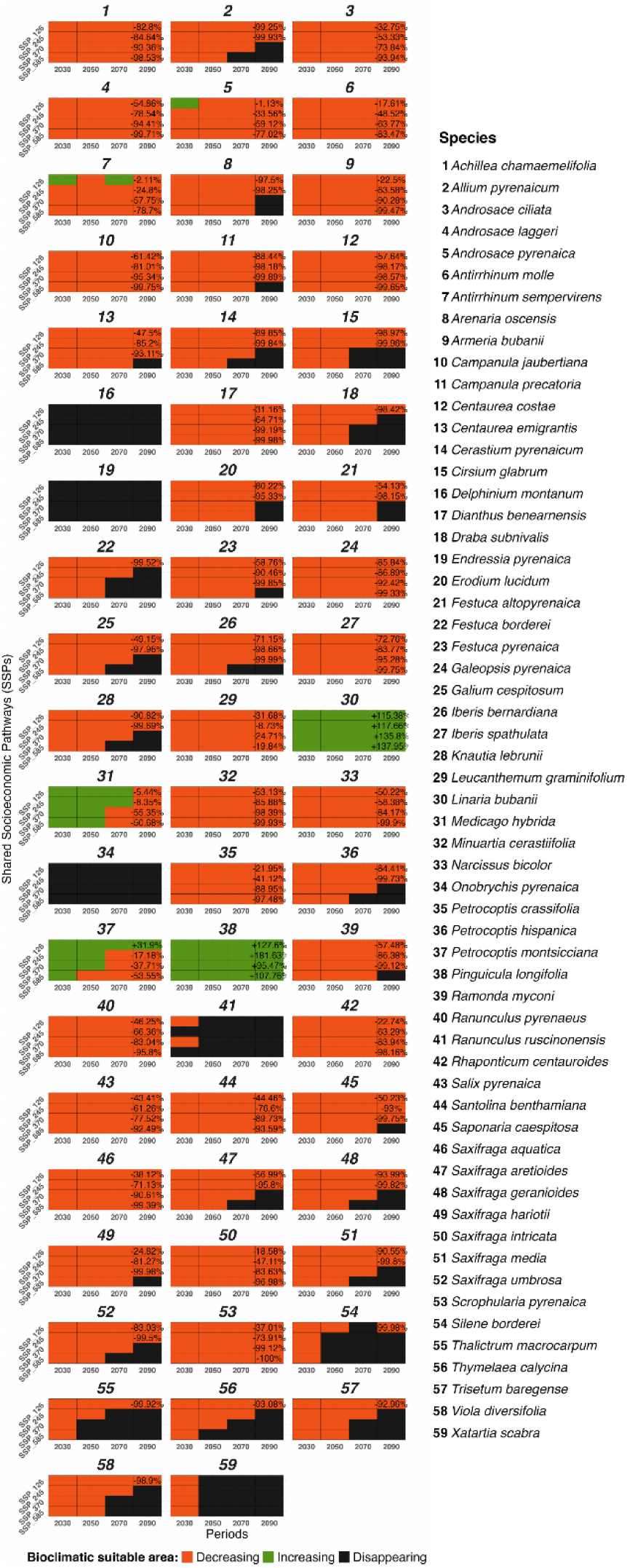
Projected extent changes in bioclimatic suitability for endemic Pyrenean plant species under Shared Socioeconomic Pathways (SSP) for the 2021-2100 period. Extent changes in suitable areas under SSP126, SSP245, SSP370, SSP585, relative to the baseline period (1970-2000). Each box represents the extent change in suitable area, with percentage values shown for the 2081-2100 period. Orange indicates a reduction in suitable areas, green expansion, and black a complete loss of suitability. 2021-2040, 2041-2060, 2061-2080, 2081-2100 periods designated as 2030, 2050, 2070, 2090.

### Spatial dynamics in bioclimatic niche suitability and upslope shift of bioclimatic hotspots

The spatial distribution of bioclimatic niche suitability for endemic Pyrenean plant species is projected to undergo significant changes by the 2081-2100 period under climate change (Figure 6). Currently, bioclimatic hotspots, where most species could find suitable bioclimatic conditions, are mainly above 1000 m, with a maximum favorability of 0.73 (Figure 6a, top panel). By the 2081-2100 period, these hotspots are projected to shrink in size and shift upward, with the extent of this change strongly linked to the emission scenario. Climate scenarios such as SSP126 show a relatively moderate reduction, with some areas persisting near current locations. In contrast, more extreme scenarios projected a pronounced contraction of suitable areas, becoming increasingly fragmented. Under SSP585, the maximum favorability value drops to 0.29, with suitable conditions restricted to areas above 2000 m in elevation (Figure 6a, lower panels).

**Figure 6.**
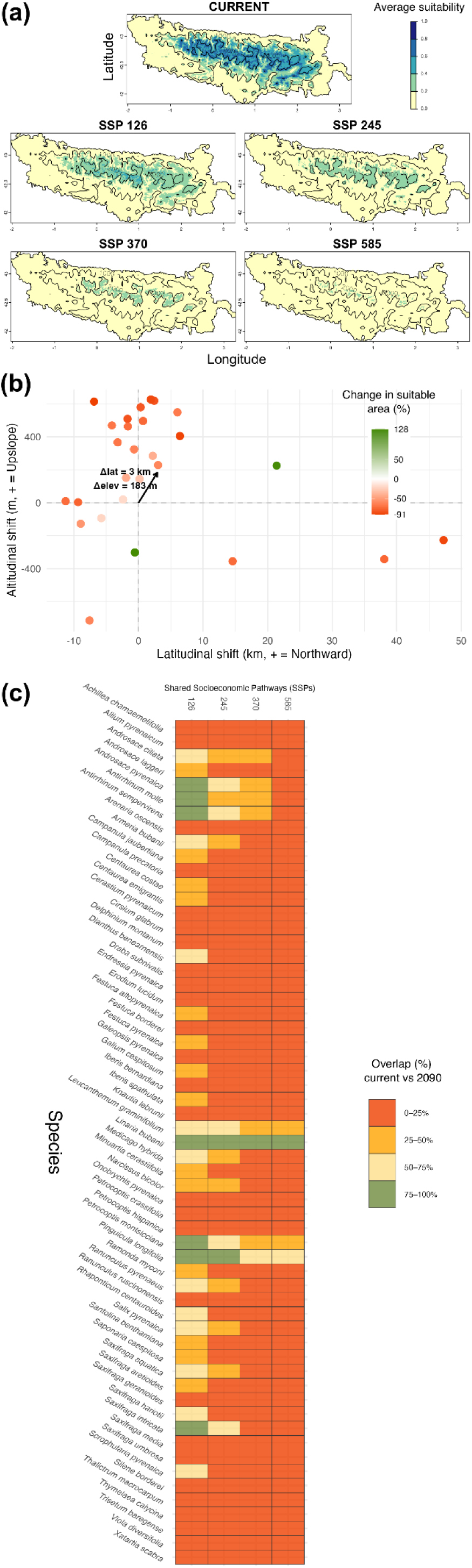
Projected distribution changes in bioclimatic suitability for endemic Pyrenean plant species between current and 2081-2100 periods. **(a)** Current and projected (2081-2100) bioclimatic hotspots, i.e. areas where the highest number of species could find suitable bioclimatic conditions, under Shared Socioeconomic Pathways (SSPs). The top panel shows the current distribution, while lower panels display projections for SSP126, SSP245, SSP370, and SSP585. Darker colors indicate higher bioclimatic suitability. Contour lines mark the 1000- and 2000-meter elevation topography. **(b)** Projected latitudinal and elevational shifts in bioclimatic suitability for the 2081– 2100 period, averaged across SSPs. Each dot represents one of the 27 species retaining suitable areas under all SSPs. Dot color reflects the relative change in suitable area (green = gain; red = loss). The black arrow indicates the overall mean displacement in latitude (Δ lat) and elevation (Δ elev) across species. **(c)** Projected spatial overlap (%) between current and future (2081-2100) distributions of bioclimatically suitable areas under SSP126, SSP245, SSP370, and SSP585. Each row represents one species, with colors indicating the proportion (%) of overlap with present-day suitable areas.

The projected mean elevation of bioclimatic suitable areas for endemic Pyrenean plant species shows a marked upward shift by the end of the century under climate change (Figure 6b). Currently, mean elevation of suitable bioclimatic area ranges between 748 and 2269 m, depending on the species. By 2081-2100, the average elevation across species is projected to increase by +182.58 m. While the lower margin remains relatively stable (671 m), the upper margin is expected to reach 2779 m. Species exhibit variable projected elevation trajectories. By the 2081-2100 period, 20 of the 27 species that still have suitable bioclimatic conditions across all SSPs display a clear upward projected trend, with mean increases ranging from +0.26 % to +38.62 %. Subsets of species that may experience a reduction in their mean elevation could see decreases from -8.93% to -41.51 %. On average, species show a +19.49 % increase in elevation, while the decrease is about -23.44 %. In addition to these vertical shifts, species also display heterogeneous latitudinal responses by 2081-2100 period. Among the 27 species included, 13 are projected to shift northward (+0.2 to +47.3 km, mean +11.1 km), while 14 shift southward (-11.3 to -0.6 km, mean -4.7 km), resulting in an overall average shift of +2.91 km northward. These results emphasize the significant influence of warming climate on the spatial distribution of suitable bioclimatic areas, underscoring the increasing vulnerability of endemic Pyrenean flora to rising temperatures.

By 2081–2100, overlap between current and future suitable bioclimatic areas is generally low. Out of 236 species x scenario combinations, 177 (75%) fall in the lowest overlap class (0-25%). At the species level, 55 of the 59 endemics experience at least one scenario with less than 25% overlap. Notably, 38 species show complete loss of overlap (0%) under the most severe scenario (SSP585). This means that while 32 species entirely disappear from the projections, 6 still retain suitable climatic conditions, but in areas completely distinct from their current distribution. In contrast, only 7 species retain relatively high stability (≥75% overlap) under at least one scenario. These results highlight the drastic spatial reshuffling of suitable areas and the limited persistence of current bioclimatic ranges under future climates.

## 4 Discussion

### Trends in bioclimatic niche suitability under climate change

This study paints a concerning picture of the future for Pyrenean endemic flora under ongoing and projected climate change. By the 2081-2100 period, 57/59 of the studied species are projected to experience substantial reduction in suitable bioclimatic areas (on average -62.35 % to -95.21 % for SSP126 and SSP585 respectively), ranging from moderate contraction in some species to complete losses in others. These findings underline the severe threat posed by changing climate to narrowly distributed species and align with patterns projected in similar mountain regions, such as the Alps (Dagnino *et al*., 2020 ; DirnbÖck *et al*., 2011 ; Dullinger *et al*., 2012 ; HÜlber *et al*., 2016 ; Rota *et al*., 2022), where endemics are acutely sensitive to rapid warming. Parallels with other mountain systems (Manes *et al*., 2021) underscore the universal challenge that climate change poses to mountain endemism which is, moreover, amplified by topography and human pressures (Elsen et al., 2020). Interestingly, not all species follow the same trajectory. While most of them show severe bioclimatic niche contractions, *Pinguicula longifolia* and *Linaria bubanii* are the only endemics projected to consistently gain suitable bioclimatic areas under all scenarios, mainly on the northern slopes and in the southeastern Pyrenees. For *P. longifolia*, this may reflect preadaptation to warmer climates, as suggested by the occurrence of a related subspecies in the drier Massif Central. For *L. bubanii*, a scree- dwelling species of high altitudes, the strong microclimatic heterogeneity of its habitat may facilitate persistence across diverse conditions and promote expansion. Despite these projected gains, both species may remain vulnerable as establishment depends on strict ecological requirements and interactions with other biotic components under climate change. These outliers highlight that while some species face heightened risks of extinction, others may find new opportunities under climate change. It also underlines the need for species-specific conservation strategies that account for this ecological variability.

Beyond species-specific patterns, this study revealed notable upward shifts in mean elevations of suitable bioclimatic niches, exceeding 180 m by the 2081-2100 period. If such a trend had been consistent over the past century, it could help explain a part of the upward plant movement already observed in the Pyrenees (Ameztegui *et al*., 2016 ; AullÓ-Maestro *et al*., 2023 ; Delpouve *et al*., 2025 ; Lenoir *et al*., 2008 ; Marshall *et al*., 2020 ; Pauli *et al*., 2012) and is consistent with documented upward trends in other various mountain ecosystems (Chan *et al*., 2024 ; Iseli *et al*., 2023). In parallel for the same period, latitudinal dynamics indicate an overall northward displacement of about +3 km. Overlap with current ranges is also very limited: 55 of the 59 endemic species retain less than 25% of their present distribution under at least one scenario, implying that many future suitable areas will occur entirely outside their current ranges. Combined with the projected upward shift, it suggests a profound spatial displacement of endemic species under ongoing warming. However, the pace at which species can track these changes remains uncertain, as local-scale factors, such as microclimatic buffering, habitat discontinuities, or dispersal limitations, may slow or constrain their responses (Maclean & Early, 2023). Hotspots of suitable bioclimatic niches, currently located above 1000 m, are projected to shift above 2000 m. This shift drastically reduces bioclimatic suitable space available for endemic flora, as high elevations cover smaller geographic extent in the Pyrenees (Elsen & Tingley, 2015). Many species risk being squeezed between warming lower slopes and shrinking refugia, exacerbating their risk of extinction. Bioclimatic hotspot maps revealed spatial patterns suggesting a future fragmentation of suitable areas, with favorable habitats increasingly confined to isolated mountaintops separated by valleys. This fragmentation likely reduces connectivity across the landscape, limiting the ability of many species to track climate change. In the Pyrenees, this issue is compounded by regional warming and aridification, already documented at a rate of 30 % faster than the global average over the past 50 years (OPCC, 2018). Combined with challenges already posed by dispersal barriers, topographic constraints, and competition from upslope-moving species, these factors intensify pressure on the native flora. The severe suitable bioclimatic area reductions projected under SSP370 and SSP585 scenarios after the 2041-2060 period is particularly alarming and suggests that environmental pressures will intensify in the second half of the century. The urgency of monitoring is reinforced by the identification of high vulnerability of northern Spanish flora, to the south of the Pyrenees, which is among the most threatened in Europe (Engler *et al*., 2011).

### Strengths and limitations of the modeling framework

The pipeline developed in this study represents a step forward in addressing mountain biodiversity questions at a broad geographic scale, particularly for analyzing climate-driven trends across multiple species. By integrating five algorithms (GLM, GAM, GBM, RF, and MXT) into an ensemble modeling framework, this approach minimizes model-specific biases and improves the reliability of projected species distributions. Moreover, the agreement with previous projections for an endemic species also analyzed here (e.g. *D. montanum*, Salvado *et al*., 2022) suggests that our ensemble approach produces ecologically consistent outcomes. RF, MXT and GBM emerged as particularly strong performers, likely due to their capacity to handle small sample sizes and capture non-linear relationships between species and environmental variables (Valavi *et al*., 2022). The pipeline addresses challenges posed by sparse data, providing consistent and interpretable results for narrowly distributed species, as demonstrated by its ability to model 75% of the endemic species in a mountainous region. It includes tools to adjust pseudo-absence sampling to obtain stable predictions despite limited occurrence data. Another component helps optimize spatial cross-validation settings ensuring robust as possible model evaluation while maximizing the number of species that can be assessed and thus studied. These features make the approach suitable for rare or endemic species, whose restricted ranges and small sample sizes could limit the applicability of standard SDMs procedures. To account for endemic Pyrenean plant species excluded from this study, the pipeline could be applied case-by-case at a smaller spatial scale (in regions of recorded occurrences). As such, it provides a valuable and adaptable framework for informing conservation strategies across a diversity of endemic taxa. Moving forward, our pipeline could be extended to species with broader distributions, adapted to biodiversity hotspots worldwide, and optimized for a more extensive analysis across multiple species. It would provide deeper insights into species and communities’ response to climate change, focusing on richness patterns and diversity dynamics.

Despite these strengths, several SDMs limitations and considerations warrant caution when interpreting the results (Collette et al., in press), particularly regarding methodological biases and decision-making processes (see Sillero *et al*., 2021; Bryn *et al*., 2021; Leroy, 2023). This study presents three main considerations. First, in large-scale studies where fine-resolution data are lacking, coarse-resolution bioclimatic data can still capture general environmental trends, reduce computational costs, and minimize biases from inaccuracies in occurrence data. However, by failing to capture fine-scale microhabitat heterogeneity in mountainous environments, they may misestimate the estimated extent of habitat suitability (Franklin *et al*., 2013 ; Gottfried *et al*., 2012 ; Nadeau *et al*., 2017 ; Randin *et al*., 2009 ; Trivedi *et al*., 2008), although the overall trends remain consistent (Tsiftsis *et al*., 2024). Second, projections beyond 2041-2060 period remain uncertain due to increasing divergence between climate models, particularly for worst climate scenarios (SSP370, SSP585) (Goberville *et al*., 2015 ; Tebaldi *et al*., 2021). Although the trajectory for most species aligns across scenarios, notable variation exists in whether suitable bioclimatic niche disappears or persists to a lesser extent, depending on pseudo-absence sets. Long-term monitoring and high-resolution climate data are essential to validate projections and improve predictive tools. Third, the exclusion of biotic interactions (e.g. competition, mutualism, pollination), and important abiotic variables (e.g. soil, hydrology, radiation, land use) reduces the ecological realism of projections. Integrating such dynamics into SDMs would enhance the accuracy of projections and offer a more nuanced understanding of species’ responses to climate change (Cosentino *et al*., 2023 ; Lee-Yaw *et al*., 2022 ; Preuss & Padial, 2021).

Beyond these methodological considerations, SDMs also fail to capture complex ecological and evolutionary processes shaping species’ responses to environmental changes (Williams *et al*., 2008). Expansion of bioclimatic suitability areas does not guarantee population growth, as dispersal barriers, establishment limitations, and biotic interactions may restrict species from fully occupying suitable areas. Likewise, suitability contraction does not imply imminent range collapse, as species can persist temporarily in suboptimal conditions before extinction, known as “extinction debt” (Cotto *et al*., 2017 ; Nomoto & Alexander, 2021).

On the adaptation perspective, rapid environmental changes may outpace the rate at which species can adapt, facing the disappearance of their suitable habitat (Cotto *et al*., 2017 ; Jump & PeÑuelas, 2005 ; Shaw & Etterson, 2012). Moreover, fragmented populations may experience divergent selective pressures, fostering local adaptation, often overlooked in SDMs. Integrating local adaptation, phenotypic plasticity (Benito-Garzon *et al*., 2019 ; Valladares *et al*., 2014), and genomic data (HÄllfors *et al*., 2016 ; Kort *et al*., 2024) could improve projections by better accounting for populations’ adaptive capacity under environmental stress. Despite these limitations, correlative SDMs based on bioclimatic variables remain valuable tools for generating insights and understanding species’ ecological niches, anticipating distributional changes, and informing conservation strategies.

### Conservation implications and Policy recommendation

From conservation and management perspectives, our findings underscore the urgent need for conservation strategies that address specific challenges faced by mountain flora in the Pyrenees. High altitude environments, where suitable bioclimatic conditions are expected to persist, represent critical bulwarks against climate change. These areas must be prioritized for legal protection as critical refugia for biodiversity preservation, since they are projected to concentrate the remaining favorable conditions for species by 2081- 2100. To ensure the long-term relevance of protected areas, conservation planning must integrate future suitability and regulate potentially harmful activities in these refugia, such as infrastructure development or unregulated tourism, particularly above 2000 m. However, traditional grazing should be promoted, as it maintains open habitats that support many endemic species adapted to these environments. In parallel, the consistent upward shift of bioclimatic hotspots and their fragmentation call for the maintenance of altitudinal corridors to support species migration and gene flow, thereby mitigating the adverse effects of isolation. Mapping these corridors based on projected trajectories could help identify priority areas for land acquisition or land-use planning. For species projected to experience a partial lose their suitable bioclimatic niche, conservation interventions such as assisted migration and translocation may provide options to prevent extinction (Ferrarini *et al*., 2016 ; Gallagher *et al*., 2014). Evaluating candidate sites for reintroduction in newly suitable areas would facilitate the implementation of these actions. Moreover, integrating predictive modeling tools with field monitoring could enhance the precision of these interventions, ensuring that resources are allocated to areas with the greatest potential for success (Elliott *et al*., 2024 ; Eyre *et al*., 2022). Critically vulnerable taxa such as *Delphinium montanum*, *Silene borderei*, and *Onobrychis pyrenaica*, for which models predict the disappearance of suitable conditions as early as 2021-2040 under all SSPs, should be urgently targeted. For these species, in situ strategies may need to be complemented by ex situ measures, such as seed banking or cultivation in botanical gardens, which represent last-resort solution to preserve genetic material and avoid extinction. Developing emergency conservation plans for such high-risk taxa would improve coordination across institutions and ensure readiness for rapid intervention. In parallel, identifying microrefugia through fine-scale topographic analysis may uncover overlooked zones of persistence and bring new areas for assisted migration. Similarly, intensified demographic and phenological monitoring within residual populations would help assess whether remnant individuals are viable or demographically declining, thereby informing prioritization. Re-evaluating conservation status in light of projected niche loss would better guide adaptive management priorities. Many endemic species are likely to face imminent extinction risks that are not yet recognized in current conservation assessments. Including climate-driven projections in red list criteria could improve the alignment between legal protection and future vulnerability. National and regional conservation strategies should be updated to incorporate forward-looking criteria, ensuring that legal frameworks remain dynamic in the face of environmental change.

The transboundary distribution of Pyrenean species across France, Spain, and Andorra requires coordinated cross-border efforts to avoid fragmented conservation initiatives, such as inconsistent Natura 2000 habitat interpretation along this frontier (Prud’homme & Ninot, 2018). Adaptive legal frameworks are essential to address these shifts, supported by collaborative efforts that align research, policy, and community engagement for enhanced conservation outcomes. Establishing a transnational conservation platform for endemic mountain species could help harmonize monitoring, protection, and restoration efforts across borders, and reduce disparities in species management. Translating scientific evidence into clear and actionable policy briefs can bridge the gap between research and implementation, fostering informed and, consequently, more effective decision-making (Guisan *et al*., 2013). Understanding how climate change affects extinction risks and habitat availability is key to prioritizing conservation actions, especially as pressures intensify. This also requires combining ecological modeling, field experimentation, and long-term monitoring to refine conservation strategies and better address species redistribution and their effects on human well-being (Bonebrake *et al*., 2018 ; Pecl *et al*., 2017). Ultimately, preserving mountain biodiversity in an era of rapid change will require collaboration among scientists, resource managers, and local communities, supported by predictive tools to capture the complex realities of a rapidly changing world.

## Supporting information

Supplemental Information Bioclimatic niche shifts in Pyrenean species

## Funding statement

This study received financial support from the Occitanie Region (Emergence 2024) and the Commissariat du Massif des Pyrénées (FNADT 2023), as well as from the Fédération de Recherche 2043, Energie & Environnement (UPVD/CNRS). It also received technical and financial support from the Interreg- POCTEFA program as part of the Floralab+ project (EFA024/01).

## Acknowledgements

We thank Romain BERTRAND (Toulouse III University) for valuable comments that contributed to shaping our methodology.

## Author Contributions

**Conceptualization**, J.A.M.B., N.C., V.H. ; **Data curation**, N.C. ; **Formal analysis**, N.C. ; **Funding Acquisition**, J.A.M.B., S.P., V.H. ; **Investigation**, N.C. ; **Methodology**, J.A.M.B., N.C., S.P. V.H.; **Project administration**, J.A.M.B., S.P., V.H. ; **Software**, N.C. ; **Supervision**, J.A.M.B., S.P., V.H ; **Validation**, J.A.M.B., N.C., S.P., V.H. ; **Visualization**, J.A.M.B., N.C., S.P., V.H. ; **Writing – original draft**, J.A.M.B., N.C., S.P., V.H. ; **Writing – review & editing**, J.A.M.B., N.C., S.P., V.H..

## Data and code availability statement

The original data supporting the findings of this study are openly available from public domain resources, including GBIF, WorldClim, and iNaturalist. The code and data required to reproduce the results are accessible in the Zenodo repository at [DOI : <u>10.5281/zenodo.17412264</u>] and DRYAD Further supporting details are available in the Supplementary Information.

## Conflict of interest statement

The authors declare no conflicts of interest.

## Notes

https://doi.org/10.5281/zenodo.17412264

### Competing Interest Statement

The authors have declared no competing interest.

### Summary of Updates

This version follows revision conducted after peer review in the journal Ecography, where the article is now accepted (DOI: 10.1002/ecog.08067). The manuscript has been thoroughly updated, including text, figures, and methods. Importantly, despite these extensive revisions and re-runs of the analyses, the main results and interpretations remain consistent with the previous version. This stability confirms that the original methodological framework was already robust, and that the new modifications primarily improved clarity, transparency, and compliance with current best practices in ecological modeling rather than changing the scientific conclusions.

