## Supplemental Information Bioclimatic niche shifts in Pyrenean species for "Projecting spatiotemporal bioclimatic niche dynamics of endemic Pyrenean plant species under climate change: how much will we lose?"

**Table of Contents:**


| | |
|---|---|
| **Appendix A: Definitions and selection of WorldClim 2.1 bioclimatic variables** | **Page 1** |
| **Appendix B: Comparison of pseudo-absence configurations across algorithms** | **Page 5** |
| **Appendix C: Detailed overview of SDMs framework according to the ODMAP protocol** | **Page 6** |
| **Appendix E: Current and future bioclimatic niche suitability continuous maps for 59 endemic Pyrenean species** | **Page 16** |
| **References for Appendices** | **Page 77** |

**Appendix A: Definitions and selection of WorldClim 2.1 bioclimatic variables**

**Table A1: Definitions of WorldClim 2.1 bioclimatic variables.** The WorldClim 2.1 public database (Fick & Hijmans, 2017) provides 19 bioclimatic variables in a raster layer format, including 11 related to temperature and 8 to precipitation. These variables are derived from interpolated average monthly climate data collected from weather stations.

| Variable | Name | Description | Unit |
|---|---|---|---|
| BIO1 | Mean annual temperature | Average annual temperature | °C |
| BIO2 | Mean diurnal range | Mean of monthly temperature ranges (Tmax - Tmin) | °C |
| BIO3 | Isothermality | (BIO2 / BIO7) (*100): Proportion of diurnal variation relative to annual variation | % |
| BIO4 | Temperature seasonality | Temperature variability (standard deviation * 100) | °C |
| BIO5 | Max temperature of warmest month | Maximum temperature of the warmest month | °C |
| BIO6 | Min temperature of coldest month | Minimum temperature of the coldest month | °C |
| BIO7 | Temperature annual range | Annual temperature range (BIO5 - BIO6) | °C |
| BIO8 | Mean temperature of wettest quarter | Mean temperature of the wettest quarter | °C |
| BIO9 | Mean temperature of driest quarter | Mean temperature of the driest quarter | °C |
| BIO10 | Mean temperature of warmest quarter | Mean temperature of the warmest quarter | °C |
| BIO11 | Mean temperature of coldest quarter | Mean temperature of the coldest quarter | °C |
| BIO12 | Annual precipitation | Total annual precipitation | mm |
| BIO13 | Precipitation of wettest month | Precipitation in the wettest month | mm |
| BIO14 | Precipitation of driest month | Precipitation in the driest month | mm |
| BIO15 | Precipitation seasonality | Precipitation variability (coefficient of variation) | % |
| BIO16 | Precipitation of wettest quarter | Total precipitation in the wettest quarter | mm |
| BIO17 | Precipitation of driest quarter | Total precipitation in the driest quarter | mm |
| BIO18 | Precipitation of warmest quarter | Total precipitation in the warmest quarter | mm |
| BIO19 | Precipitation of coldest quarter | Total precipitation in the coldest quarter | mm |

**Selection of environmental variables:**

To reduce redundancy in models, correlated variables (Pearson r > 0.8) at presence locations for each species were grouped into clusters. Strongly negatively correlated variables (r < -0.8) can capture complementary environmental gradients (e.g. warm summers vs. cold winters) and were therefore kept in the analysis. From each cluster, the highest-ranked variable was retained based on a predefined ecological priority list : BIO1 & BIO12 > BIO5 (Román-Palacios & Wiens, 2020) > BIO6 > BIO7 > BIO15 > BIO10 > BIO18 > BIO11 > BIO13 > BIO14 > BIO16 > BIO17 > BIO4 > BIO8 > BIO9 > BIO19 > BIO3 > BIO2.

**Figure 1A: Selected bioclimatic variables in species distribution models.** Number of species (out of 59) for which each bioclimatic variable was retained in the final models after variable selection.

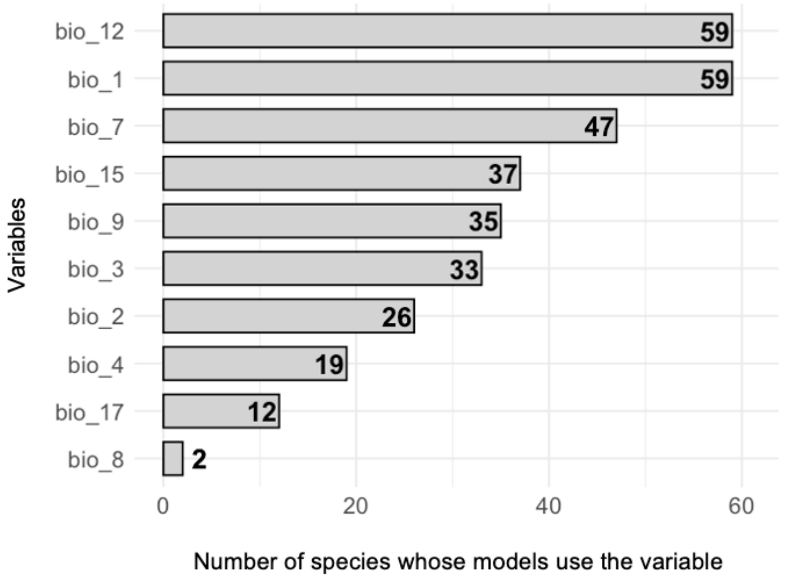


**Table 2A: Species and their selected bioclimatic variables for modeling.** For each species, the subset of bioclimatic variables retained after correlation filtering and variable selection.

| scientific_name | variables | scientific_name | variables |
|---|---|---|---|
| *Achillea chamaemelifolia* | bio_3, bio_9, bio_1, bio_12 | *Linaria bubanii* | bio_2, bio_3, bio_9, bio_1, bio_12 |
| *Allium pyrenaicum* | bio_4, bio_7, bio_8, bio_9, bio_15, bio_1, bio_12 | *Medicago hybrida* | bio_4, bio_8, bio_9, bio_15, bio_1, bio_3, bio_12 |
| *Androsace ciliata* | bio_4, bio_7, bio_1, bio_12 | *Minuartia cerastiifolia* | bio_2, bio_1, bio_7, bio_12 |
| *Androsace laggeri* | bio_4, bio_7, bio_9, bio_15, bio_1, bio_3, bio_12 | *Narcissus bicolor* | bio_3, bio_9, bio_15, bio_1, bio_7, bio_12 |
| *Androsace pyrenaica* | bio_2, bio_1, bio_7, bio_12 | *Onobrychis pyrenaica* | bio_3, bio_15, bio_1, bio_4, bio_12 |
| *Antirrhinum molle* | bio_3, bio_9, bio_1, bio_4, bio_12 | *Petrocoptis crassifolia* | bio_4, bio_7, bio_15, bio_1, bio_3, bio_12 |
| *Antirrhinum sempervirens* | bio_2, bio_15, bio_1, bio_7, bio_12 | *Petrocoptis hispanica* | bio_3, bio_15, bio_1, bio_4, bio_12 |
| *Arenaria oscensis* | bio_3, bio_15, bio_1, bio_7, bio_12 | *Petrocoptis montsicciana* | bio_3, bio_9, bio_1, bio_4, bio_12 |
| *Armeria bubanii* | bio_1, bio_7, bio_12 | *Pinguicula longifolia* | bio_2, bio_3, bio_15, bio_1, bio_7, bio_12 |
| *Campanula jaubertiana* | bio_4, bio_7, bio_9, bio_15, bio_1, bio_3, bio_12 | *Ramonda myconi* | bio_3, bio_9, bio_15, bio_1, bio_7, bio_12 |
| *Campanula precatoria* | bio_2, bio_9, bio_1, bio_7, bio_12, bio_17 | *Ranunculus pyrenaeus* | bio_2, bio_4, bio_7, bio_9, bio_15, bio_1, bio_12, bio_17 |
| *Centaurea costae* | bio_3, bio_9, bio_15, bio_1, bio_7, bio_12 | *Ranunculus ruscinonensis* | bio_2, bio_9, bio_15, bio_1, bio_7, bio_12 |
| *Centaurea emigrantis* | bio_3, bio_1, bio_7, bio_12 | *Rhaponticum centauroides* | bio_2, bio_3, bio_15, bio_1, bio_12 |

| | | | |
|---|---|---|---|
| *Cerastium pyrenaicum* | bio_4, bio_7, bio_9, bio_1, bio_3, bio_12, bio_17 | *Salix pyrenaica* | bio_2, bio_9, bio_15, bio_1, bio_7, bio_12, bio_17 |
| *Cirsium glabrum* | bio_2, bio_3, bio_15, bio_1, bio_7, bio_12 | *Santolina benthamiana* | bio_3, bio_9, bio_15, bio_1, bio_4, bio_12 |
| *Delphinium montanum* | bio_7, bio_9, bio_1, bio_12 | *Saponaria caespitosa* | bio_3, bio_9, bio_15, bio_1, bio_7, bio_12 |
| *Dianthus benearnensis* | bio_3, bio_9, bio_15, bio_1, bio_7, bio_12 | *Saxifraga aquatica* | bio_4, bio_7, bio_9, bio_1, bio_3, bio_12, bio_17 |
| *Draba subnivalis* | bio_1, bio_7, bio_12 | *Saxifraga aretioides* | bio_2, bio_9, bio_15, bio_1, bio_7, bio_12, bio_17 |
| *Endressia pyrenaica* | bio_4, bio_7, bio_9, bio_1, bio_12 | *Saxifraga geranioides* | bio_2, bio_9, bio_1, bio_7, bio_12, bio_17 |
| *Erodium lucidum* | bio_3, bio_9, bio_15, bio_1, bio_7, bio_12 | *Saxifraga hariotii* | bio_15, bio_1, bio_7, bio_12 |
| *Festuca altopyrenaica* | bio_2, bio_3, bio_15, bio_1, bio_7, bio_12 | *Saxifraga intricata* | bio_2, bio_15, bio_1, bio_7, bio_12 |
| *Festuca borderei* | bio_4, bio_7, bio_9, bio_15, bio_1, bio_3, bio_12 | *Saxifraga media* | bio_2, bio_9, bio_1, bio_7, bio_12, bio_17 |
| *Festuca pyrenaica* | bio_2, bio_9, bio_15, bio_1, bio_7, bio_12 | *Saxifraga umbrosa* | bio_2, bio_9, bio_15, bio_1, bio_7, bio_12, bio_17 |
| *Galeopsis pyrenaica* | bio_2, bio_3, bio_9, bio_15, bio_1, bio_12 | *Scrophularia pyrenaica* | bio_2, bio_3, bio_9, bio_15, bio_1, bio_7, bio_12 |
| *Galium cespitosum* | bio_2, bio_15, bio_1, bio_7, bio_12 | *Silene borderei* | bio_4, bio_9, bio_1, bio_3, bio_12 |
| *Iberis bernardiana* | bio_2, bio_15, bio_1, bio_7, bio_12 | *Thalictrum macrocarpum* | bio_2, bio_15, bio_1, bio_7, bio_12 |
| *Iberis spathulata* | bio_4, bio_7, bio_9, bio_1, bio_3, bio_12 | *Thymelaea calycina* | bio_2, bio_15, bio_1, bio_7, bio_12, bio_17 |
| *Knautia lebrunii* | bio_2, bio_3, bio_1, bio_7, bio_12, bio_17 | *Trisetum baregense* | bio_2, bio_15, bio_1, bio_7, bio_12 |
| *Leucanthemum graminifolium* | bio_3, bio_9, bio_15, bio_1, bio_7, bio_12 | *Viola diversifolia* | bio_4, bio_1, bio_3, bio_12, bio_17 |
| | | *Xatartia scabra* | bio_7, bio_9, bio_1, bio_12 |

**Appendix B: Comparison of pseudo-absence configurations across algorithms**

Evaluation of alternative pseudo-absence configurations (recommended by (Barbet-Massin et al., 2012) and the Biomod2 team (Gueguen et al., 2025)) across all five algorithms and 59 species. Results indicate that in average, environmentally constrained pseudo-absences in equal number to presences consistently yielded the best evaluation scores across all algorithms, leading us to adopt this configuration for the final analyses.

**Table B: Cross-validation results for alternative pseudo-absence sampling strategies.** Tested configurations included environmentally constrained (*env_*) or random (*ran_*) sampling with pseudo-absences equal to the number of presences (*xn*), three times the number of presences (*x3n*), or fixed at 10,000 (*10k*). Each value corresponds to the mean across three runs (i.e., three distinct pseudo-absence sets) for the same parameter combination, thus providing triplicates for evaluation. The evaluation procedure was identical to that used in the main text, relying on three metrics: AUC-PR (discrimination), Boyce Index (calibration), and Sensitivity (classification). Despite slightly higher performance of GBM under random pseudo-absence sampling with the number of presences same as absence, we retained a single configuration across all algorithms to ensure comparability of models and performance.

| | GAM | | | GLM | | | GBM | | | RF | | | MXT | | |
|---|---|---|---|---|---|---|---|---|---|---|---|---|---|---|---|
| Configuration | AUC-PR | Boyce Index | Sensitivity | AUC-PR | Boyce Index | Sensitivity | AUC-PR | Boyce Index | Sensitivity | AUC-PR | Boyce Index | Sensitivity | AUC-PR | Boyce Index | Sensitivity |
| ran_xn | 0,812 | 0,616 | 0,919 | 0,821 | 0,740 | 0,916 | 0,813 | 0,518 | 0,895 | 0,821 | 0,772 | 0,937 | 0,817 | 0,650 | 0,918 |
| Algorithm metric average | 0,782 | | | 0,826 | | | **0,742** | | | 0,843 | | | 0,795 | | |
| ran_x3n | 0,651 | 0,644 | 0,926 | 0,649 | 0,713 | 0,924 | 0,672 | 0,540 | 0,885 | 0,670 | 0,748 | 0,935 | 0,641 | 0,648 | 0,908 |
| Algorithm metric average | 0,740 | | | 0,762 | | | 0,699 | | | 0,784 | | | 0,732 | | |
| ran_10k | /* | | | /* | | | /* | | | /* | | | /* | | |
| env_xn | 0,885 | 0,590 | 0,926 | 0,912 | 0,779 | 0,943 | 0,882 | 0,392 | 0,894 | 0,909 | 0,808 | 0,952 | 0,917 | 0,690 | 0,941 |
| Algorithm metric average | **0,800** | | | **0,878** | | | 0,723 | | | **0,890** | | | 0,849 | | |
| env_x3n | 0,793 | 0,586 | 0,945 | 0,832 | 0,728 | 0,937 | 0,828 | 0,398 | 0,913 | 0,836 | 0,778 | 0,958 | 0,835 | 0,653 | 0,933 |
| Algorithm metric average | 0,775 | | | 0,832 | | | 0,713 | | | 0,857 | | | 0,807 | | |
| ran_10k | /* | | | /* | | | /* | | | /* | | | /* | | |

*Configurations with 10,000 pseudo-absences (10k) are not shown, as empty validation blocks occurred for some species, preventing their evaluation. To retain the maximum number of species in the analyses, this option was therefore excluded.

**Appendix C: Detailed overview of SDM framework according to the ODMAP protocol**

Overview, Data, Model, Assessment, and Prediction (ODMAP) protocol (Zurell et al., 2020) provides a standardized framework for transparently and reproducibly reporting species distribution models.

**Table C: Overview, Data, Model, Assessment, and Prediction (ODMAP) protocol**. The following table details our study methodology according to the ODMAP structure. **Data and code availability links are provided in the Overview element, under the *Software, Codes, and Data* section**.

| ODMAP element | Contents |
|---|---|
| **OVERVIEW** | |
| *Authorship* | · **Authors:** Noèmie COLLETTE; Sébastien PINEL; Valérie HINOUX; Joris BERTRAND |
| | · **Contact email:** |
| | · **Title:** Predicting spatiotemporal bioclimatic niche dynamics of endemic Pyrenean plant species under climate change: how many will we lose? |
| | · **DOI:** 10.1101/2025.03.19.644085 |
| *Model objective* | · **SDM purpose:** forecast / transfer |
| | · **Main target output:** continuous and binary bioclimatic suitability maps |
| *Taxon* | Endemic Pyrenean plants |
| *Location* | The Pyrenees massif (Körner *et al.*, 2011, Snethlage *et al.*, 2022), southwestern Europe |
| *Scale of analysis* | · **Spatial Extent (Lon / Lat):** -2,03, 3,29, 41.82, 43.39 (xmin, xmax, ymin, ymax) |
| | · **Spatial resolution:** 1kmx1km |
| | · **Temporal extent/time period:** 1970-2100 |
| | · **Temporal resolution:** 5 periods 1970-2000, 2021-2040, 2041-2060, 2061-2080, and 2081-2100 (designated as current, 2030, 2050, 2070, 2090 respectively) |
| | · **Type of extent boundary:** natural |
| *Biodiversity data overview* | · **Observation type:** citizen science, field survey |
| | · **Response/data type:** presence-only |
| *Type of predictors* | Climatic |
| *Conceptual model* | · **Hypotheses about species-environment relationship:** species distributions are mainly driven by temperature and precipitation. Biotic interactions have a negligible impact. All populations of a given species respond uniformly to climatic variables over time, without accounting for local adaptations or genetic variability. |
| *Assumptions* | (1) Species are in equilibrium with the environment; (2) Biases in the modeling system are minimal; (3) Species niches are conserved over time; (4) All model variables are related to species occurrence. |
| *SDM algorithms* | · **Model algorithms:** Maximum entropy (Maxent), Generalized Linear Model (GLM), Gradient Boosting Model (GBM), Generalized Additive Model (GAM), Random Forest (RF) |

| | |
|---|---|
| | · **Model complexity:** we let the data determine model complexity. Model settings were automatically optimized to achieve the best performance. Tested settings were chosen to generate an intermediately complex response while preventing excessive overfitting yet still covering a sufficiently broad range. |
| | · **Model averaging:** Median of models over a maximum of 10 repetitions of each model algorithm (maximum 50 models). |
| *Model workflow* | **(1)** Species occurrence data downloading & cleaning: Species selection (endemic to the Atlas of Pyrenean Flora (Pironon *et al.*, 2022), downloaded from open databases)), data filtering (>1970, ≤ 1 km uncertainty) and spatial thinning.<br>**(2)** Selecting and processing climate data. Download WorldClim 2.1 current (1970-2000) and future (2021-2100) bioclimatic variables under four Shared Socioeconomic Pathways (SSP126, 245, 370, 585) on 0.5 arcs/min resolution. All future climate projections (CMIP6 GCMs) available were combined into a median raster for each period and SSP. Highly positively correlated variables (r > 0.8) were clustered, selecting the highest-ranked variable from each cluster based on an ecological priority list.<br>**(3)** Environmental constraints sampling of pseudo-absence points. One set per run and species: environmentally constrained pseudo-absences in equal number to presences.<br>**(4)** Individual model fitting. Parameter tuning using four spatial block cross-validation with block size set to two thirds of environmentally independent folds. Evaluation metrics: AUC-PR, Sensitivity (MaxSSS threshold), Boyce Index. The best parameter combination per run was retained based on the score set as calculated as a weighted average of 0.5 Boyce Index, 0.25 AUC-PR, and 0.25 Sensitivity.<br>**(5)** Ensemble modeling generation. Ensemble modeling was applied at two distinct stages: (i) for current conditions, based on model runs calibrated and validated (Boyce Index > 0.5, maximum 5 algorithms x 10 pseudo absence set = 50 models) under present-day climate; and (ii) for future projections, by projecting these same validated models to every combination of future period and scenario, and then creating an ensemble for each period-scenario combination.<br>**(6)** Analysis of bioclimatic niche suitability dynamics. Binarized maps for gain/loss and overlap of suitable habitat, combined with elevation data and latitude for assessment of spatial shifts over time. Identification of bioclimatic hotspot areas using continuous suitability maps. |
| *Software, codes and data* | Computational environment:<br>Processor: Apple M2 Max<br>Total number of cores: 12<br>Total RAM memory: 64 GB<br><br>Analyses were conducted with R version 4.4.1 (R Core Team, 2024) in Rstudio environment (Posit team, 2024).<br><br>Key packages:<br>Retrieval occurrence data from GBIF – packages 'geodata' (Hijmans et al., 2024) and 'rgbif' (Chamberlain & Boettiger, 2017; Chamberlain et al., 2025).<br>Retrieval occurrence data from iNaturalist – package 'rinat' (Barve & Hart, 2025).<br>Data cleaning – packages 'fuzzySim' (Barbosa, 2015) and 'sf' (Pebesma, 2018; Pebesma & Bivand, 2023)<br>Spatial thinning of occurrences, raster manipulation, and model projections – package 'terra' (Hijmans, 2024).<br>Generation of environmental constraints pseudo-absences – package 'flexsdm' (Velazco et al., 2022).<br>MaxEnt – package 'maxnet' (Phillips, 2021). |

Generalized Additive Models (GAM) – package 'mgcv' (Wood, 2017)
Generalized Boosted Regression Models (GBM) – package 'gbm' (Ridgeway & Developers, 2024)
Random Forest – package 'randomForest' (Liaw & Wiener, 2001).
Block cross-validation – package 'blockCV' (Valavi et al., 2019).
Computation of AUC-PR – package 'PRROC' package (Keilwagen et al., 2014; Grau et al., 2015).
Computation of sensitivity, Boyce Index and MaxSSS threshold– package 'modEvA' (Barbosa et al., 2013).
Computation of environmental similarity index (Multivariate Environmental Similarity Surface, MESS) – package 'predicts' (Hijmans, 2023).
Parallelization using mclapply – package 'parallel' (R Core Team, 2024) *(note: this approach does not work on Windows and Linux).*

```
> sessionInfo()
attached base packages:
 [1] parallel splines  grid   tools   stats
 [6] graphics grDevices utils   datasets methods
[11] base
other attached packages:
 [1] qpdf_1.3.3      ggsignif_0.6.4
 [3] dunn.test_1.3.6   fields_16.2
 [5] viridisLite_0.4.2  spam_2.10-0
 [7] ggtext_0.1.2     RColorBrewer_1.1-3
 [9] raster_3.6-30    sp_2.1-4
[11] flexsdm_1.3.4    MASS_7.3-60.2
[13] PRROC_1.3.1     Hmisc_5.1-3
[15] ecospat_4.1.1    reshape2_1.4.4
[17] blockCV_3.1-4    plotmo_3.6.4
[19] plotrix_3.8-4   Formula_1.2-5
[21] modEvA_3.18.2    corrplot_0.94
[23] randomForest_4.7-1.2 gbm_2.2.2
[25] gam_1.22-5      foreach_1.5.2
[27] maxnet_0.1.4     predicts_0.1-11
[29] collinear_1.1.1   lubridate_1.9.3
[31] forcats_1.0.0    purrr_1.0.2
[33] readr_2.1.5     tidyr_1.3.1
[35] tibble_3.2.1     tidyverse_2.0.0
[37] gridExtra_2.3    stringr_1.5.1
[39] tidyterra_0.6.1   fuzzySim_4.10.7
[41] rgbif_3.8.0     rinat_0.1.9
[43] sf_1.0-17      dplyr_1.1.4
[45] usethis_3.0.0    ggplot2_3.5.1
[47] geodata_0.6-2    terra_1.7-78

loaded via a namespace (and not attached):
 [1] shape_1.4.6.1   rstudioapi_0.16.0
 [3] jsonlite_1.8.8   wk_0.9.3
 [5] magrittr_2.0.3   farver_2.1.2
 [7] rmarkdown_2.28   fs_1.6.4
 [9] ragg_1.3.2     vctrs_0.6.5
[11] askpass_1.2.0   base64enc_0.1-3
[13] htmltools_0.5.8.1 curl_5.2.2
[15] s2_1.1.7      KernSmooth_2.23-24
[17] htmlwidgets_1.6.4 plyr_1.8.9
[19] commonmark_1.9.1  whisker_0.4.1
[21] lifecycle_1.0.4  iterators_1.0.14
[23] pkgconfig_2.0.3  Matrix_1.7-0
[25] R6_2.5.1      fastmap_1.2.0
[27] digest_0.6.37   colorspace_2.1-1
```

| | |
|---|---|
| | [29] patchwork_1.3.0 textshaping_0.4.0<br>[31] labeling_0.4.3 fansi_1.0.6<br>[33] urltools_1.7.3 timechange_0.3.0<br>[35] httr_1.4.7 mgcv_1.9-1<br>[37] compiler_4.4.1 proxy_0.4-27<br>[39] withr_3.0.1 doParallel_1.0.17<br>[41] htmlTable_2.4.3 backports_1.5.0<br>[43] DBI_1.2.3 spThin_0.2.0<br>[45] maps_3.4.2 classInt_0.4-10<br>[47] oai_0.4.0 units_0.8-5<br>[49] foreign_0.8-86 nnet_7.3-19<br>[51] glue_1.7.0 nlme_3.1-164<br>[53] gridtext_0.1.5 checkmate_2.3.2<br>[55] cluster_2.1.6 generics_0.1.3<br>[57] gtable_0.3.5 tzdb_0.4.0<br>[59] class_7.3-22 data.table_1.16.0<br>[61] hms_1.1.3 xml2_1.3.6<br>[63] utf8_1.2.4 markdown_1.13<br>[65] pillar_1.9.0 lattice_0.22-6<br>[67] survival_3.6-4 tidyselect_1.2.1<br>[69] knitr_1.48 crul_1.5.0<br>[71] xfun_0.47 stringi_1.8.4<br>[73] lazyeval_0.2.2 evaluate_1.0.0<br>[75] codetools_0.2-20 httpcode_0.3.0<br>[77] kernlab_0.9-33 cli_3.6.3<br>[79] rpart_4.1.23 systemfonts_1.1.0<br>[81] munsell_0.5.1 Rcpp_1.0.13<br>[83] triebeard_0.4.1 dotCall64_1.1-1<br>[85] glmnet_4.1-8 scales_1.3.0<br>[87] e1071_1.7-16 crayon_1.5.3<br>[89] rlang_1.1.4 |
| | · **Data availability:** code available at [DOI : 10.5281/zenodo.17412264] |
| | · **Data availability:** GBIF Occurrences data: https://doi.org/10.15468/dl.r84ym3 and the exact GBIF dataset used (as the data were downloaded using the geodata package while the DOI was generated with the rgbif package, which may result in minor discrepancies): [DOI : 10.5281/zenodo.17412264] iNaturalist [DOI : 10.5281/zenodo.17412264] Environmental data: https://www.worldclim.org/ |
| DATA | |
| *Biodiversity data* | · **Taxon names:** *Achillea chamaemelifolia, Allium pyrenaicum, Androsace ciliata, Androsace laggeri, Androsace pyrenaica, Antirrhinum molle, Antirrhinum sempervirens, Arenaria oscensis, Armeria bubanii, Campanula jaubertiana, Campanula precatoria, Centaurea costae, Centaurea emigrantis, Cerastium pyrenaicum, Cirsium glabrum, Delphinium montanum, Dianthus benearnensis, Draba subnivalis, Endressia pyrenaica, Erodium lucidum, Festuca altopyrenaica, Festuca borderei, Festuca pyrenaica, Galeopsis pyrenaica, Galium cespitosum, Iberis bernardiana, Iberis spathulata, Knautia lebrunii, Leucanthemum graminifolium, Linaria bubanii, Medicago hybrida, Minuartia cerastiifolia, Narcissus bicolor, Onobrychis pyrenaica, Petrocoptis crassifolia, Petrocoptis hispanica, Petrocoptis montsicciana, Pinguicula longifolia, Ramonda myconi, Ranunculus pyrenaeus, Ranunculus ruscinonensis, Rhaponticum centauroides, Salix pyrenaica, Santolina benthamiana, Saponaria caespitosa, Saxifraga aquatica, Saxifraga aretioides, Saxifraga geranioides, Saxifraga hariotii, Saxifraga intricata, Saxifraga media, Saxifraga umbrosa, Scrophularia pyrenaica, Silene borderei, Thalictrum macrocarpum, Thymelaea calycina, Trisetum baregense, Viola diversifolia, Xatartia* |

| | |
|---|---|
| | *scabra.* |
| | · **Ecological level:** species |
| | · **Data source: GBIF:** https://doi.org/10.15468/dl.r84ym3 , iNaturalist: https://www.inaturalist.org/ [accessed 18 February 2025] [DOI : 10.5281/zenodo.17412264] |
| | · **Sampling design:** opportunistic & standardized monitoring sampling |
| | · **Sample size per taxon:** |

| **Species** | **Raw (study area, 1970+, uncertainty ≤ 1 km)** | **Gridded (after spatial thinning)** |
|---|---|---|
| *Achillea chamaemelifolia* | 182 | 132 |
| *Allium pyrenaicum* | 25 | 25 |
| *Androsace ciliata* | 256 | 112 |
| *Androsace laggeri* | 355 | 258 |
| *Androsace pyrenaica* | 311 | 133 |
| *Antirrhinum molle* | 224 | 142 |
| *Antirrhinum sempervirens* | 194 | 138 |
| *Arenaria oscensis* | 26 | 22 |
| *Armeria bubanii* | 103 | 80 |
| *Campanula jaubertiana* | 17 | 15 |
| *Campanula precatoria* | 228 | 171 |
| *Centaurea costae* | 42 | 39 |
| *Centaurea emigrantis* | 61 | 51 |
| *Cerastium pyrenaicum* | 204 | 112 |
| *Cirsium glabrum* | 182 | 88 |
| *Delphinium montanum* | 138 | 40 |
| *Dianthus benearnensis* | 164 | 130 |
| *Draba subnivalis* | 100 | 81 |
| *Endressia pyrenaica* | 325 | 202 |
| *Erodium lucidum* | 53 | 41 |
| *Festuca altopyrenaica* | 17 | 17 |
| *Festuca borderei* | 105 | 92 |
| *Festuca pyrenaica* | 124 | 103 |
| *Galeopsis pyrenaica* | 530 | 369 |
| *Galium cespitosum* | 238 | 172 |
| *Iberis bernardiana* | 205 | 118 |
| *Iberis spathulata* | 255 | 148 |
| *Knautia lebrunii* | 78 | 67 |
| *Leucanthemum graminifolium* | 153 | 73 |
| *Linaria bubanii* | 49 | 40 |
| *Medicago hybrida* | 540 | 370 |
| *Minuartia cerastiifolia* | 80 | 61 |
| *Narcissus bicolor* | 237 | 194 |
| *Onobrychis pyrenaica* | 65 | 39 |
| *Petrocoptis crassifolia* | 75 | 61 |
| *Petrocoptis hispanica* | 95 | 58 |
| *Petrocoptis montsicciana* | 45 | 43 |
| *Pinguicula longifolia* | 313 | 106 |
| *Ramonda myconi* | 2105 | 985 |
| *Ranunculus pyrenaeus* | 854 | 583 |
| *Ranunculus ruscinonensis* | 39 | 39 |
| *Rhaponticum centauroides* | 27 | 22 |
| *Salix pyrenaica* | 1112 | 668 |
| *Santolina benthamiana* | 79 | 66 |
| *Saponaria caespitosa* | 227 | 139 |
| *Saxifraga aquatica* | 805 | 448 |

| | |
|---|---|
| | *Saxifraga aretioides* 328 199<br>*Saxifraga geranioides* 937 525<br>*Saxifraga hariotii* 151 106<br>*Saxifraga intricata* 215 165<br>*Saxifraga media* 424 243<br>*Saxifraga umbrosa* 926 646<br>*Scrophularia pyrenaica* 173 95<br>*Silene borderei* 52 36<br>*Thalictrum macrocarpum* 298 156<br>*Thymelaea calycina* 93 34<br>*Trisetum baregense* 51 49<br>*Viola diversifolia* 256 87<br>*Xatartia scabra* 208 74 |
| | · **Mask:** We clipped all data to the boundary of the study region. |
| | · **Details on scaling:** Spatial thinning, retaining one occurrence per grid cell. |
| | · **Details on data cleaning/filtering steps:** Occurrences since 1970 within the study area and ≤ 1 km coordinate uncertainty were retained, fitting the resolution of environmental variables. The definition of broad distribution is based on the distribution of occurrences across spatial blocks during the model assessment step. To retain a species, each block must contain at least one presence. Records with suspicious coordinates were removed, and the details of these coordinates are available in the code, section "#Removing erroneous coordinates". Species eliminated and reason are documented in "ATLAS OF PYRENEAN FLORA – ENDEMIC.xlsx" in Species eliminated sheet. |
| | · **Details on background data derivation:** We adopt environmentally constrained pseudo-absences in equal number to presences, as its consistently yielded the best evaluation scores across all algorithms against other configuration (environmentally constrained or random sampling with pseudo-absences equal to the number of presences, three times the number of presences, or fixed at 10,000, as recommended by Barbet-Massin et al., 2012 and the Biomod2 team (Gueguen et al., 2025)). We generated 10 independent replicates (10 pseudo-absence sets) per species (Biomod2 team recommendation). |
| | · **Details on potential errors and biases in data:** The biases encountered are those related to citizen science (collected using different sampling efforts and methodology, misidentification potential, collected in more accessible areas...). |
| *Data partitioning* | · **Selection of training data:** 4 block cross-validation, with blocks assigned to the training set in each iteration |
| | · **Selection of validation data:** Validation data were drawn from spatially independent blocks (determining through blockCV::cv_spatial_autocor function and divided by 1.5 to balance data sufficiency, resulting in a block size of [79,785/1.5] m) excluded from the training set during each cross-validation iteration. |
| *Predictor variables* | · **State predictor variables used:** list of 19 bioclimatic variables (11 related to temperature and 8 to precipitation) available on WorldClim 2.1 (Fick & Hijmans, 2017) |
| | · **Details on data sources:** WorldClim 2.1 https://www.worldclim.org/ [accessed 16 January 2025] |
| | · **Spatial resolution and spatial extent of raw data:** 1 km x 1 km, world |

| | |
|---|---|
| | · **Map projection (coordinate reference system):** EPSG:4326 |
| | · **Temporal resolution and temporal extent of raw data:** 1970-2000 |
| | · **Details on data processing and on spatial, temporal and thematic scaling:** We clipped all data to the boundary of the study region. |
| | · **Details on measurement errors and bias, when known:** See Fick & Hijmans, 2017. Potential biases may arise from interpolation errors, particularly in our study area with sparse meteorological stations or complex topography. |
| *Transfer data for projection* | · **Details on data sources, spatial extent and resolution:** same as *Predictor variables* |
| | · **Temporal extent/time period:** 2021-2100 |
| | · **Temporal resolution:** 4 periods 2021-2040, 2041-2060, 2061-2080, and 2081-2100 (designated as 2030, 2050, 2070, 2090 respectively) |
| | · **Models and scenarios used:** All General Circulation Models available on WorldClim 2.1 for Shared Socioeconomic Pathways (SSPs) SSP126, SSP245, SSP370 & SSP585: ACCESS-CM2, CMCC-ESM2, EC-Earth3-Veg, UKESM1-0-LL, GISS-E2-1-G, INM-CM5-0, IPSL-CM6A-LR, MIROC6, MPI-ESM1-2-HR, MRI-ESM2-0, BCC-CSM2-MR. |
| | · **Details on data processing and scaling:** We clipped all data to the boundary of the study region. The General Circulation Models were aggregated by taking the median for each period and SSP. |
| | · **Quantification of novel environmental conditions and novel environmental combinations:** environmental novelty was assessed using the Multivariate Environmental Similarity Surface (Elith et al., 2010). |
| MODEL | |
| *Variable pre-selection* | The initial variables were selected based on their availability for future periods. |
| *Multicollinearity* | To reduce redundancy in models, correlated variables (Pearson r > 0.8) at presence locations for each species were grouped into clusters. Negatively correlated variables can capture complementary environmental gradients (e.g. warm summers vs. cold winters) and were therefore kept in the analysis. From each cluster, the highest-ranked variable was retained based on a predefined ecological priority list : BIO1 & BIO12 > BIO5 (Román-Palacios & Wiens, 2020) > BIO6 > BIO7 > BIO15 > BIO10 > BIO18 > BIO11 > BIO13 > BIO14 > BIO16 > BIO17 > BIO4 > BIO8 > BIO9 > BIO19 > BIO3 > BIO2. |

| | |
|---|---|
| *Model settings* | **· Models settings:**<br>For model tuning:<br>GBM:<br>n.trees_vals <- c(100, 200, 500)<br>interaction.depth_vals <- c(1, 2, 3)<br>shrinkage_vals <- c(0.01, 0.1, 0.05)<br>n.minobsinnode_vals <- c(5, 10, 15)<br>bag.fraction_vals = c(0.4, 0.5, 0.6)<br>MXT:<br>regmult_vals <- c(0.5, 1, 2, 3)<br>classes_vals <- c("l", "q", "lq", "lqh", "lqhp", "lqhpt")<br>RF:<br>ntree_vals <- c(500, 1000)<br>mtry_vals <- c(floor(sqrt(length(vars_sel))), floor(length(vars_sel) / 2), length(vars_sel))<br>nodesize_vals <- c(5, 10, 20)<br>GAM:<br>sp_vals <- c(0.1, 0.5, 1, 2, 5)<br>GLM:<br>glm_form <- as.formula(paste("presence ~", paste("poly(", vars_sel, ", 2)", collapse = " + "), "+", paste(vars_sel, collapse = ":")))<br>glm_mod <- glm(formula = glm_form, family = binomial(link = "logit"), data = dat_random)<br>glm_mod_opt <- stepAIC(glm_mod, direction = "both")<br><br>Default setting for the other parameters.<br><br>For modeling:<br>GBM:<br>gbm_mod <- gbm::gbm(formula = gbm_form, data = dat_env_const, shrinkage = best_shrinkage, interaction.depth = best_interaction.depth, n.trees = best_n.trees, n.minobsinnode = best_n.minobsinnode, bag.fraction = best_bag.fraction, distribution = "bernoulli")<br>MXT:<br>mxt_mod <- maxnet::maxnet(p = dat_random$presence, data = dat_random[, vars_sel], regmult = best_regmult, maxnet.formula(p = dat_random$presence, data = dat_random[, vars_sel], classes = best_classes))<br>RF:<br>rf_form <- reformulate(termlabels = vars_sel, response = "presence_fact")<br>rf_mod <- randomForest::randomForest(formula = rf_form, data = dat_env_const, na.action = na.exclude, keep.forest = TRUE, importance = TRUE, nodesize=best_nodesize, mtry=best_mtry, ntree = best_ntree)<br>GAM:<br>gam_form <- reformulate(termlabels = supply(vars_sel, function(v) paste0("s(", v, ", sp = ", best_sp, ")")), response = "presence")<br>gam_mod <- mgcv::gam(formula = gam_form, family = binomial, data = dat_random, weights = weights_GAM, method = "REML")<br>GLM:<br>glm_mod <- glm(formula = glm_form, family = binomial(link = "logit"), data = dat_random, weights = weights_GLM)<br><br>GLM, GAM, GBM and RF prevalence (presence/absence ratio) was removed to convert presence/absence model into a favorability model with fuzzySim::Fav function. |
| *Model estimates* | **· Assessment of model coefficients:** Not analyzed. |

| | |
|---|---|
| | · **Assessment of variable importance:** Variable importance was assessed by quantifying how much deviance each variable explains. Two summary tables of variable importance are generated upon code execution, one presenting the mean scores per predictor and algorithm across all runs, and another providing details for each individual run. |
| *Model selection / Model averaging / Ensembles* | · **Model selection strategy:** Models were selected based on cross-validation performance using three metrics: Boyce Index, Sensitivity, and AUC-PR. For each algorithm and iteration, the set of parameters yielding the highest performance score (calculated as a weighted average of 0.5 Boyce Index, 0.25 AUC-PR, and 0.25 Sensitivity) was retained. Only models with a Boyce Index > 0.5 were included in the final ensemble model to ensure robustness and reliability. |
| | · **Model averaging method / ensemble method:** Ensemble modeling was applied at two distinct stages: (i) for current conditions, based on model runs calibrated and validated (Boyce Index > 0.5, maximum 5 algorithms x 10 pseudo absence set = 50 models) under present-day climate; and (ii) for future projections, by projecting these same validated models to every combination of future period and scenario, and then creating an ensemble for each period–scenario combination. Consensus ensemble maps were computed as pixel-wise median across all algorithms and replicates. |
| *Non-independence correction/analyses* | · **Method for addressing spatial autocorrelation in residuals:** Spatial autocorrelation was addressed by analyzing a variogram of the environmental predictors to determine an appropriate block size for spatial cross-validation (determining through blockCV::cv_spatial_autocor function and divided by 1.5 to balance data sufficiency, resulting in a block size of [79,785/1.5] m). Residuals were not explicitly analyzed for spatial autocorrelation after model fitting. |
| *Threshold selection* | · **Details on threshold selection:** Continuous maps were converted into binary 'suitable–unsuitable' maps using a threshold based on maximising the sum of sensitivity (True Positive Rate) and specificity (True Negative Rate) (Liu et al., 2013, 2016). |
| ASSESSMENT | |
| *Performance statistics* | · **Performance statistics estimated validation data:** Performance metrics are computed for all models across all parameters sets. After selecting the best models (one per algorithm for each run, based on a weighted score calculated as a weighted average of 0.5 Boyce Index, 0.25 AUC-PR, and 0.25 Sensitivity), the values of each metric are averaged to provide an overall assessment per algorithm (with standard deviation).<br>The same approach is applied to the models selected for the ensemble model and averaged across algorithms to be used as ensemble model performance. Algorithms performances were compared using Kruskal-Wallis, with Dunn's post hoc test (Bonferroni correction) identifying pairwise differences between algorithm for each evaluation metric.<br>In the code provided, it is also possible to evaluate the ensemble model with cross-validation, but this is not executed here as it may lead to the loss of additional species. This is because changes in the pseudo-absence/presence ratio (as all pseudo-absence data from individual models are merged) can cause previously valid blocks (containing presence records) to become empty. |
| *Plausibility check* | · **Response plots:** Response plots were generated for all 10 models per algorithm and saved in each species' folder upon code execution. No plausibility checks were conducted. |
| | · **Expert judgements:** Output binary maps and predicted trends were supported by finding from previous study (for *Delphinium montanum,* Salvado *et al.*, 2022) and align with observed patterns in the Pyrenees (Lenoir et al., 2008; Pauli et al., 2012; Ameztegui et al., 2016) and Alps, |

| | |
|---|---|
| | another mountain range in Europe (Dirnböck et al., 2011; Dullinger et al., 2012; Hülber et al., 2016; Dagnino et al., 2020; Rota et al., 2022). |
| PREDICTION | |
| *Prediction output* | · **Prediction units:** Bioclimatic niche suitability maps are expressed on a continuous scale (0–1). Gain and loss of habitat were estimated from binary predictions (suitable (1)/unsuitable (0)). |
| *Uncertainty quantification* | · **Algorithmic uncertainty:** A 5% divergence threshold was applied to identify pixels showing significant disagreement among the predictions of the algorithms retained for the median ensemble model. This calculation provides an indication of where predictions are less consistent between algorithms; these pixels were masked on the current distribution map to focus on more reliable areas. |
| | · **Uncertainty in scenarios:** No quantification of scenario uncertainty was performed. |
| | · **Visualization/treatment of novel environments:** To assess SDM reliability under novel conditions (i.e. conditions in future periods deviating from the calibration range), the Multivariate Environmental Similarity Surface (MESS) index (Elith et al., 2010) was applied. Negative MESS values, indicating extrapolation, were masked for visualization on future distribution maps to highlight most reliable predictions. |

**Appendix D: Current and future bioclimatic niche suitability continuous maps for 59 endemic Pyrenean species**

This appendix provides continuous maps of bioclimatic niche suitability for 59 endemic plant species in the Pyrenees, modeled under current and future climatic conditions. Suitability is represented as a gradient from blue (low bioclimatic suitability) to yellow (high bioclimatic suitability). The maps are displayed in alphabetical order by species name.

List of species with displayed maps:

*Achillea chamaemelifolia, Allium pyrenaicum, Androsace ciliata, Androsace laggeri, Androsace pyrenaica, Antirrhinum molle, Antirrhinum sempervirens, Arenaria oscensis, Armeria bubanii, Campanula jaubertiana, Campanula precatoria, Centaurea costae, Centaurea emigrantis, Cerastium pyrenaicum, Cirsium glabrum, Delphinium montanum, Dianthus benearnensis, Draba subnivalis, Endressia pyrenaica, Erodium lucidum, Festuca altopyrenaica, Festuca borderei, Festuca pyrenaica, Galeopsis pyrenaica, Galium cespitosum, Iberis bernardiana, Iberis spathulata, Knautia lebrunii, Leucanthemum graminifolium, Linaria bubanii, Medicago hybrida, Minuartia cerastiifolia, Narcissus bicolor, Onobrychis pyrenaica, Petrocoptis crassifolia, Petrocoptis hispanica, Petrocoptis montsicciana, Pinguicula longifolia, Ramonda myconi, Ranunculus pyrenaeus, Ranunculus ruscinonensis, Rhaponticum centauroides, Salix pyrenaica, Santolina benthamiana, Saponaria caespitosa, Saxifraga aquatica, Saxifraga aretioides, Saxifraga geranioides, Saxifraga hariotii, Saxifraga intricata, Saxifraga media, Saxifraga umbrosa, Scrophularia pyrenaica, Silene borderei, Thalictrum macrocarpum, Thymelaea calycina, Trisetum baregense, Viola diversifolia, Xatartia scabra.*

For each species, we provide:

- Current bioclimatic niche suitability based on current climatic conditions (1970-2000), with areas of uncertainty (standard deviation > 5% between models) masked in black. The colored horizontal lines are a graphical artefact.
- Future projections for four periods (2021-2040, 2041-2060, 2061-2080, 2081-2100 designated as 2030, 2050, 2070, 2090), under four Shared Socioeconomic Pathways (SSP126, SSP245, SSP370, SSP585, best to worst-case climate scenarios, respectively), with areas of uncertainty (climatic conditions beyond the calibration range) masked in grey.

Current bioclimatic suitability in the Pyrenees for *Achillea chamaemelifolia*

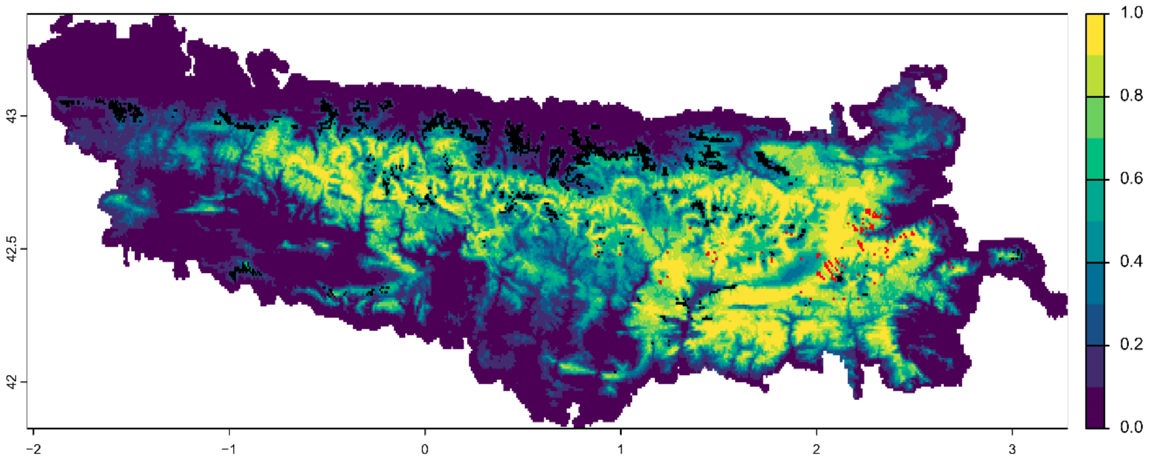


Symbols:

- Variability >=5% among predictions
- Field observation

Future bioclimatic suitability in the Pyrenees by 2090 for *Achillea chamaemelifolia*

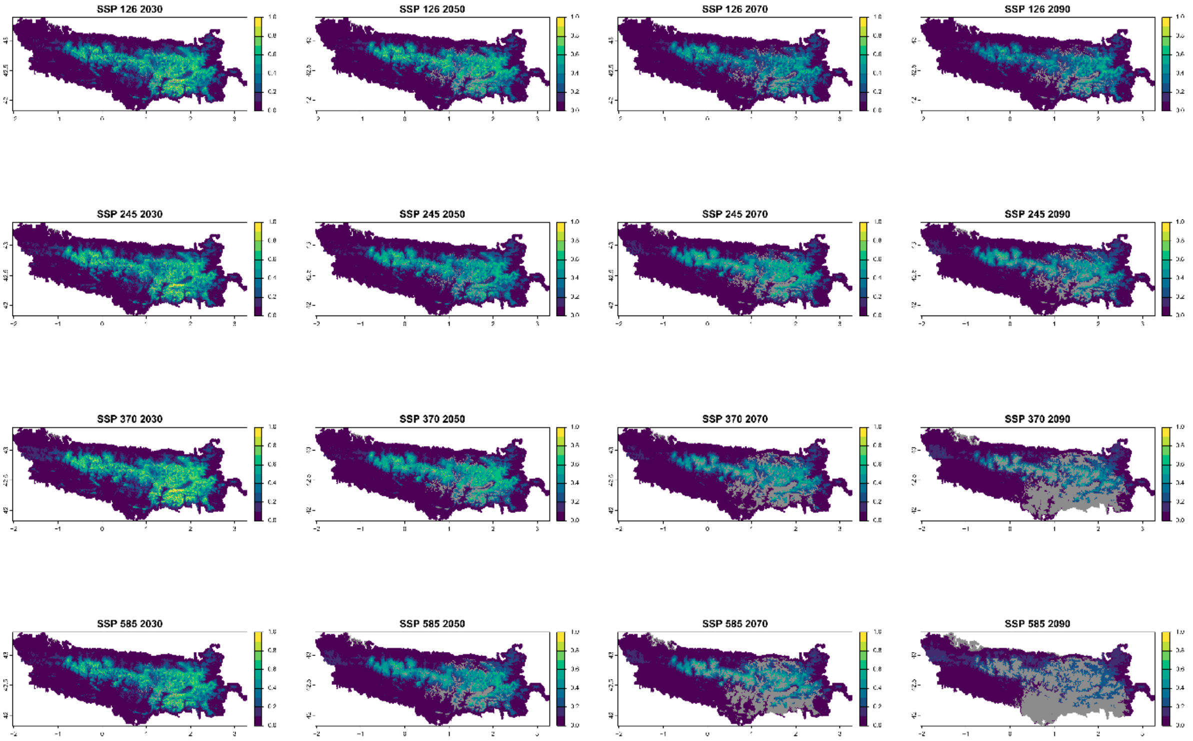


Legend: Non−analogous bioclimatic conditions (MESS < 0)

Current bioclimatic suitability in the Pyrenees for *Allium pyrenaicum*

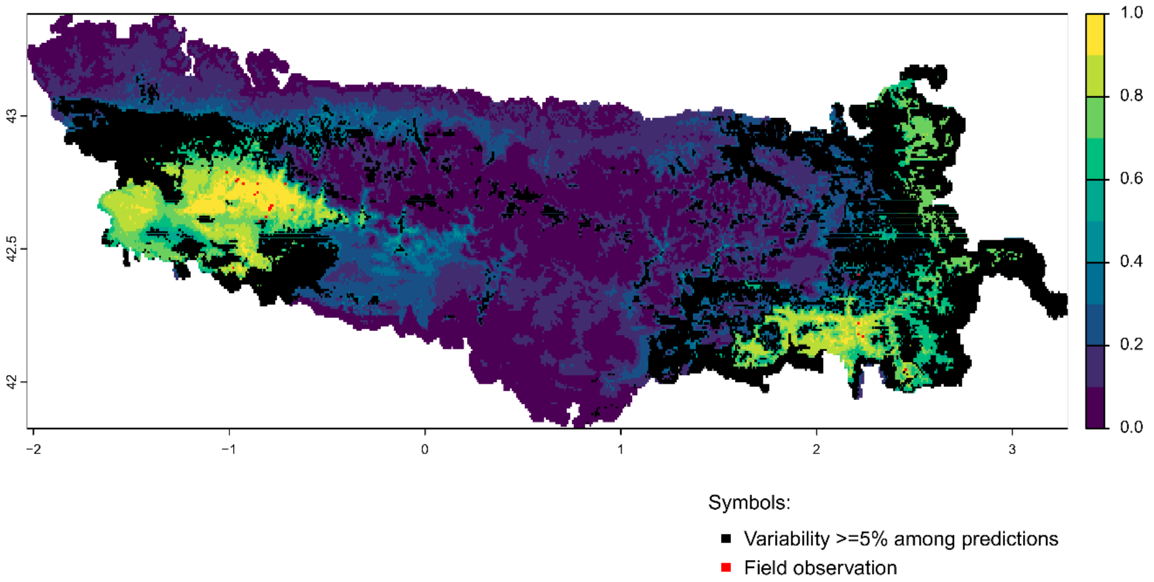


Future bioclimatic suitability in the Pyrenees by 2090 for *Allium pyrenaicum*

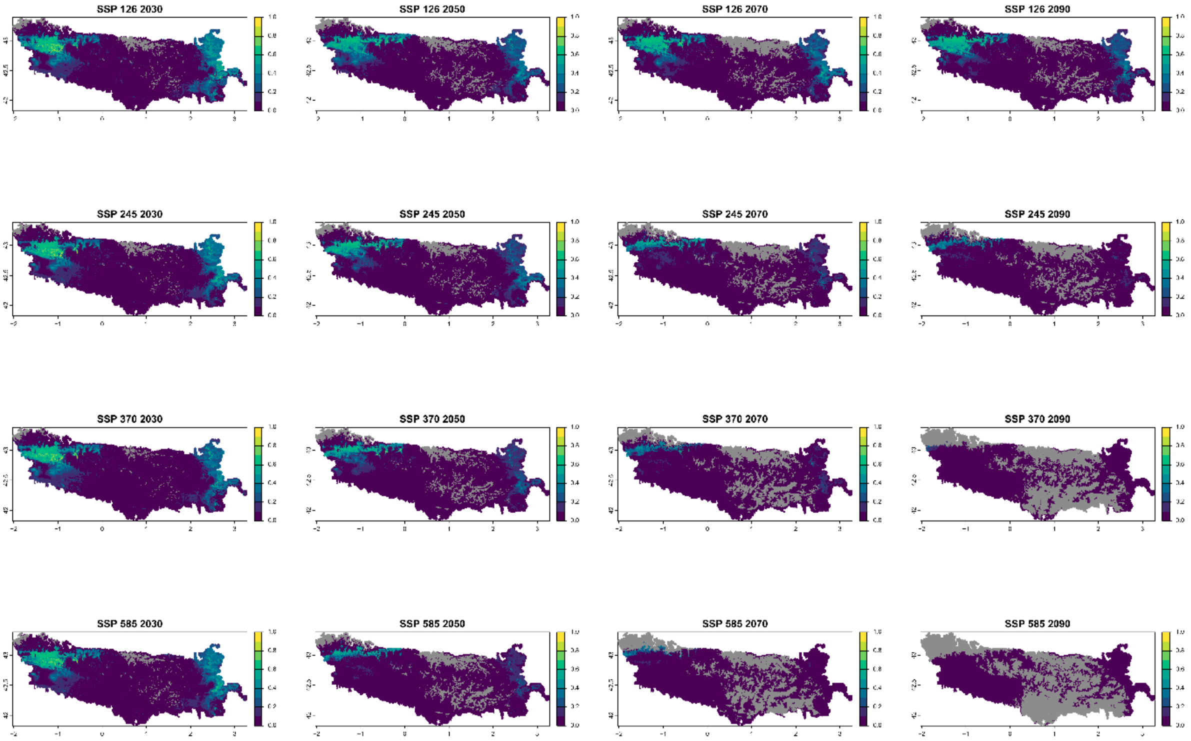


Legend: Non−analogous bioclimatic conditions (MESS < 0)

Current bioclimatic suitability in the Pyrenees for *Androsace ciliata*

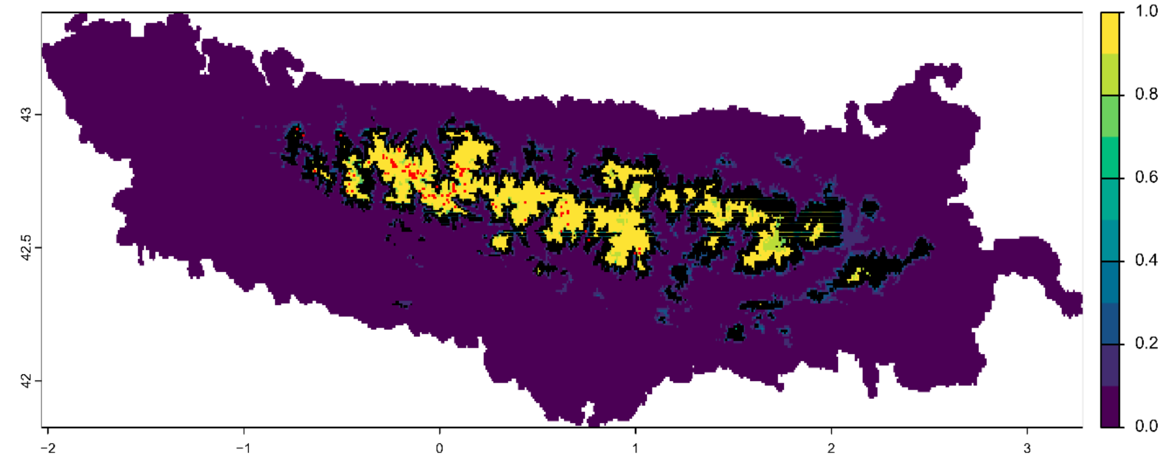


Symbols:

- Variability >=5% among predictions
- Field observation

Future bioclimatic suitability in the Pyrenees by 2090 for *Androsace ciliata*

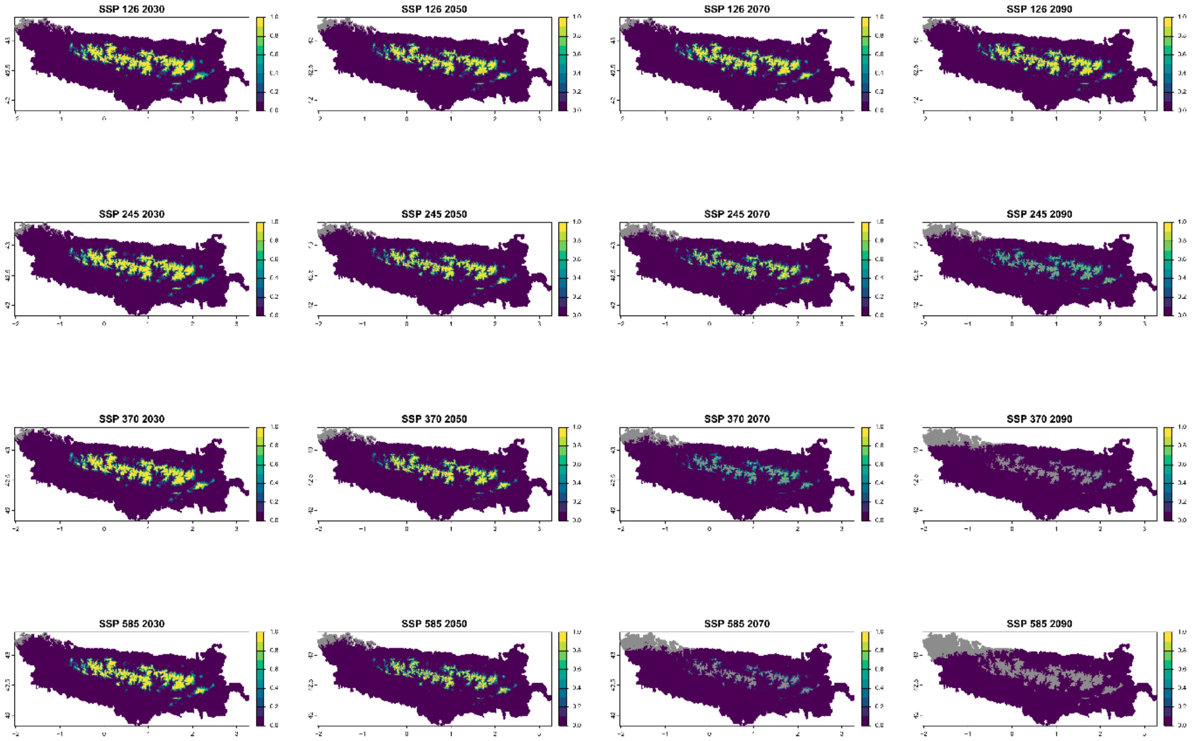


Legend: Non−analogous bioclimatic conditions (MESS < 0)

Current bioclimatic suitability in the Pyrenees for *Androsace laggeri*

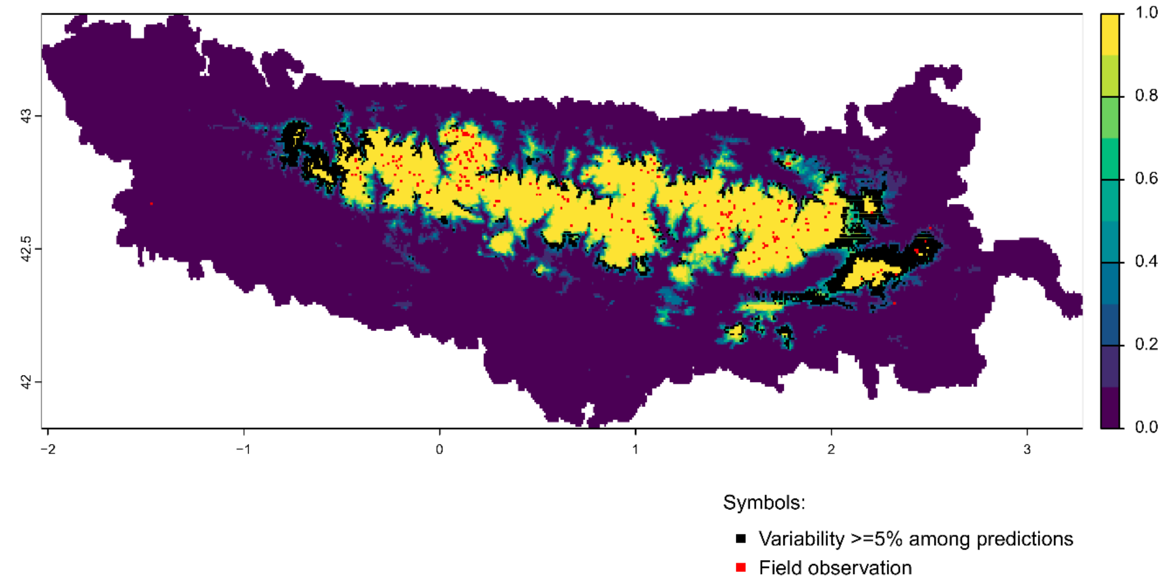


Future bioclimatic suitability in the Pyrenees by 2090 for *Androsace laggeri*

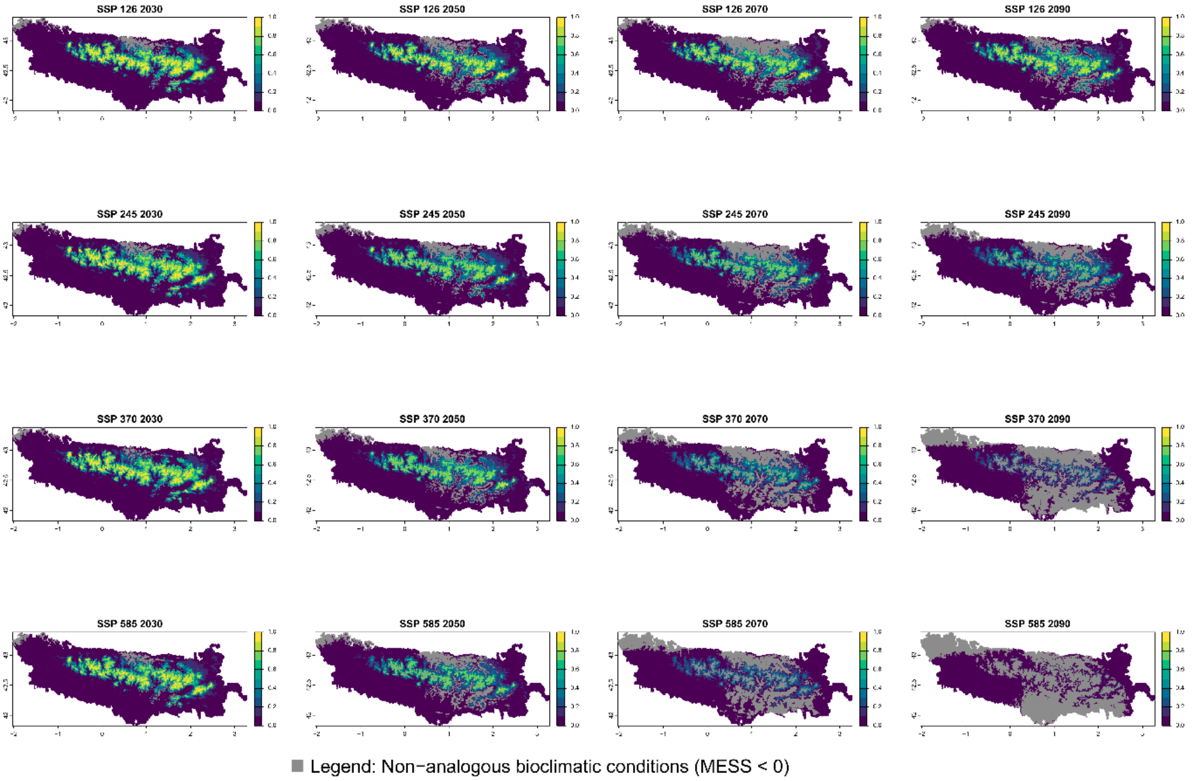

Current bioclimatic suitability in the Pyrenees for *Androsace pyrenaica*

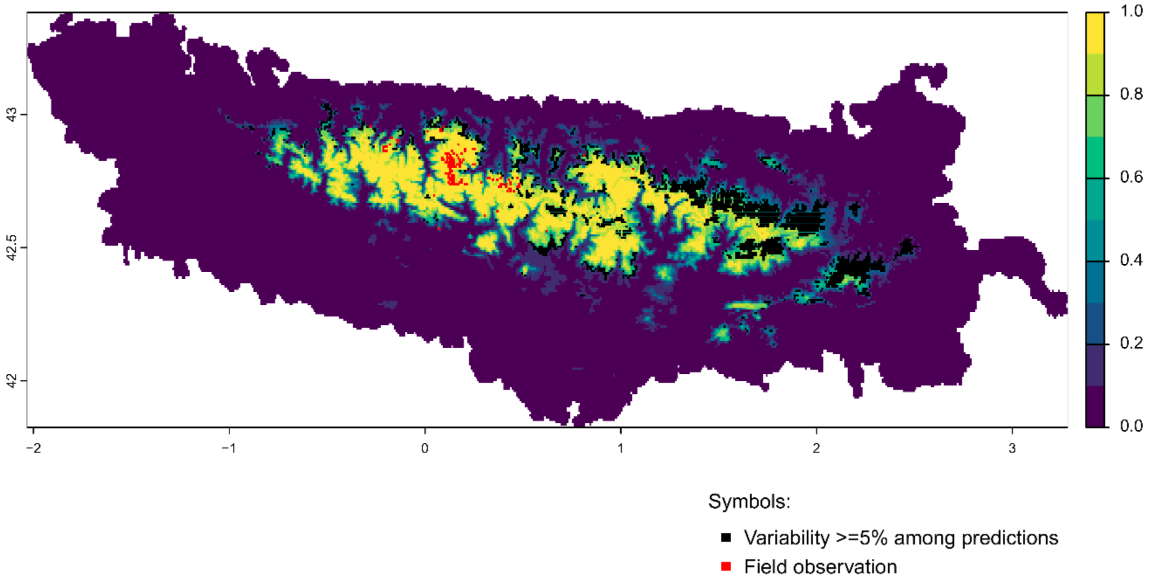


Future bioclimatic suitability in the Pyrenees by 2090 for *Androsace pyrenaica*

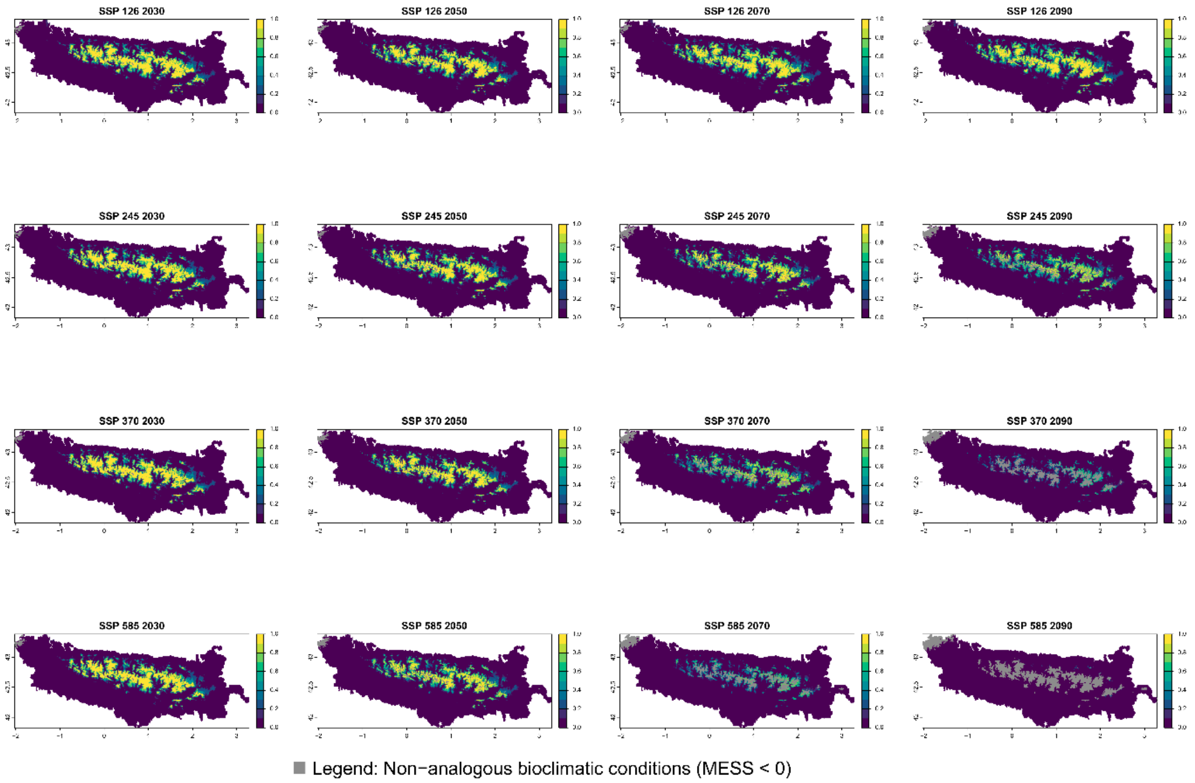

Current bioclimatic suitability in the Pyrenees for *Antirrhinum molle*

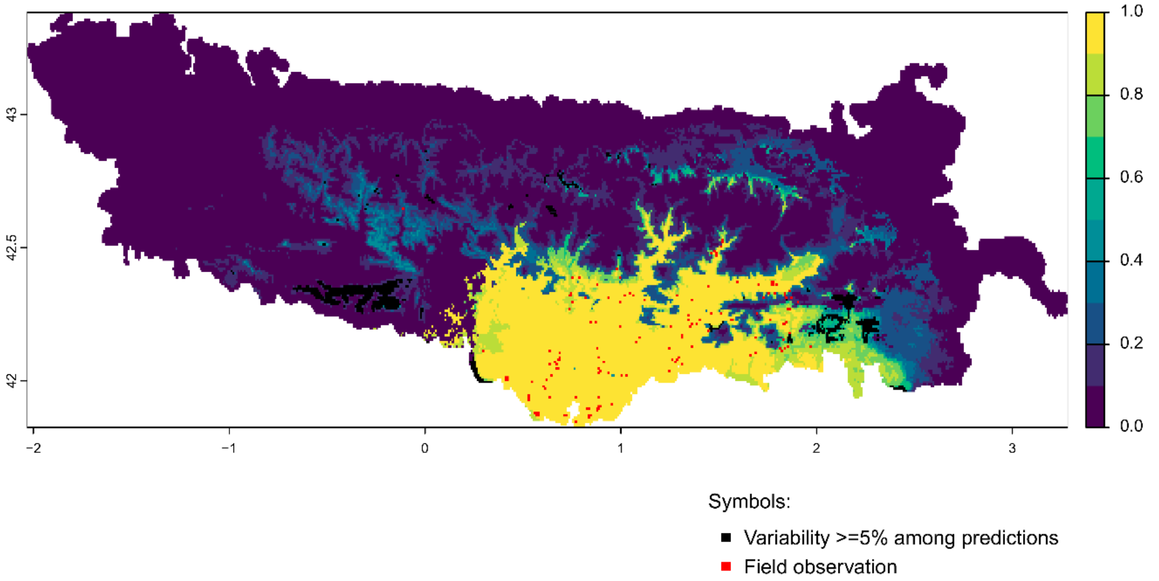


Future bioclimatic suitability in the Pyrenees by 2090 for *Antirrhinum molle*

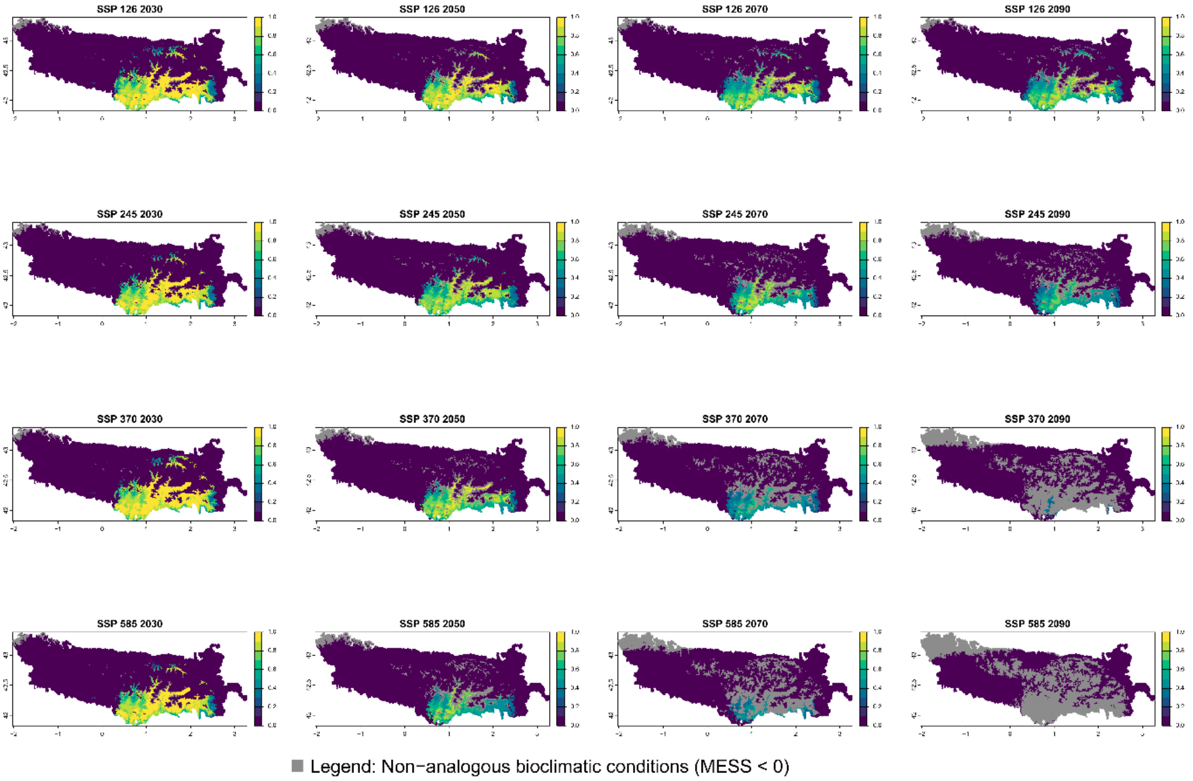

Current bioclimatic suitability in the Pyrenees for *Antirrhinum sempervirens*

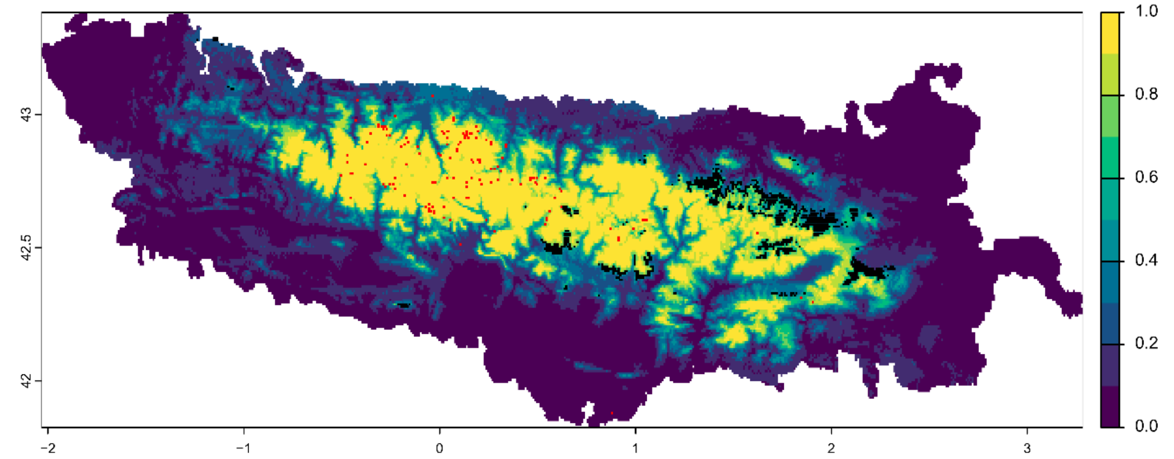


Symbols:
- ■ Variability >=5% among predictions
- ■ Field observation

Future bioclimatic suitability in the Pyrenees by 2090 for *Antirrhinum sempervirens*

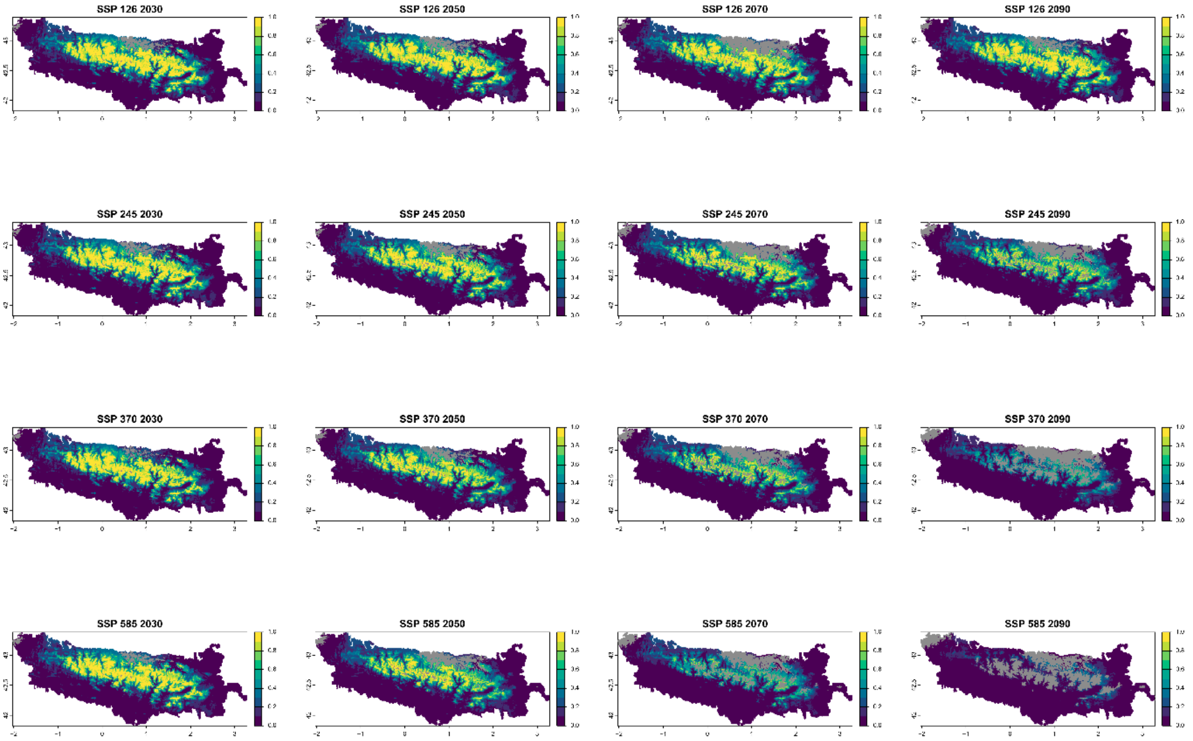


■ Legend: Non−analogous bioclimatic conditions (MESS < 0)

Current bioclimatic suitability in the Pyrenees for *Arenaria oscensis*

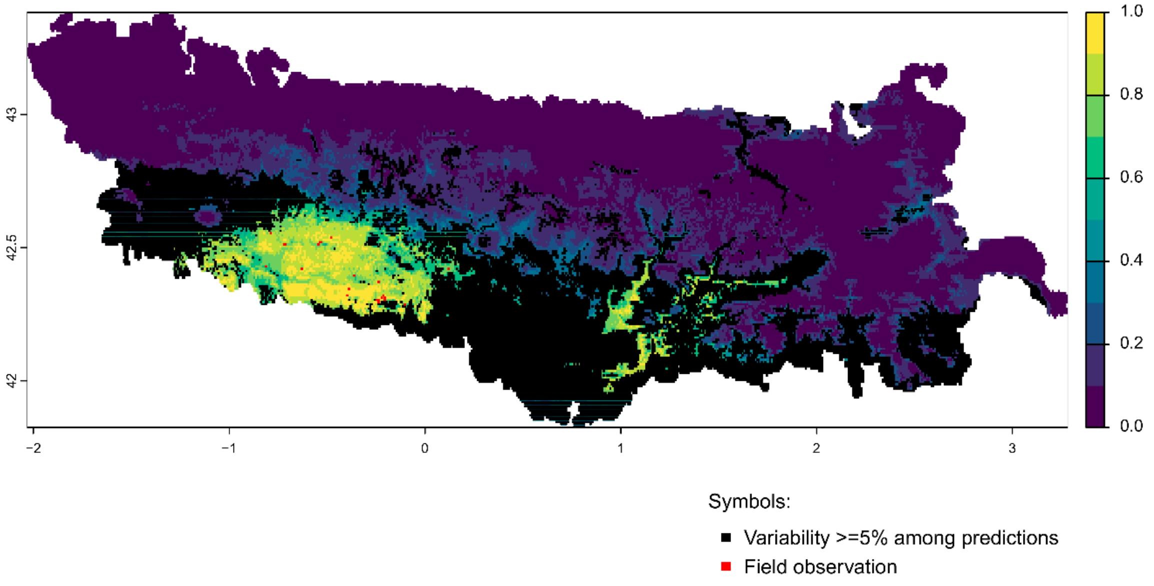


Future bioclimatic suitability in the Pyrenees by 2090 for *Arenaria oscensis*

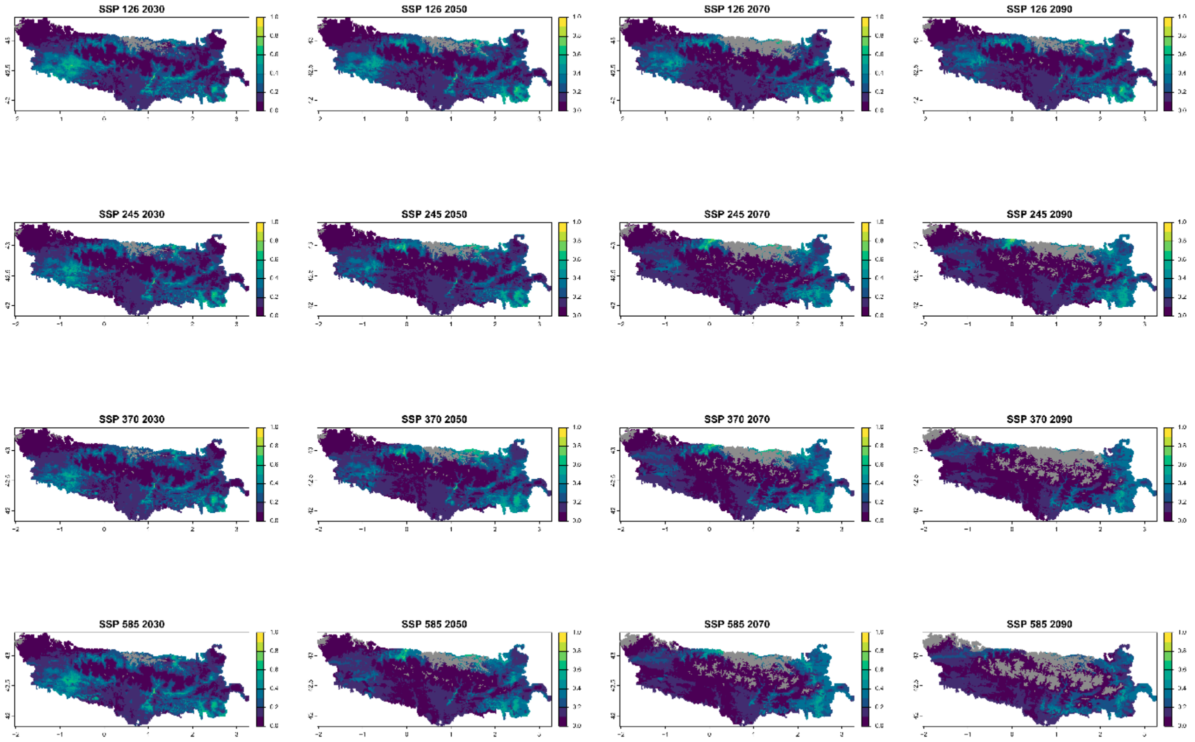


Legend: Non−analogous bioclimatic conditions (MESS < 0)

Current bioclimatic suitability in the Pyrenees for *Armeria bubanii*

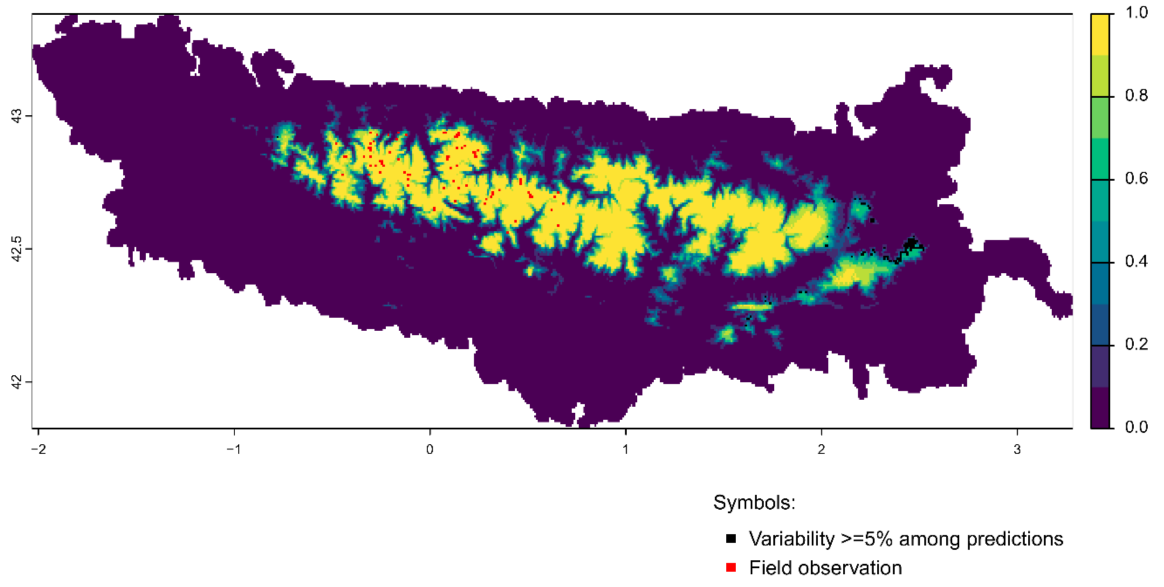


Future bioclimatic suitability in the Pyrenees by 2090 for *Armeria bubanii*

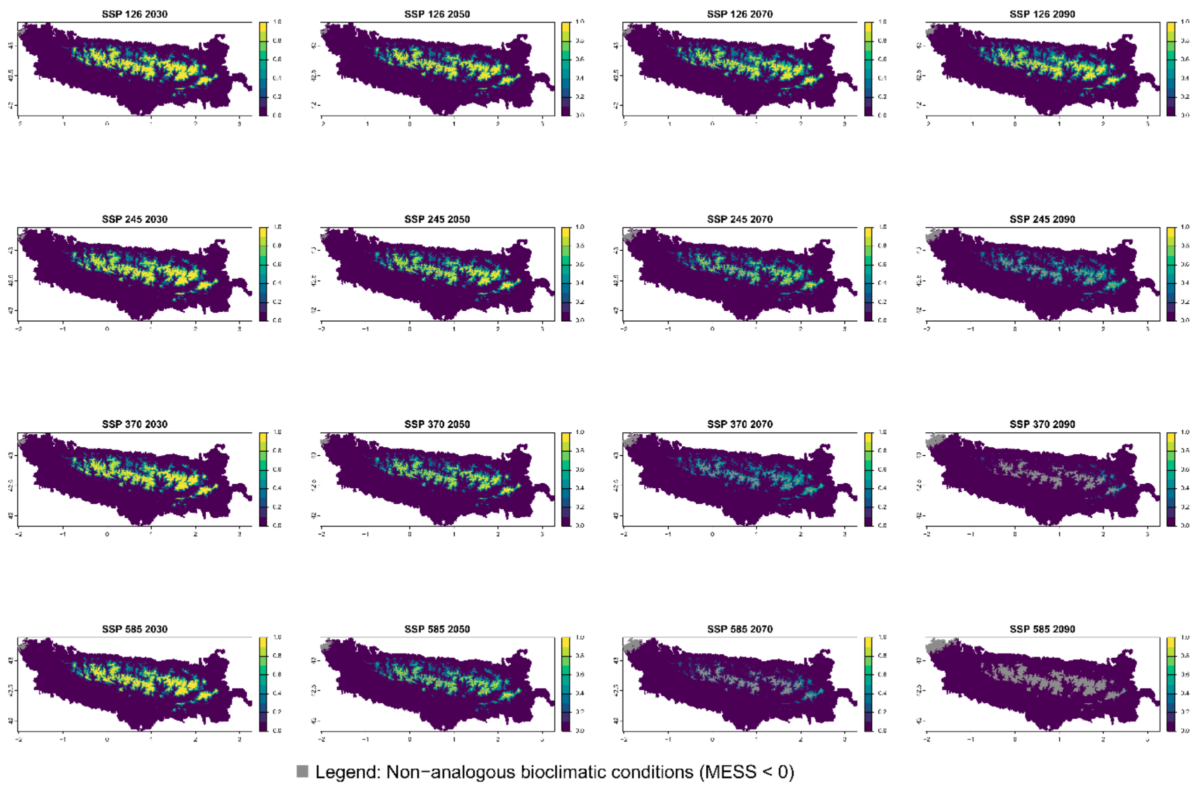

Current bioclimatic suitability in the Pyrenees for *Campanula jaubertiana*

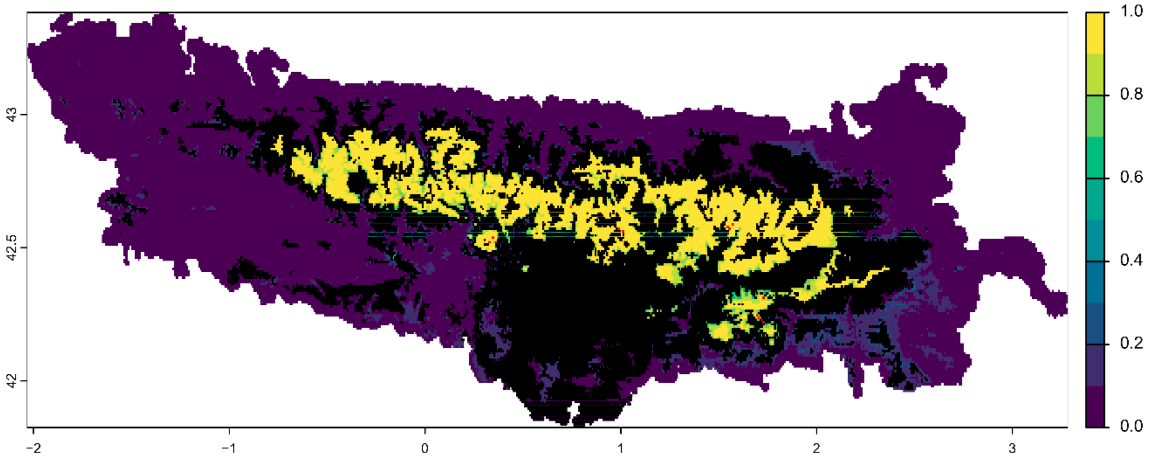


Symbols:

- Variability >=5% among predictions
- Field observation

Future bioclimatic suitability in the Pyrenees by 2090 for *Campanula jaubertiana*

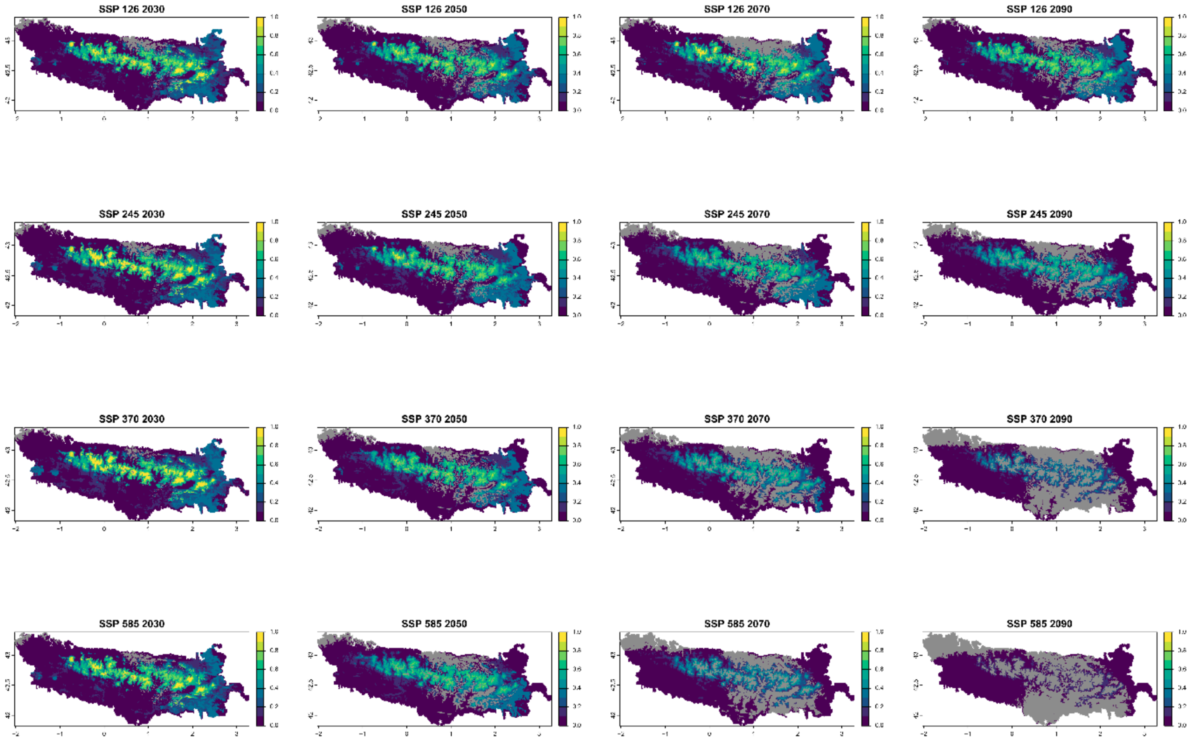


Legend: Non−analogous bioclimatic conditions (MESS < 0)

Current bioclimatic suitability in the Pyrenees for *Campanula precatoria*

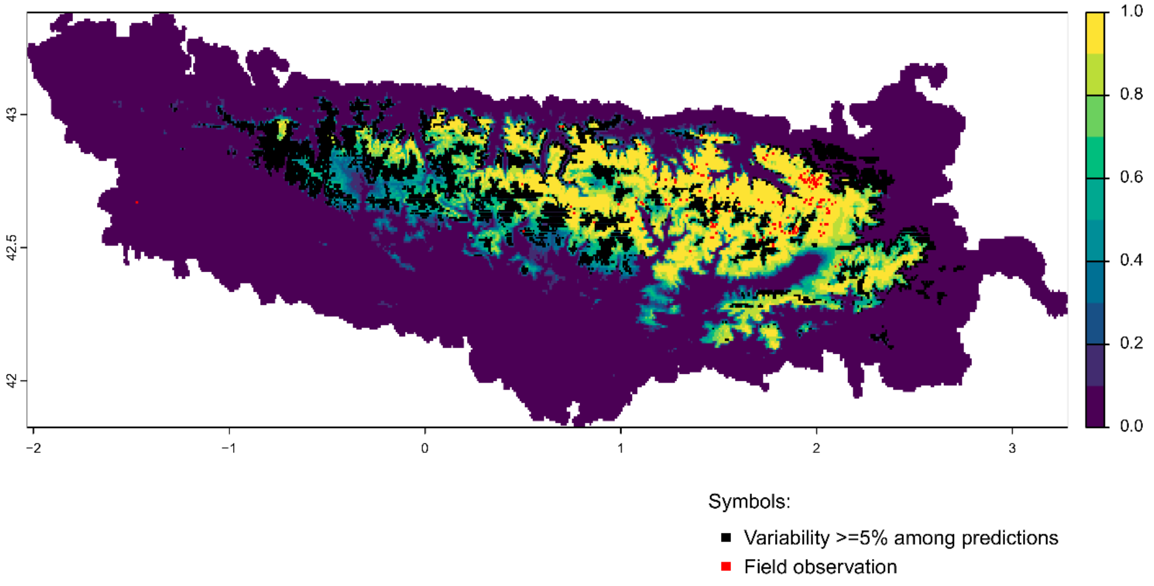


Future bioclimatic suitability in the Pyrenees by 2090 for *Campanula precatoria*

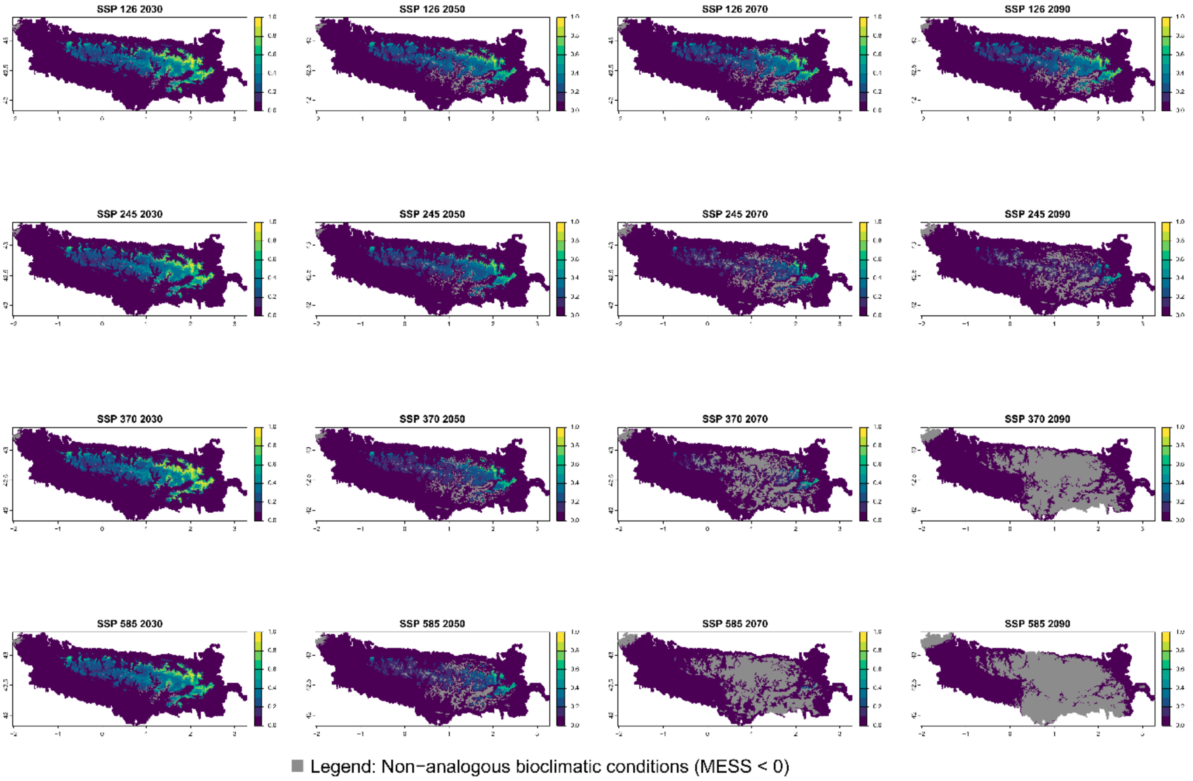

Current bioclimatic suitability in the Pyrenees for *Centaurea costae*

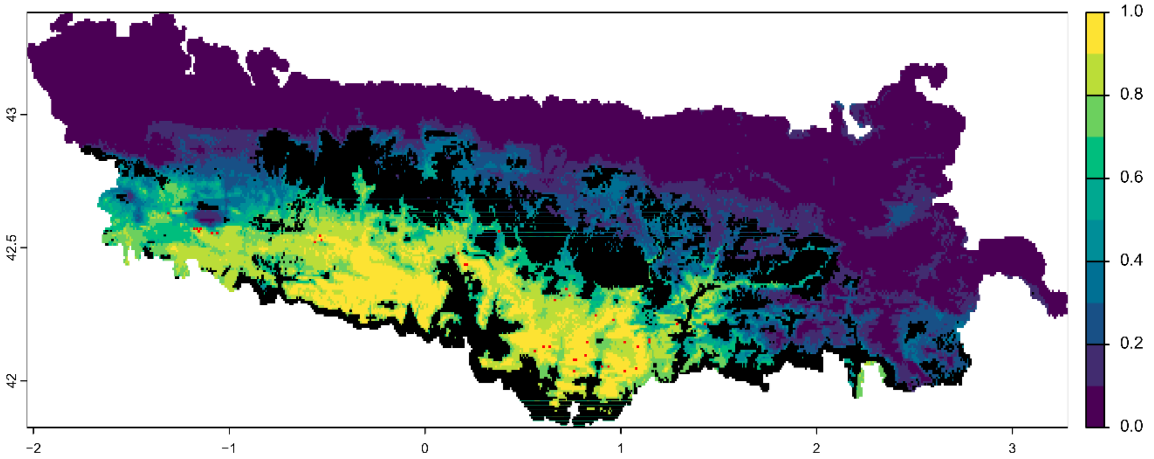


Symbols:

- ■ Variability >=5% among predictions
- ■ Field observation

Future bioclimatic suitability in the Pyrenees by 2090 for *Centaurea costae*

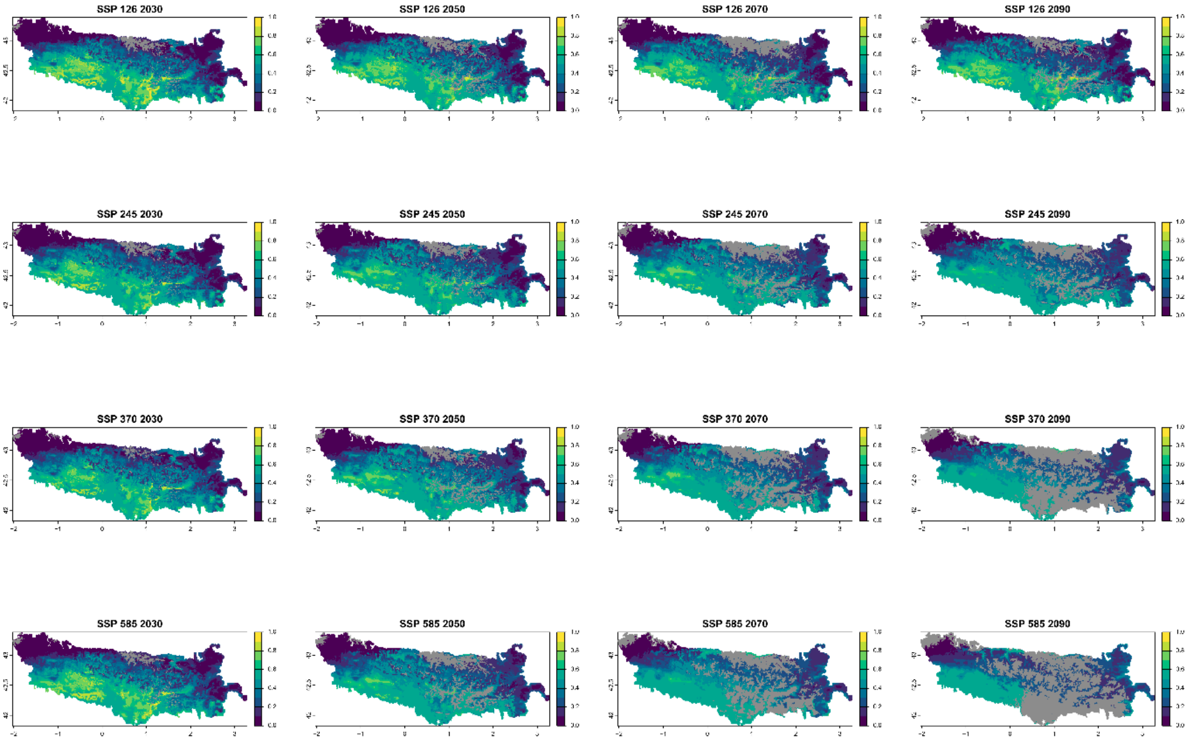


■ Legend: Non−analogous bioclimatic conditions (MESS < 0)

Current bioclimatic suitability in the Pyrenees for *Centaurea emigrantis*

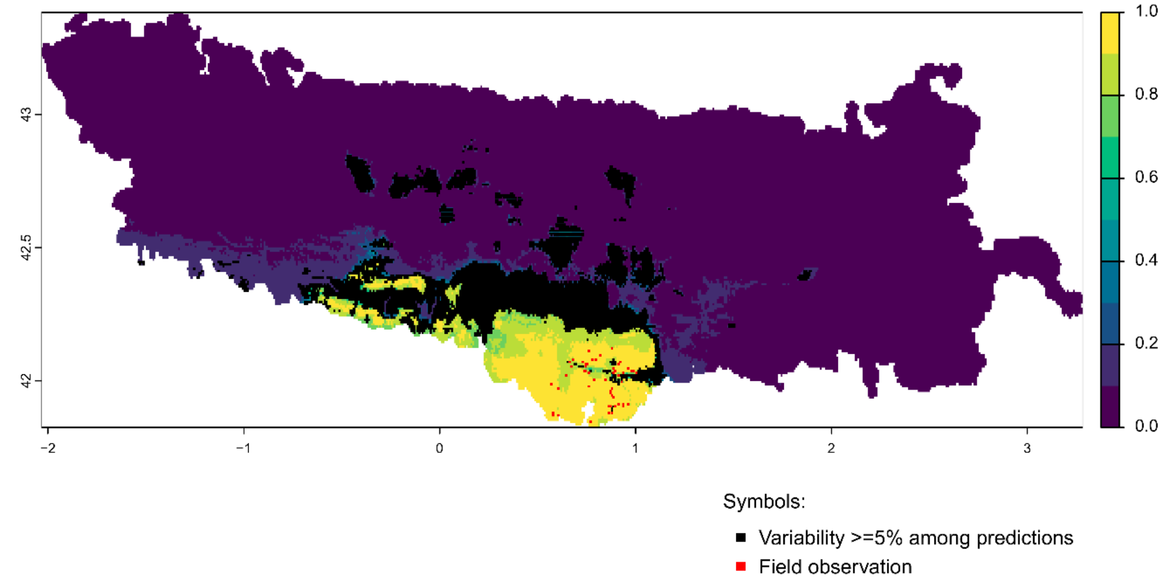


Future bioclimatic suitability in the Pyrenees by 2090 for *Centaurea emigrantis*

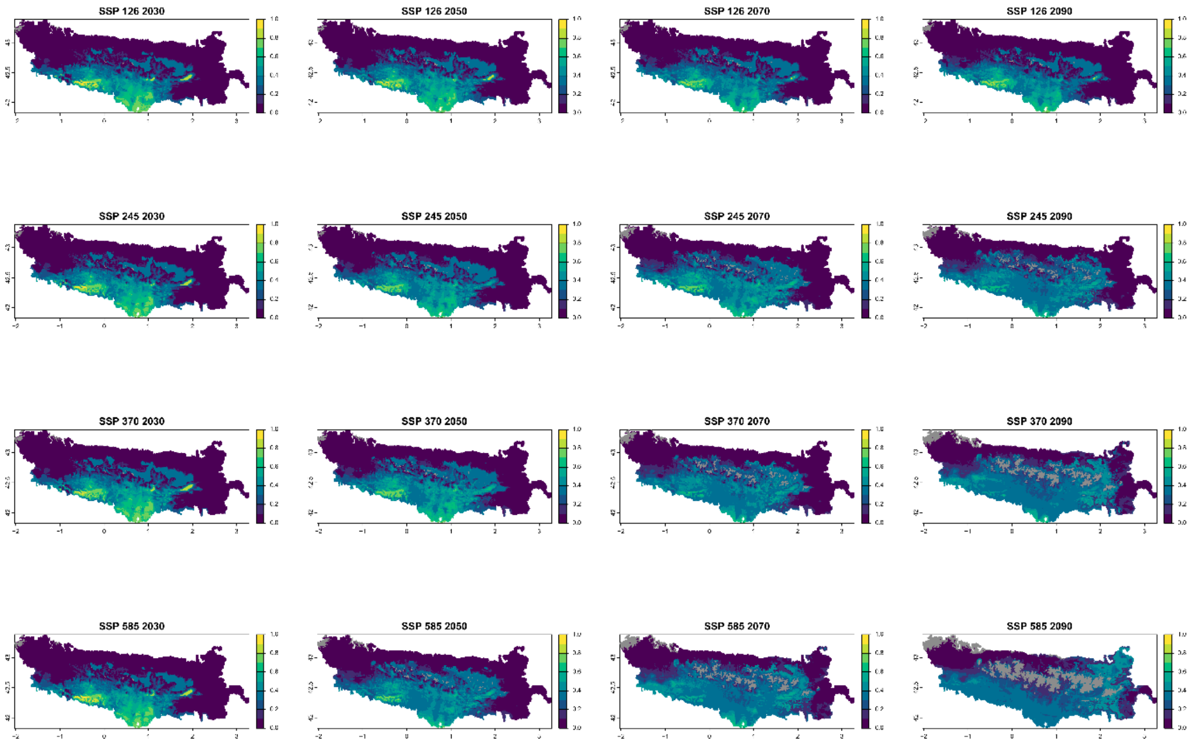


Legend: Non−analogous bioclimatic conditions (MESS < 0)

Current bioclimatic suitability in the Pyrenees for *Cerastium pyrenaicum*

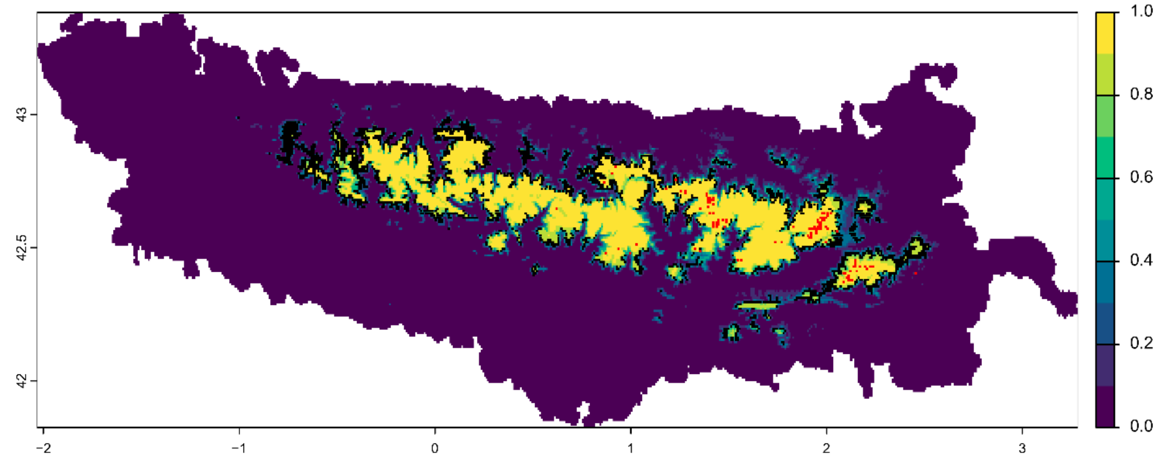


Symbols:

- Variability >=5% among predictions
- Field observation

Future bioclimatic suitability in the Pyrenees by 2090 for *Cerastium pyrenaicum*

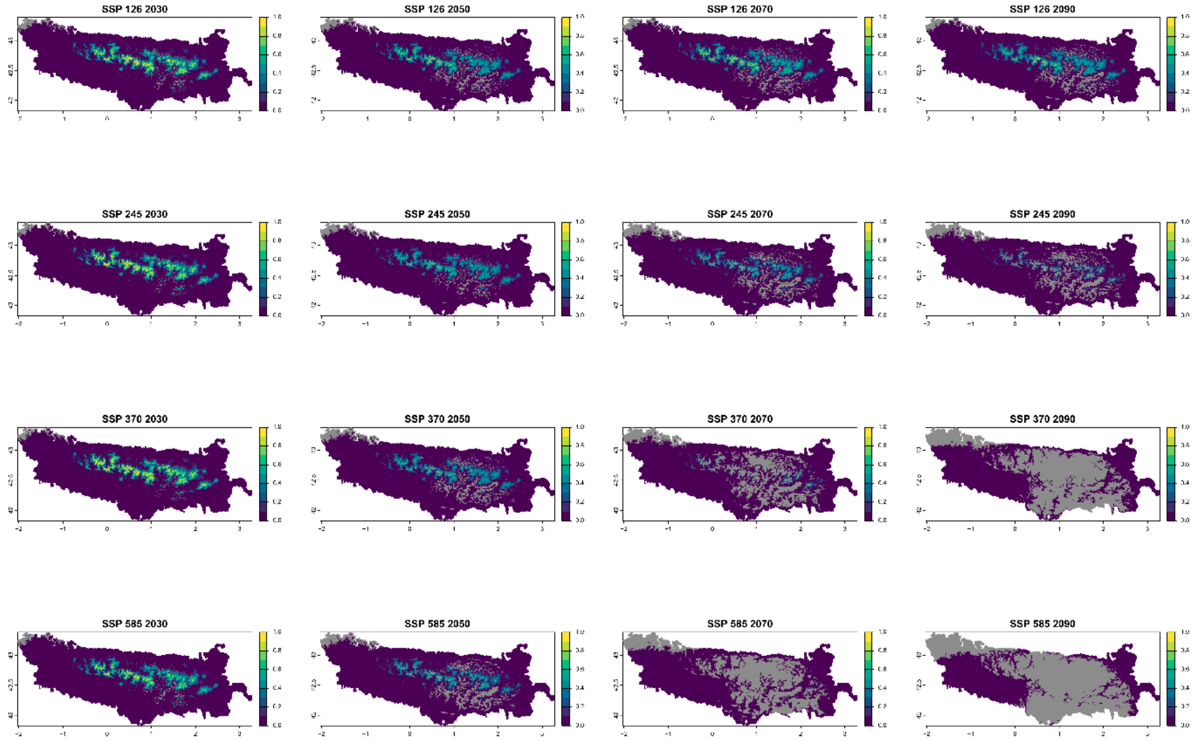


Legend: Non−analogous bioclimatic conditions (MESS < 0)

Current bioclimatic suitability in the Pyrenees for *Cirsium glabrum*

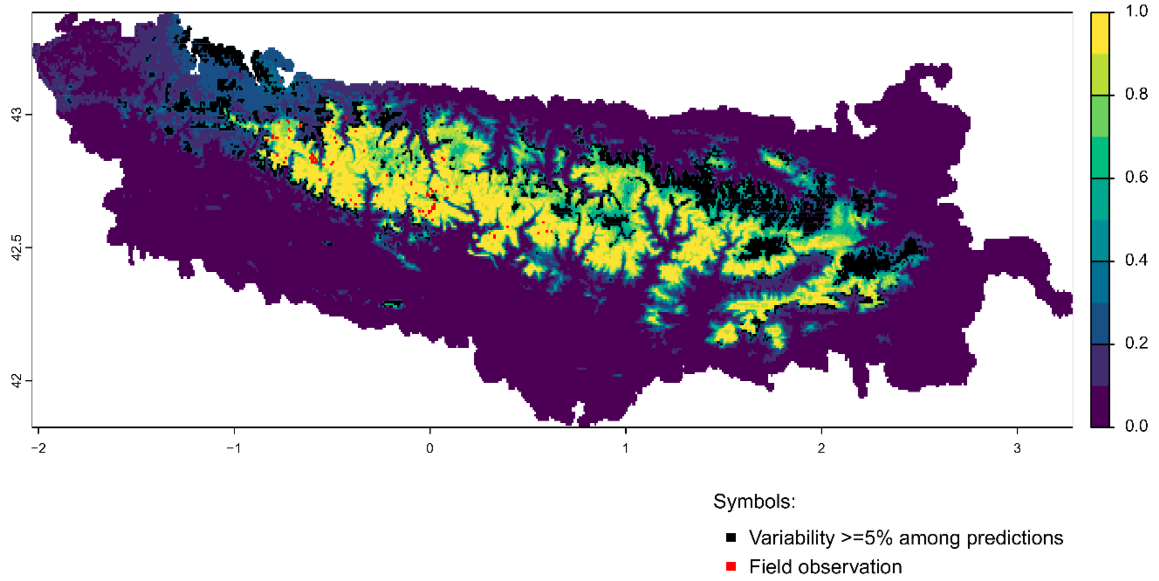


Future bioclimatic suitability in the Pyrenees by 2090 for *Cirsium glabrum*

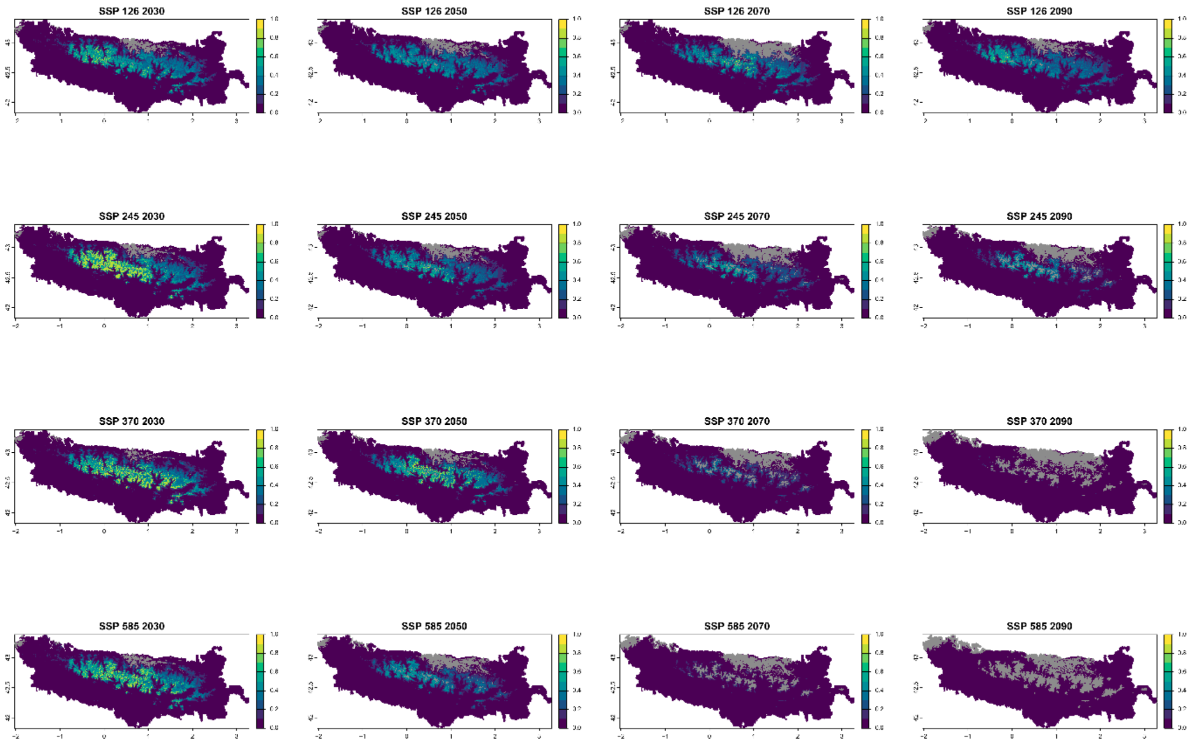


Legend: Non−analogous bioclimatic conditions (MESS < 0)

### Current bioclimatic suitability in the Pyrenees for *Delphinium montanum*

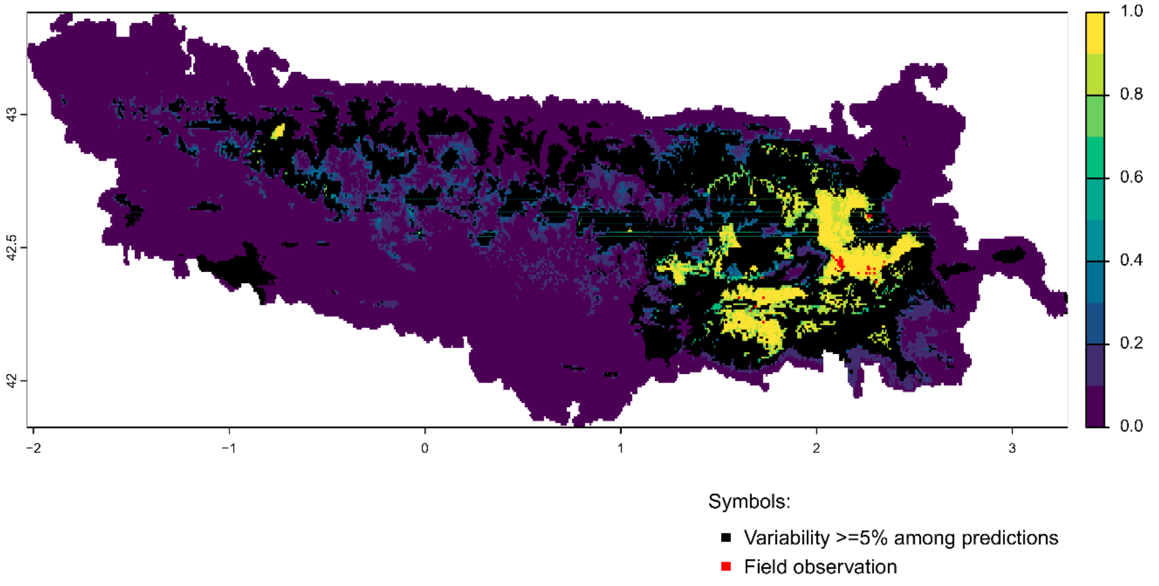


Future bioclimatic suitability in the Pyrenees by 2090 for *Delphinium montanum*

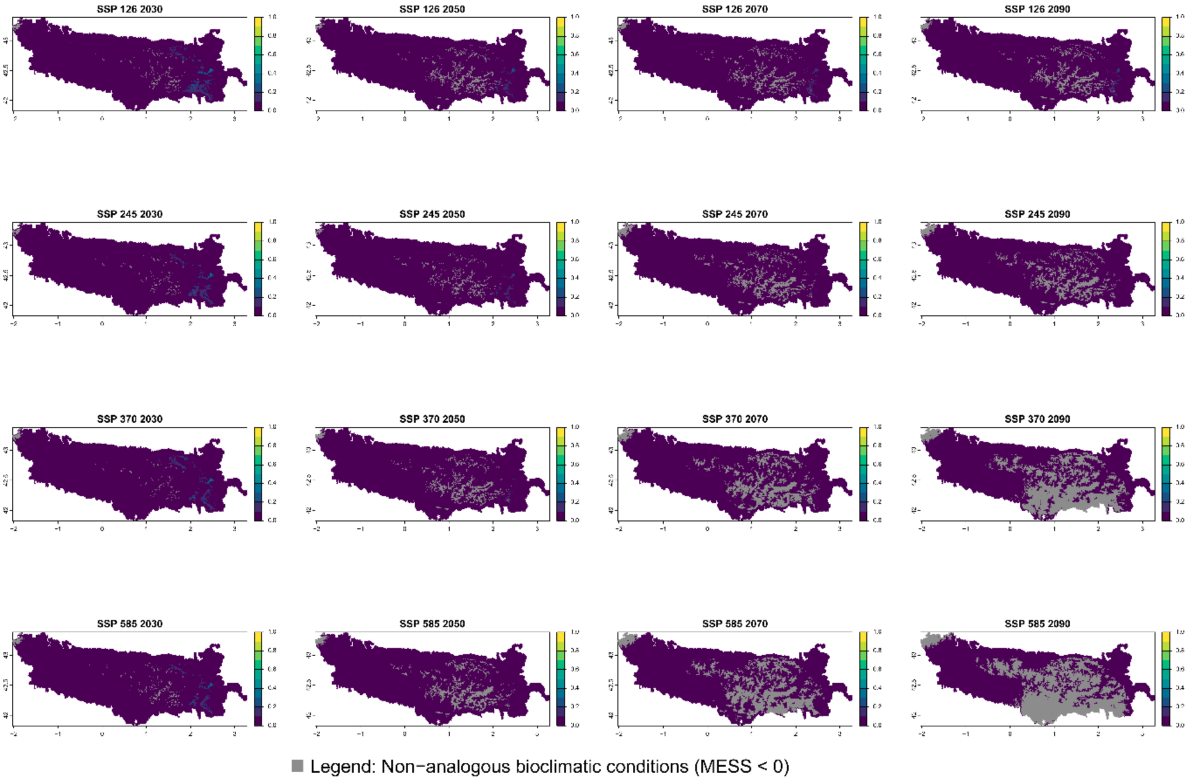

### Current bioclimatic suitability in the Pyrenees for *Dianthus benearnensis*

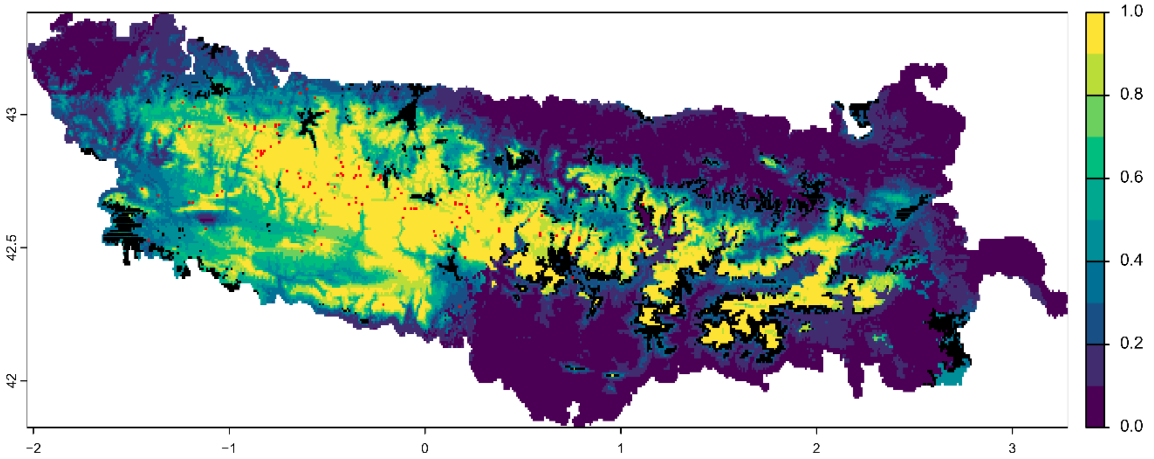


Symbols:

- ■ Variability >=5% among predictions
- ■ Field observation

Future bioclimatic suitability in the Pyrenees by 2090 for *Dianthus benearnensis*

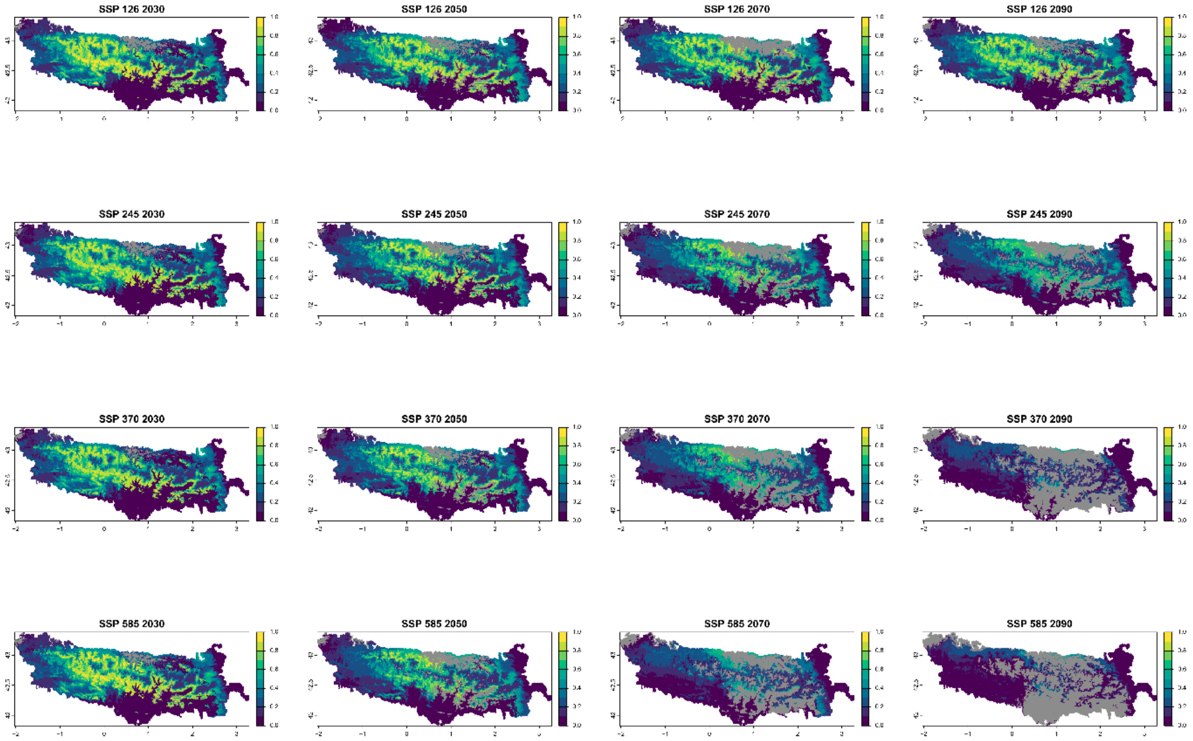


■ Legend: Non−analogous bioclimatic conditions (MESS < 0)

Current bioclimatic suitability in the Pyrenees for *Draba subnivalis*

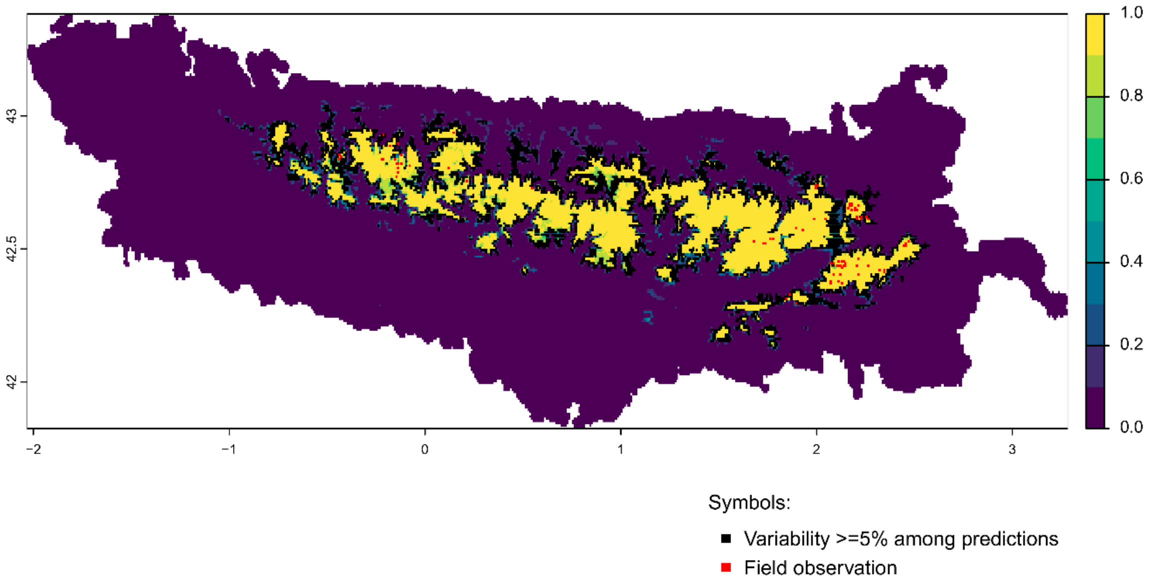


Future bioclimatic suitability in the Pyrenees by 2090 for *Draba subnivalis*

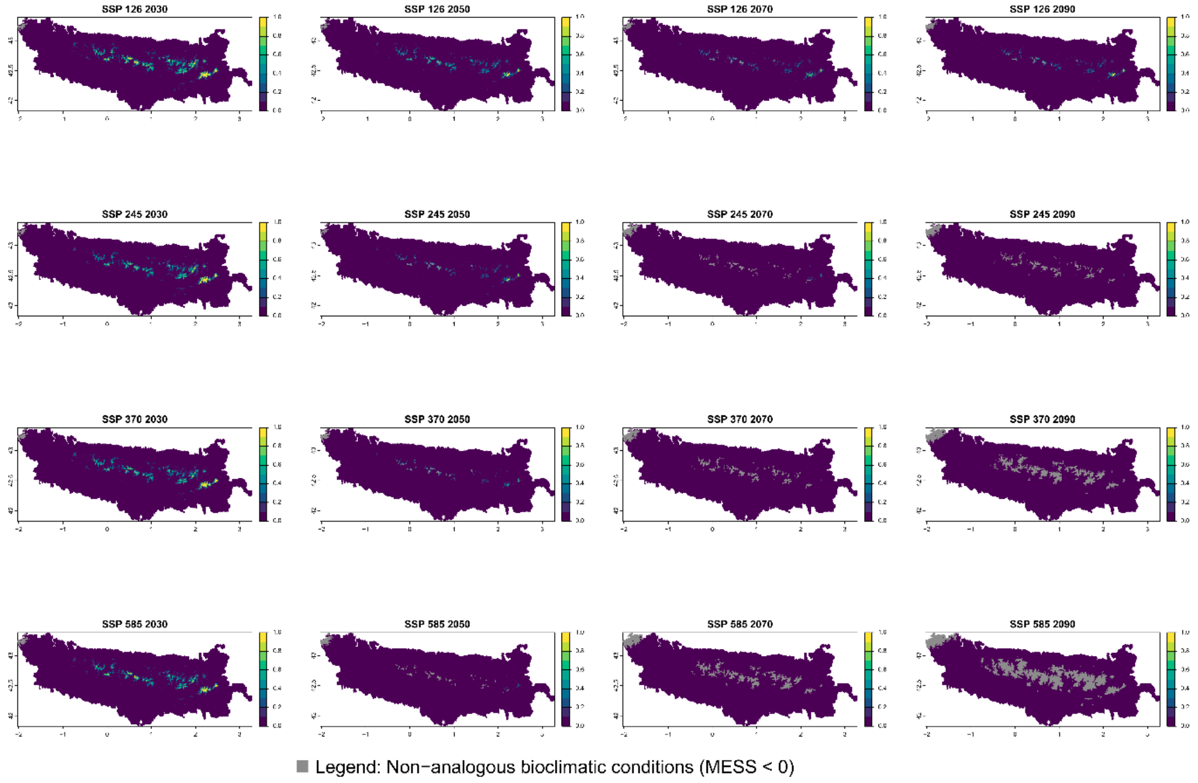

Current bioclimatic suitability in the Pyrenees for *Endressia pyrenaica*

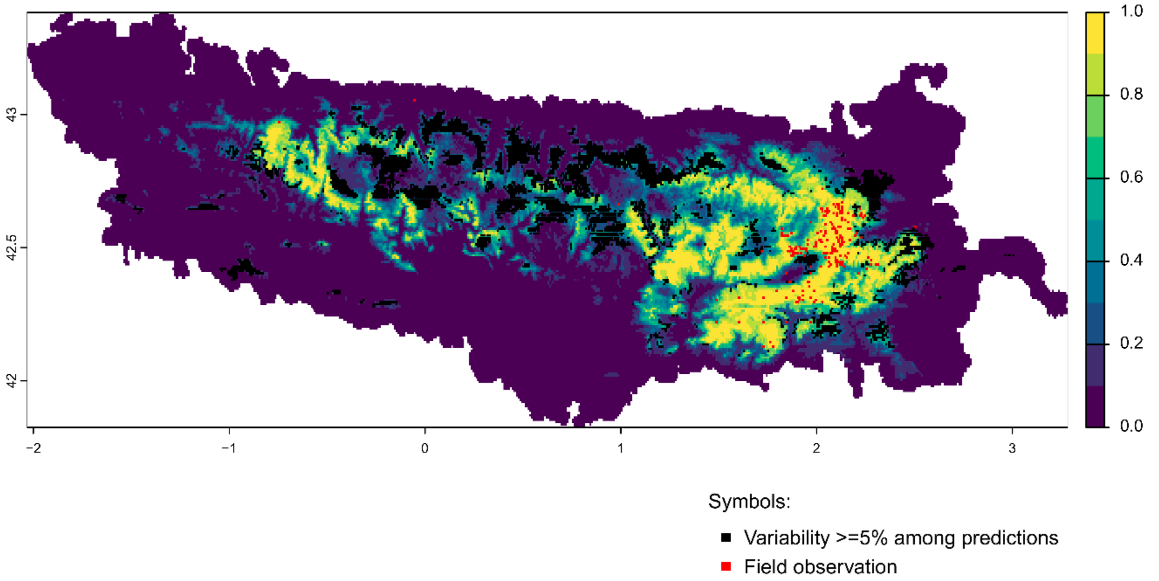


Future bioclimatic suitability in the Pyrenees by 2090 for *Endressia pyrenaica*

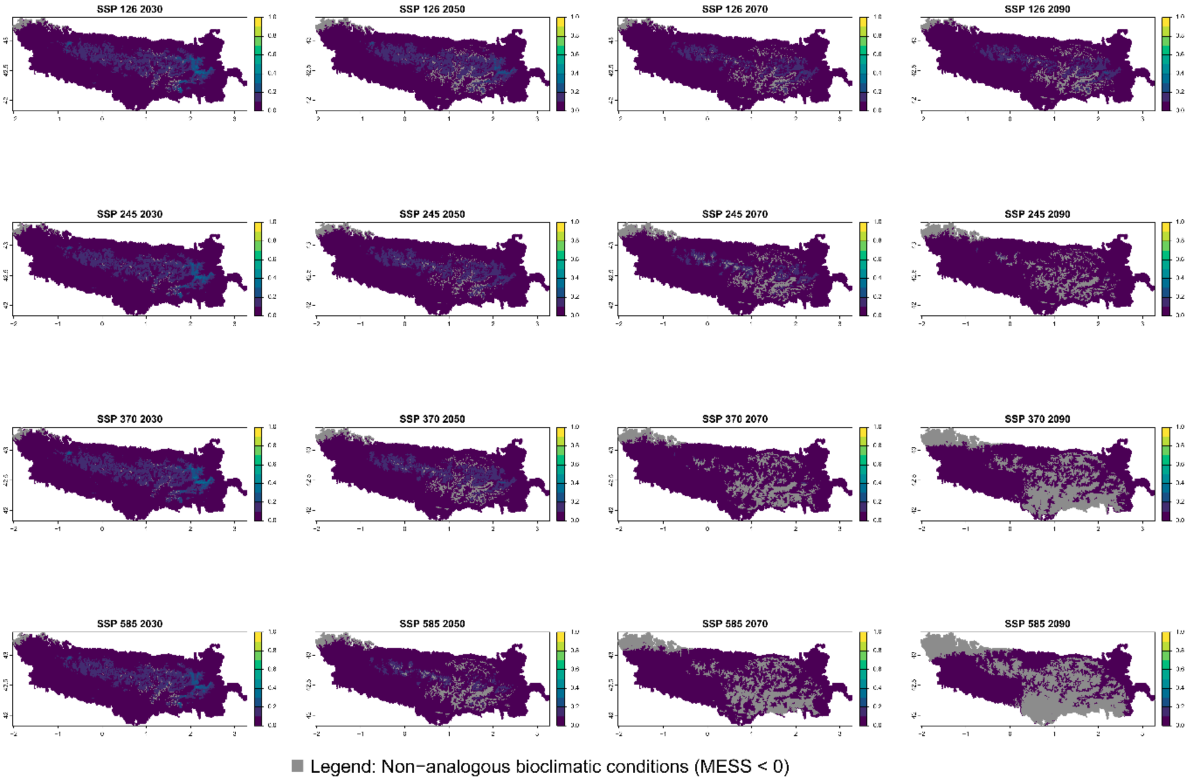

Current bioclimatic suitability in the Pyrenees for *Erodium lucidum*

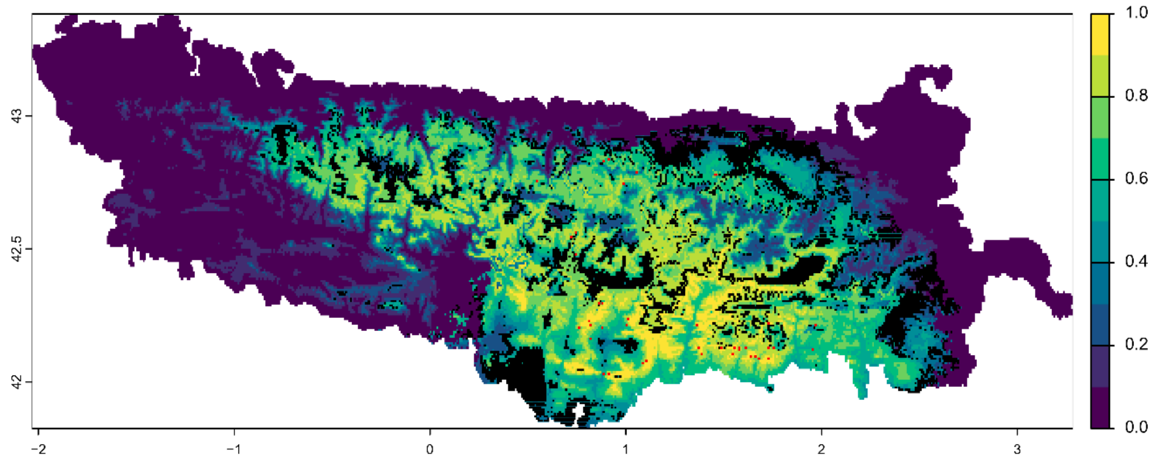


Symbols:

■ Variability >=5% among predictions

■ Field observation

Future bioclimatic suitability in the Pyrenees by 2090 for *Erodium lucidum*

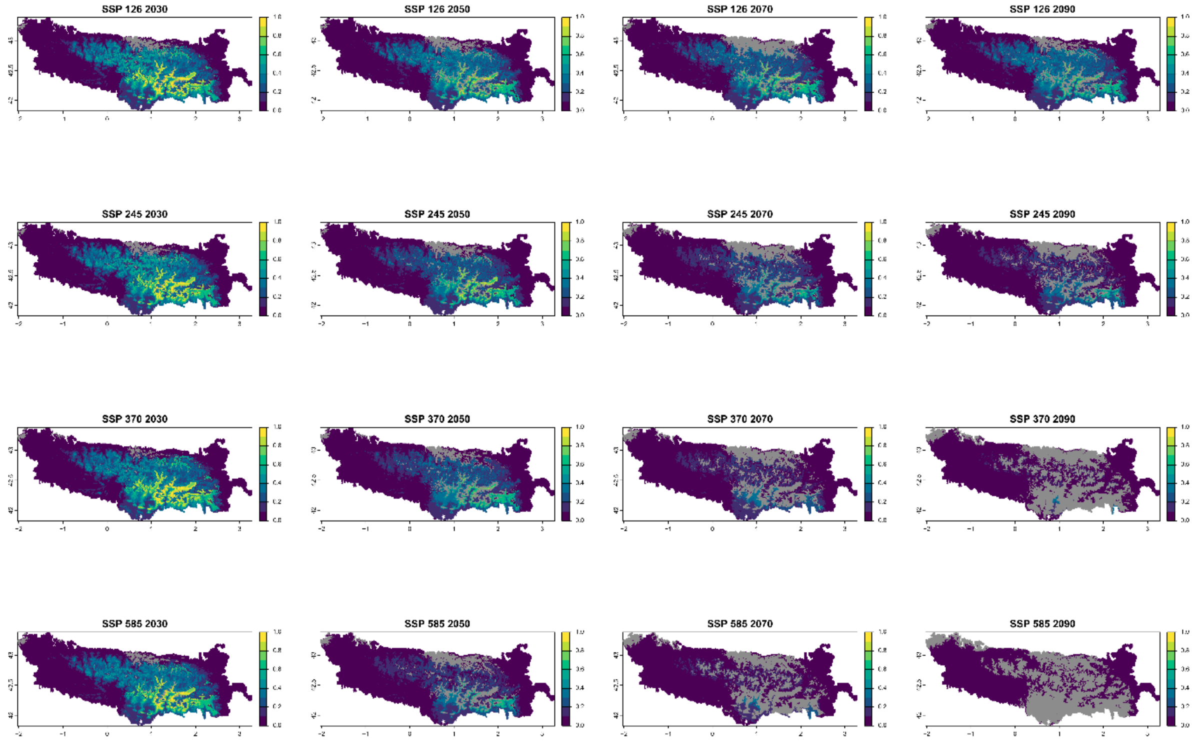


■ Legend: Non−analogous bioclimatic conditions (MESS < 0)

Current bioclimatic suitability in the Pyrenees for *Festuca altopyrenaica*

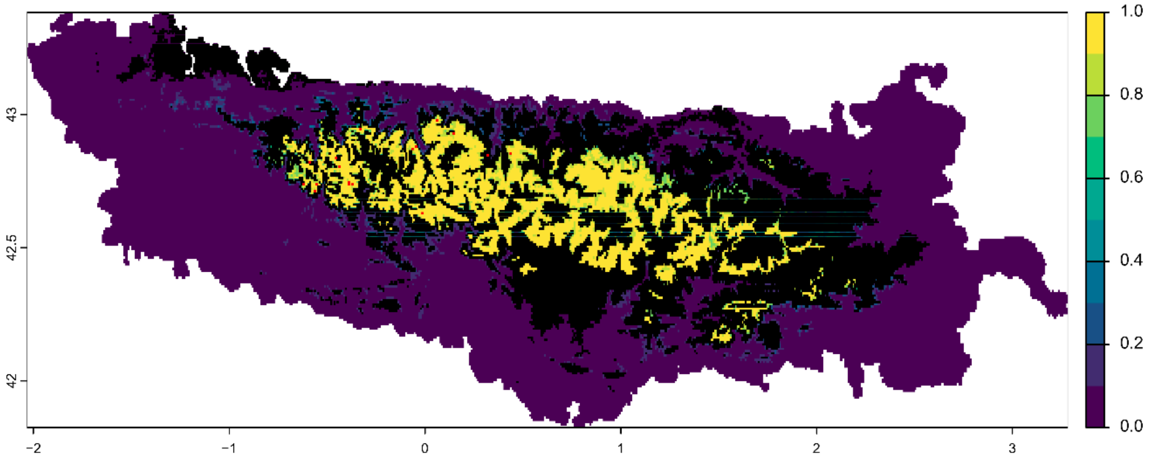


Symbols:

■ Variability >=5% among predictions

■ Field observation

Future bioclimatic suitability in the Pyrenees by 2090 for *Festuca altopyrenaica*

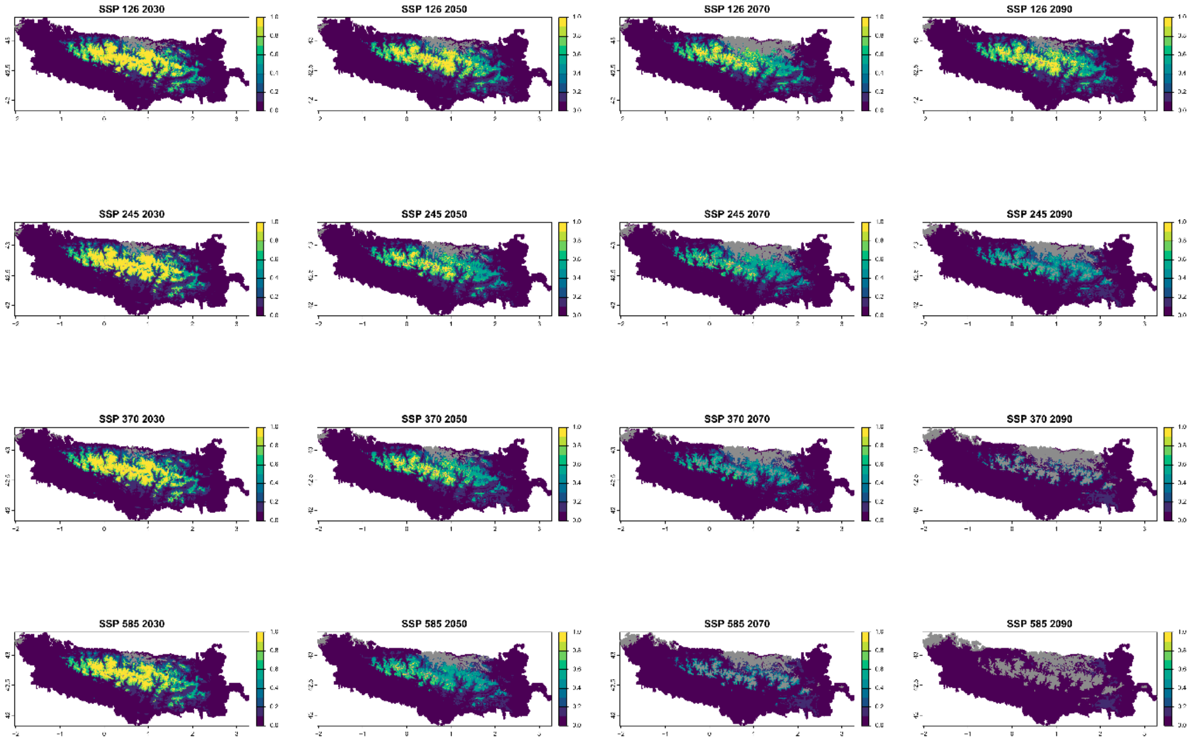


■ Legend: Non−analogous bioclimatic conditions (MESS < 0)

Current bioclimatic suitability in the Pyrenees for *Festuca borderei*

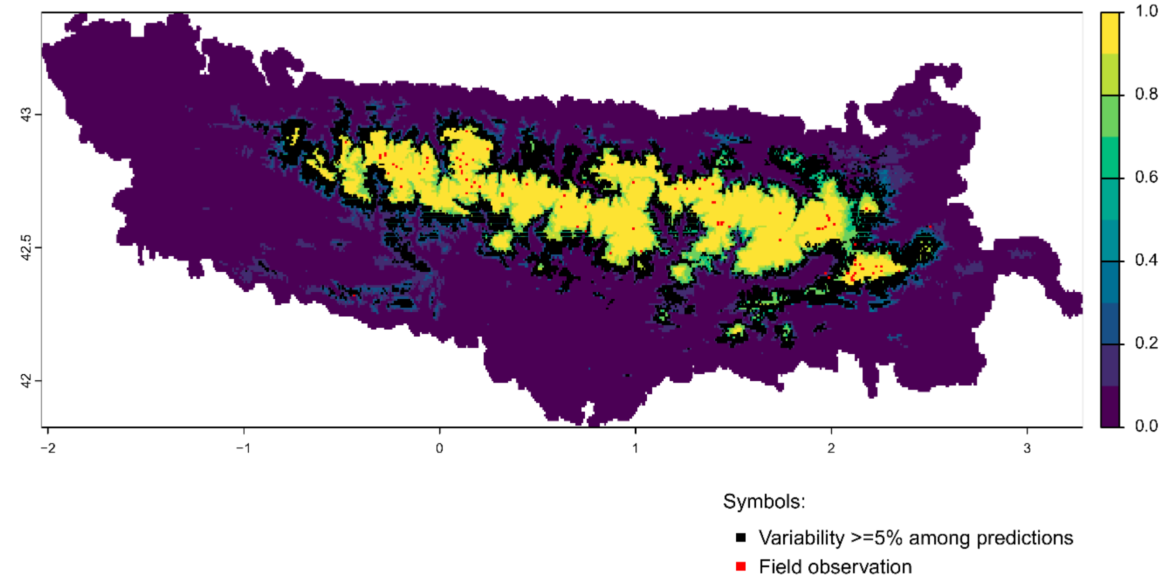


Future bioclimatic suitability in the Pyrenees by 2090 for *Festuca borderei*

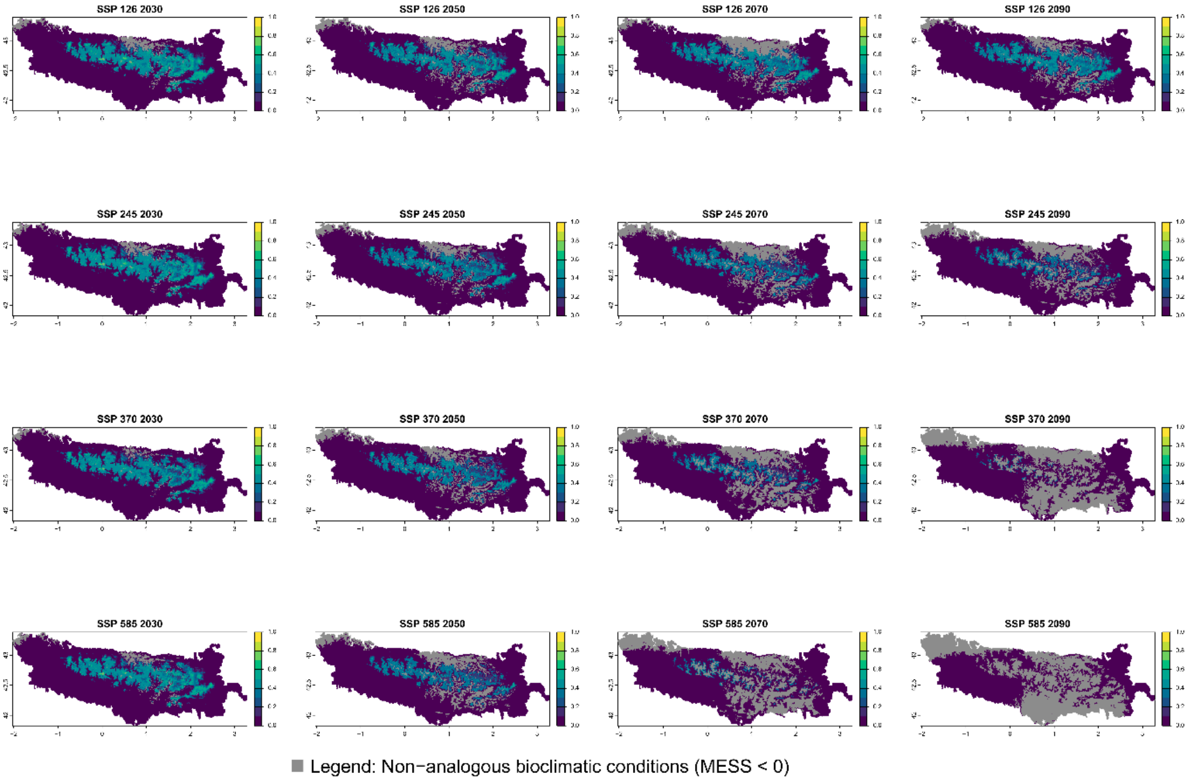

Current bioclimatic suitability in the Pyrenees for *Festuca pyrenaica*

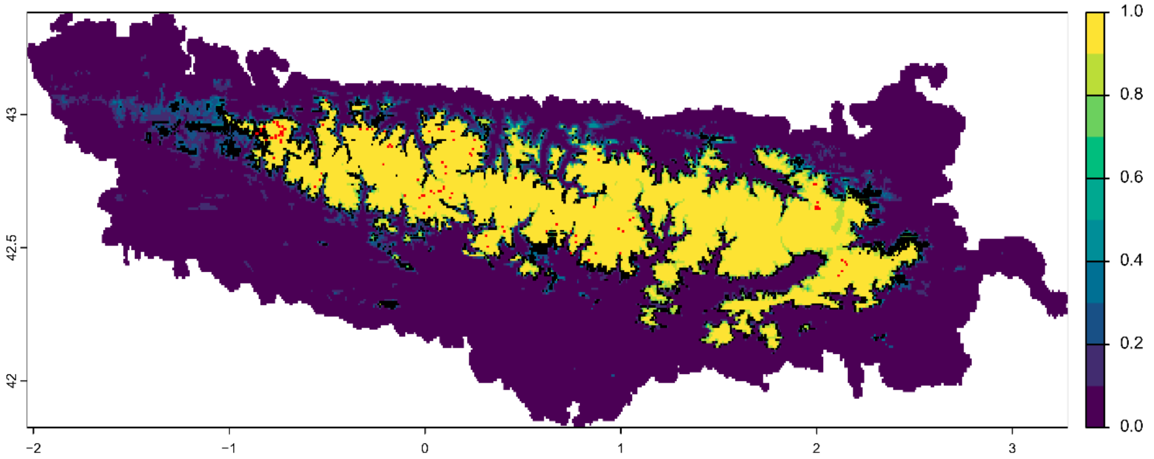


Symbols:

■ Variability >=5% among predictions

■ Field observation

Future bioclimatic suitability in the Pyrenees by 2090 for *Festuca pyrenaica*

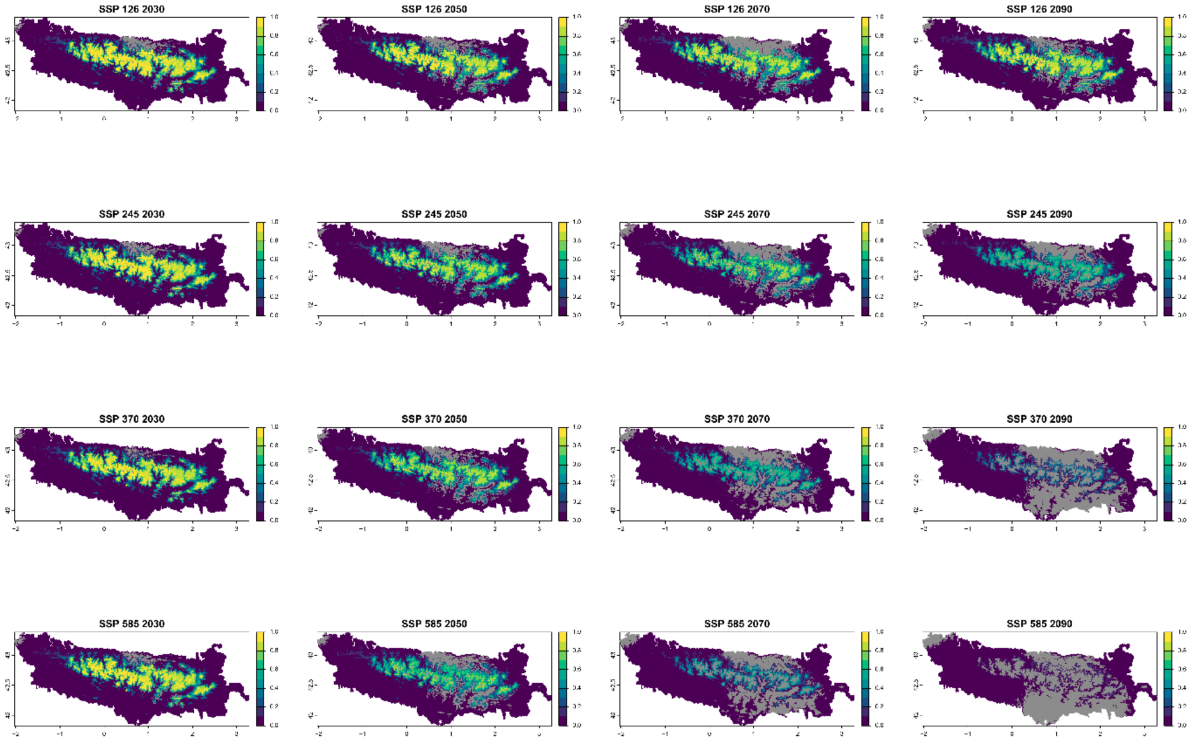


■ Legend: Non−analogous bioclimatic conditions (MESS < 0)

### Current bioclimatic suitability in the Pyrenees for *Galeopsis pyrenaica*

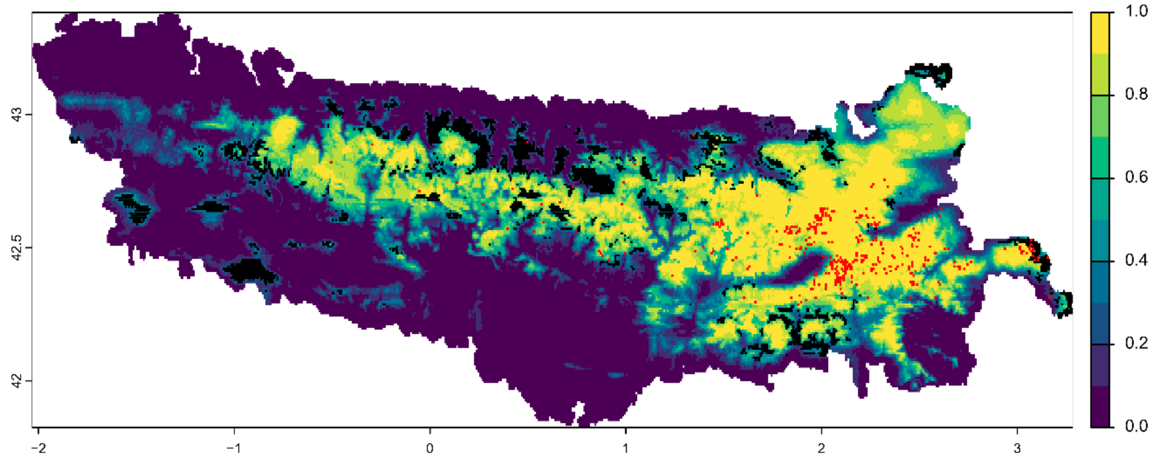


Symbols:

- Variability >=5% among predictions
- Field observation

Future bioclimatic suitability in the Pyrenees by 2090 for *Galeopsis pyrenaica*

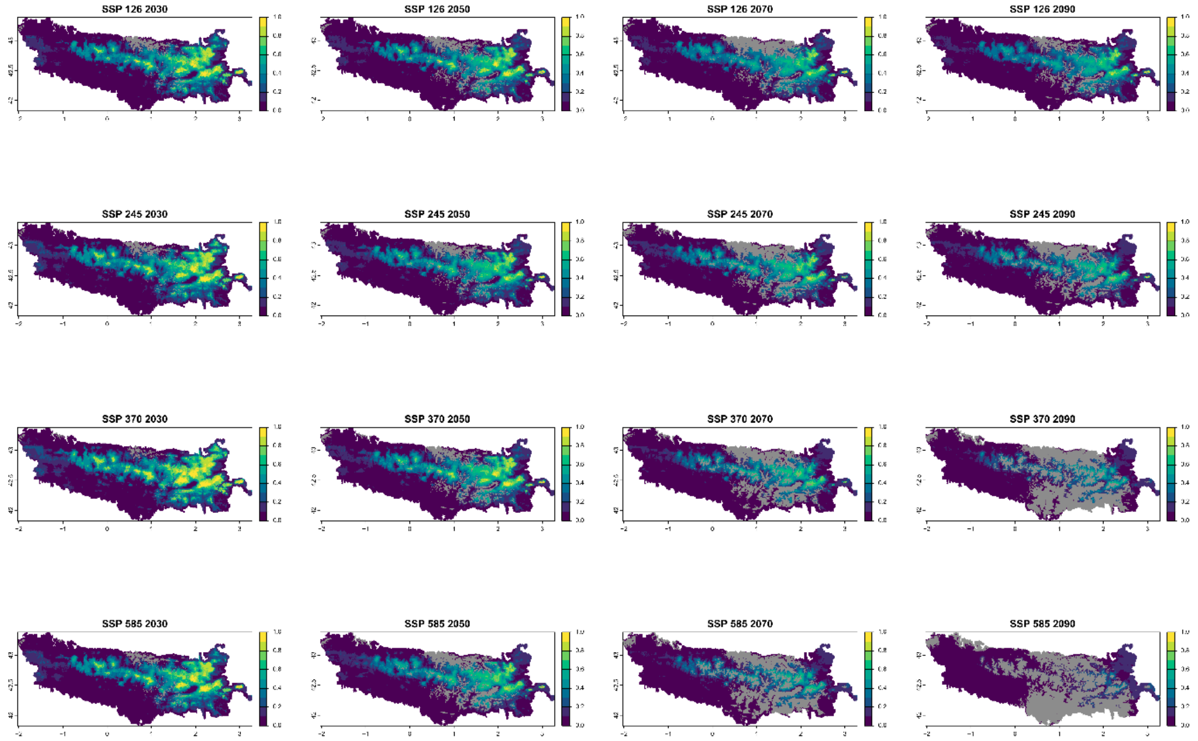


Legend: Non−analogous bioclimatic conditions (MESS < 0)

Current bioclimatic suitability in the Pyrenees for *Galium cespitosum*

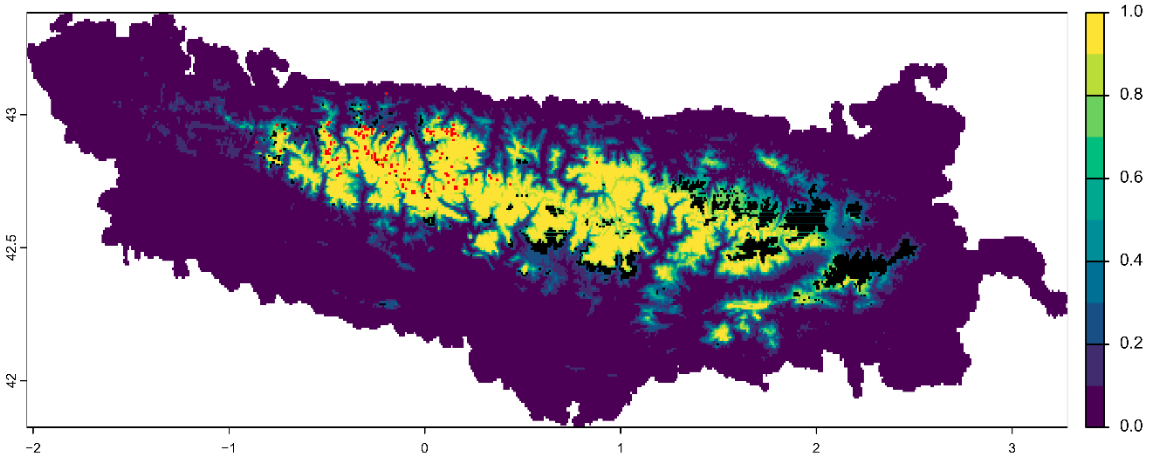


Symbols:

■ Variability >=5% among predictions

■ Field observation

Future bioclimatic suitability in the Pyrenees by 2090 for *Galium cespitosum*

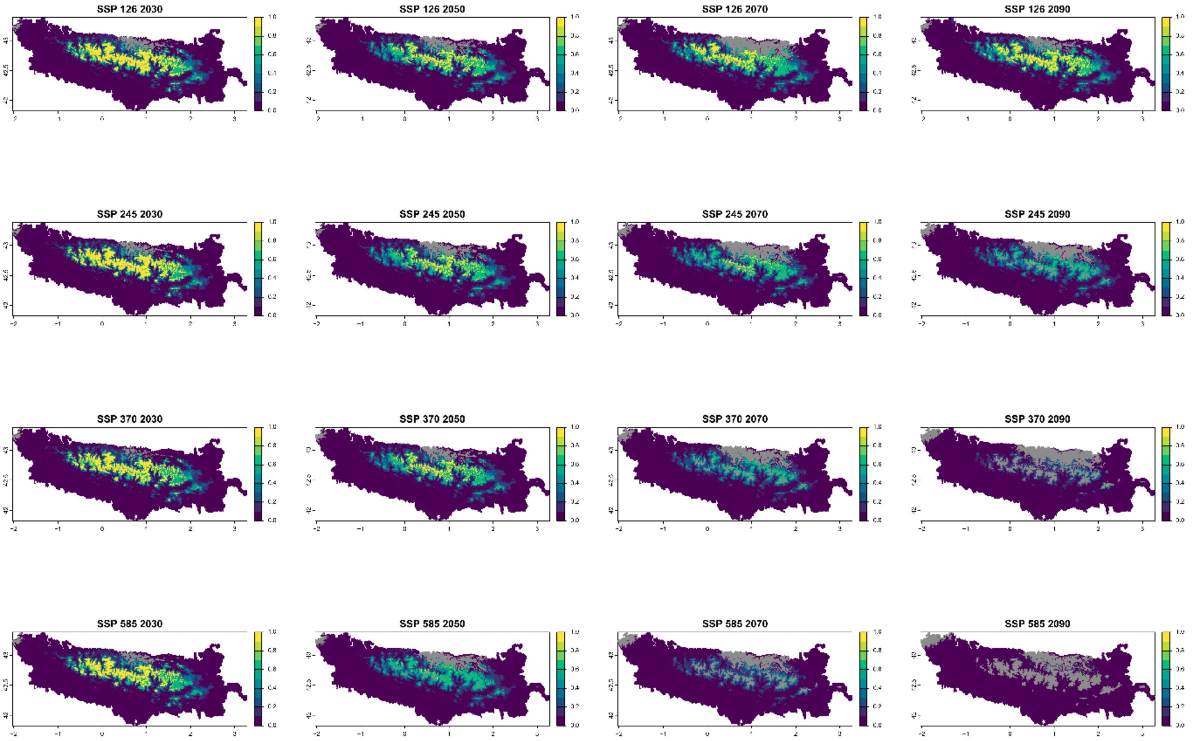


■ Legend: Non−analogous bioclimatic conditions (MESS < 0)

Current bioclimatic suitability in the Pyrenees for *Iberis bernardiana*

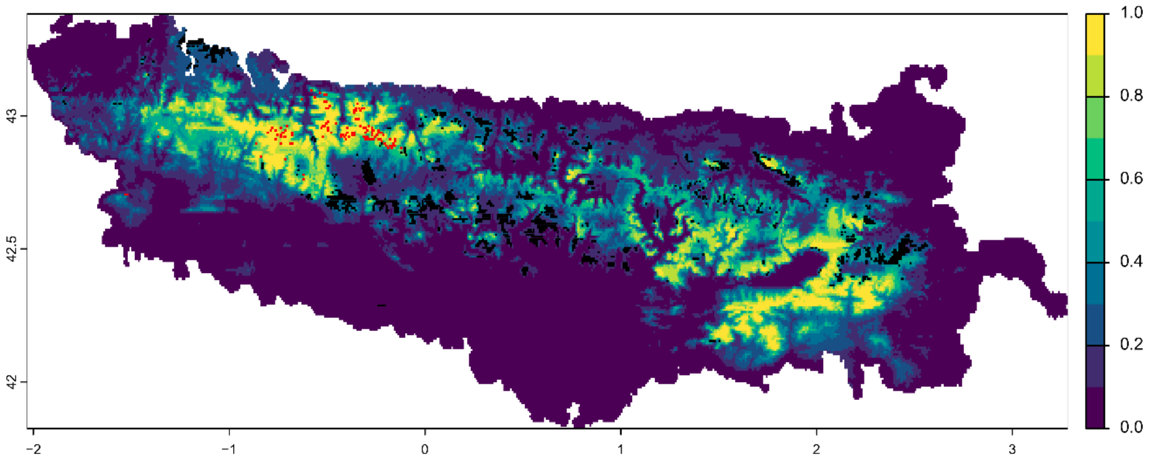


Symbols:

- Variability >=5% among predictions
- Field observation

Future bioclimatic suitability in the Pyrenees by 2090 for *Iberis bernardiana*

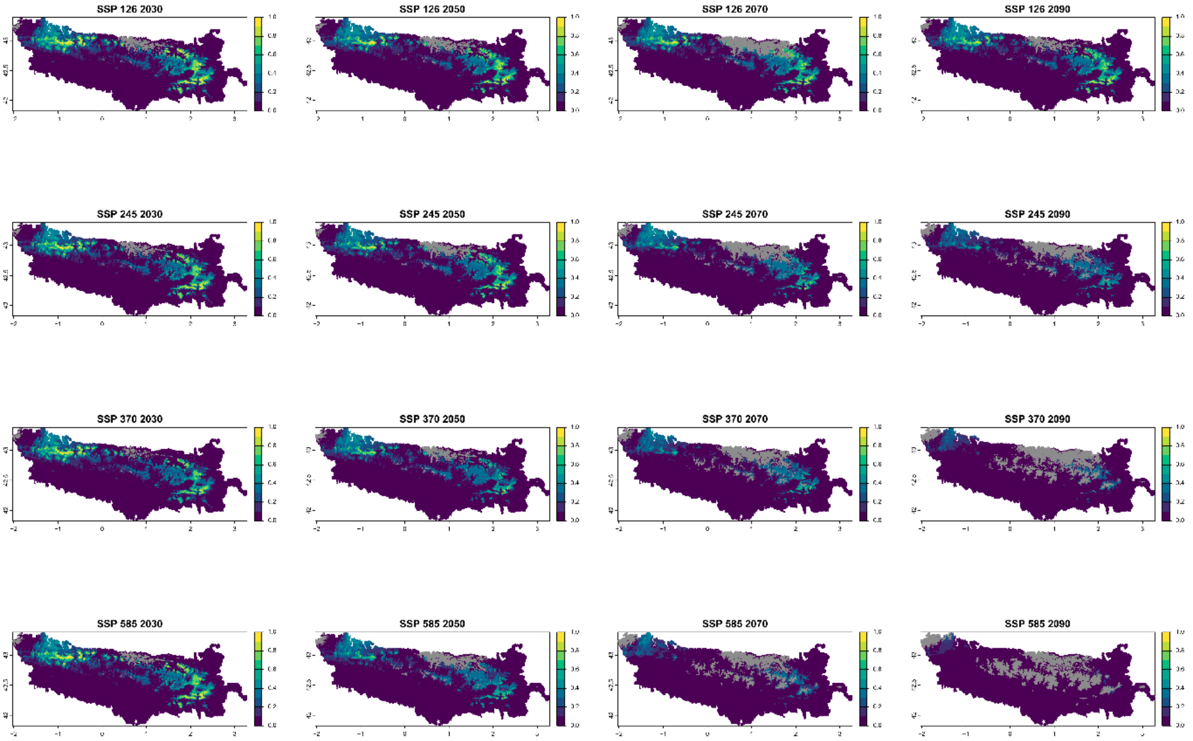


Legend: Non−analogous bioclimatic conditions (MESS < 0)

### Current bioclimatic suitability in the Pyrenees for *Iberis spathulata*

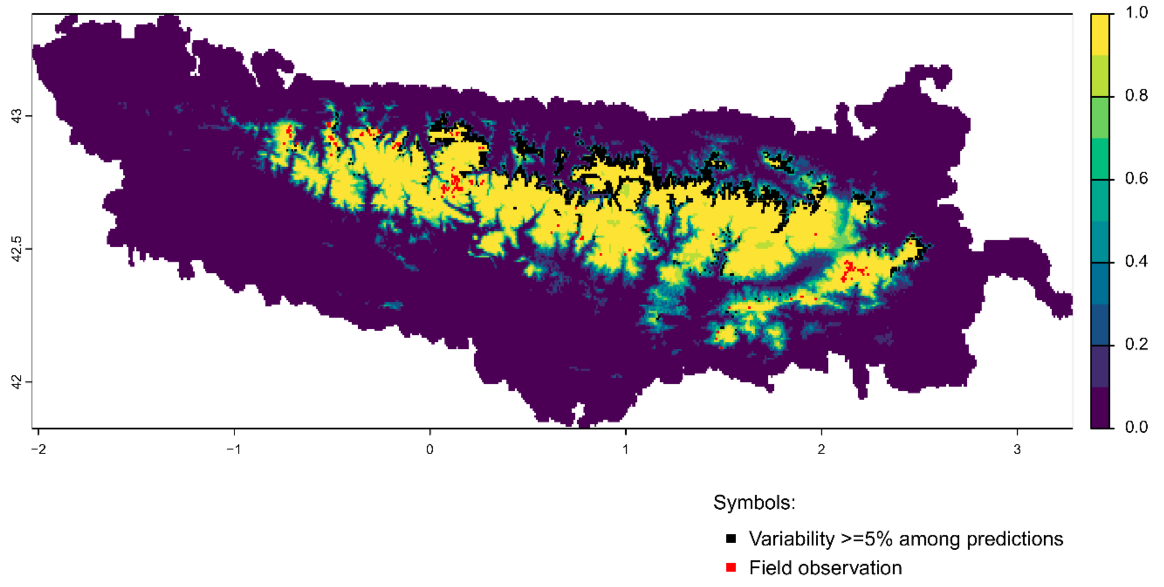


Future bioclimatic suitability in the Pyrenees by 2090 for *Iberis spathulata*

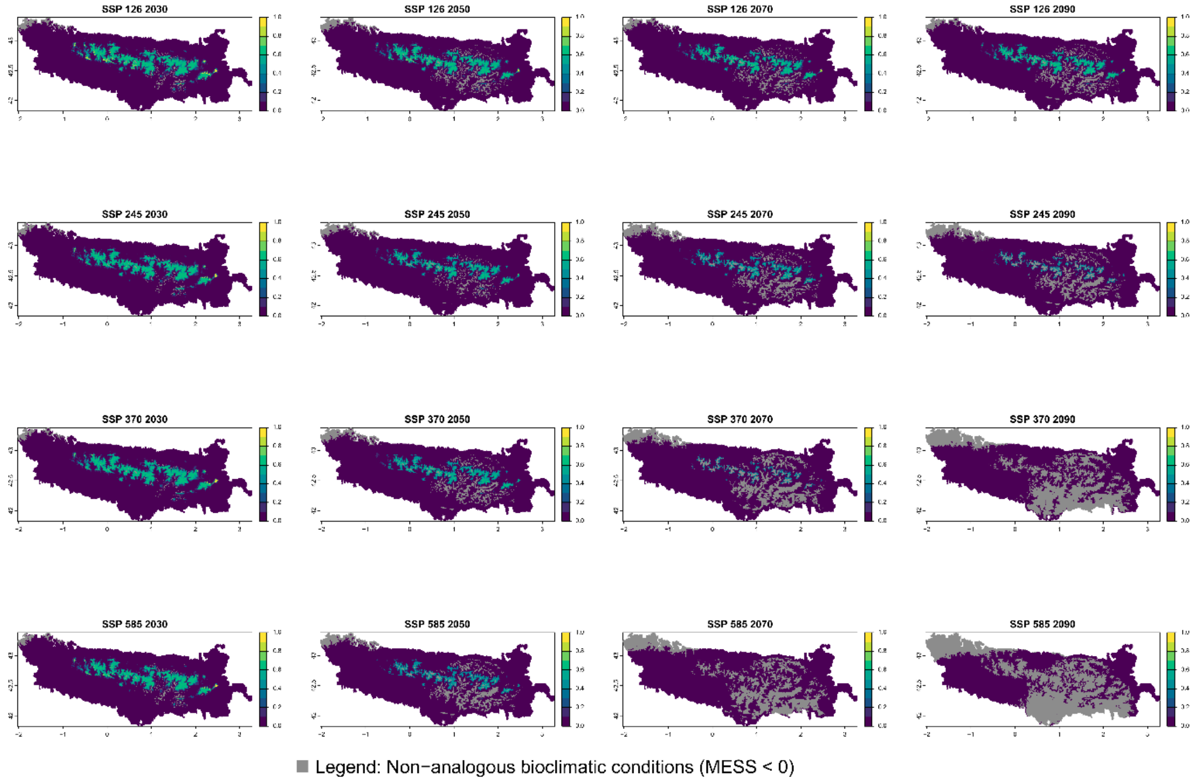

Current bioclimatic suitability in the Pyrenees for *Knautia lebrunii*

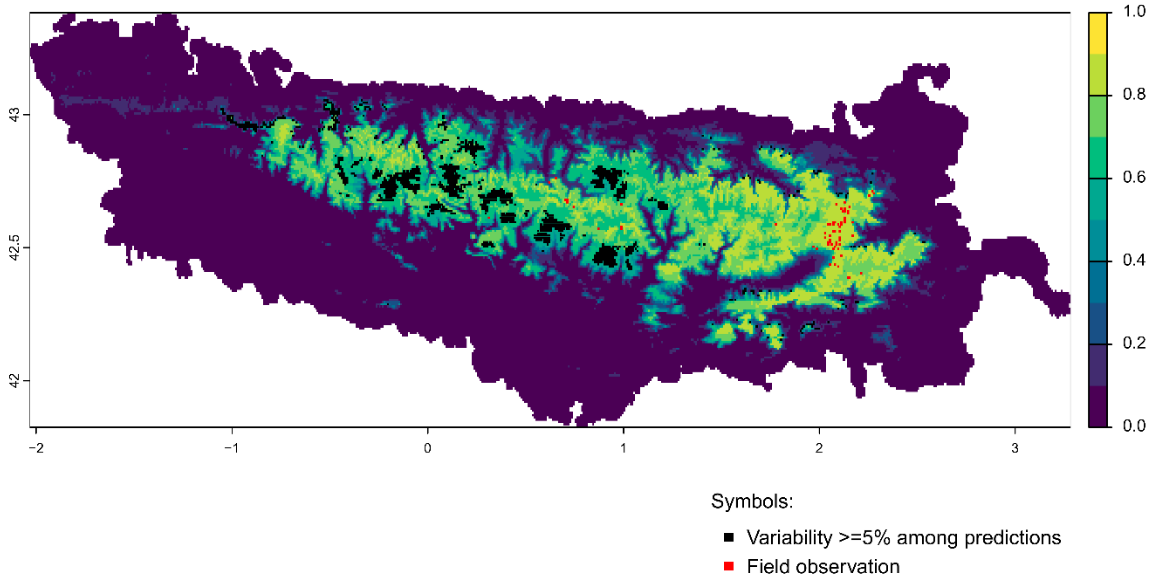


Future bioclimatic suitability in the Pyrenees by 2090 for *Knautia lebrunii*

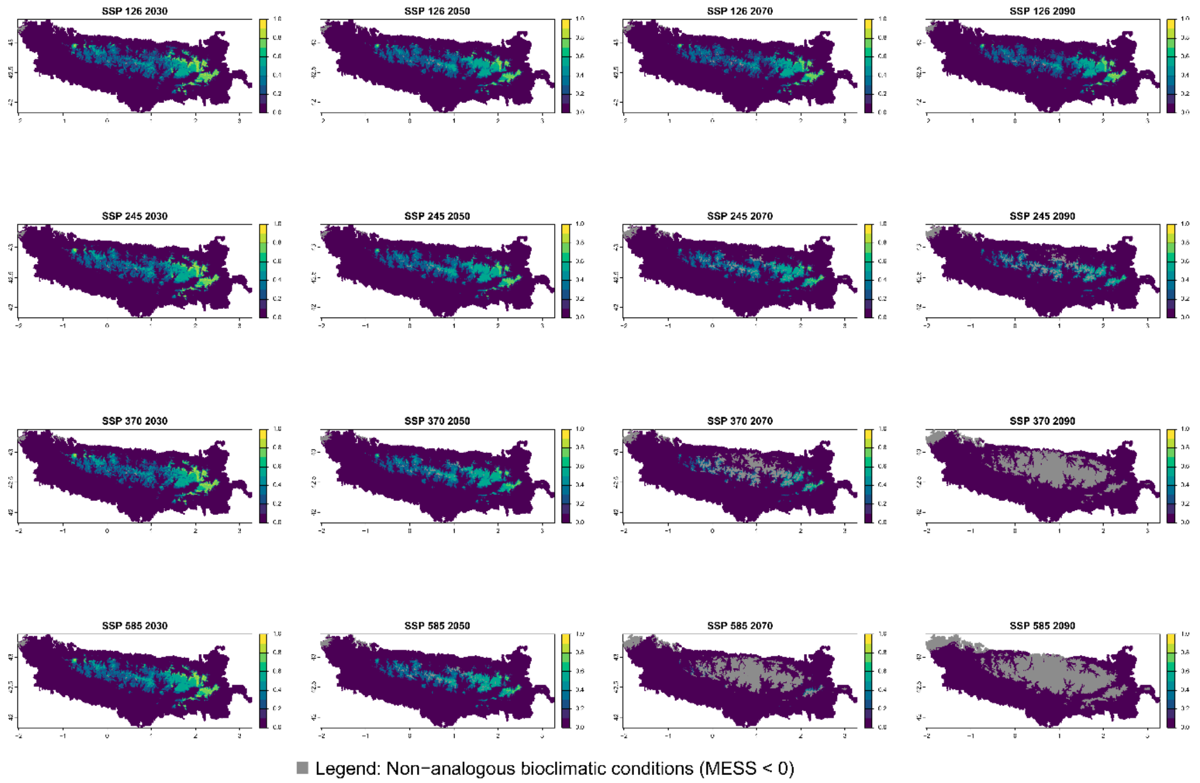

Current bioclimatic suitability in the Pyrenees for *Leucanthemum graminifolium*

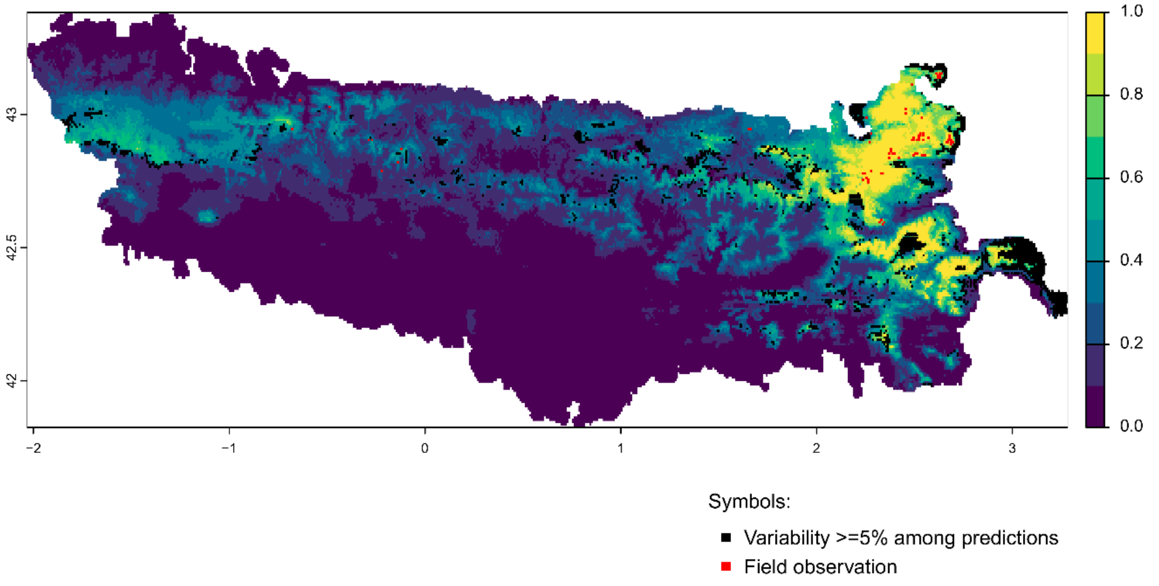


Future bioclimatic suitability in the Pyrenees by 2090 for *Leucanthemum graminifolium*

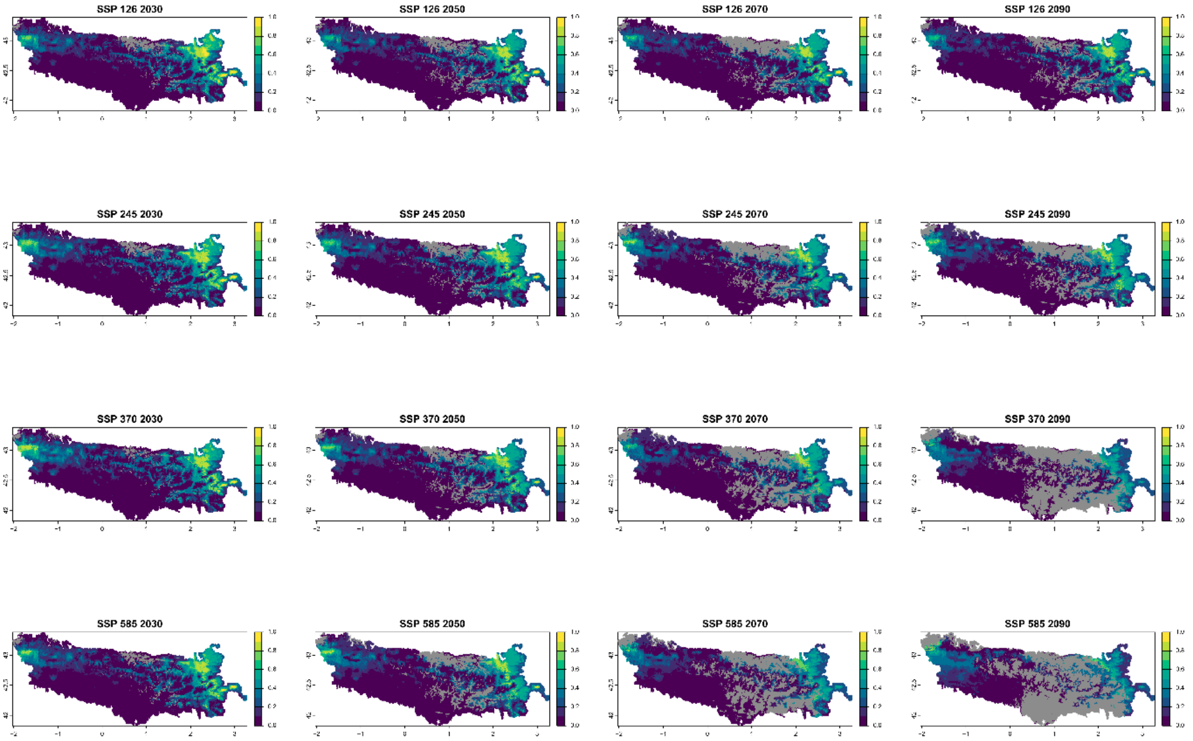


Legend: Non−analogous bioclimatic conditions (MESS < 0)

Current bioclimatic suitability in the Pyrenees for *Linaria bubanii*

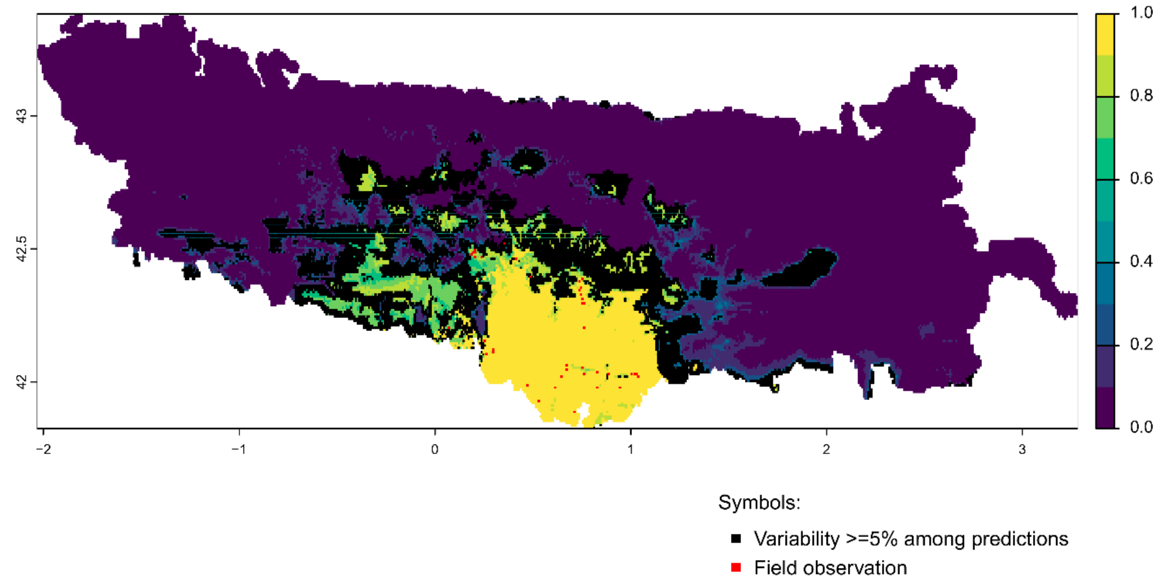


Future bioclimatic suitability in the Pyrenees by 2090 for *Linaria bubanii*

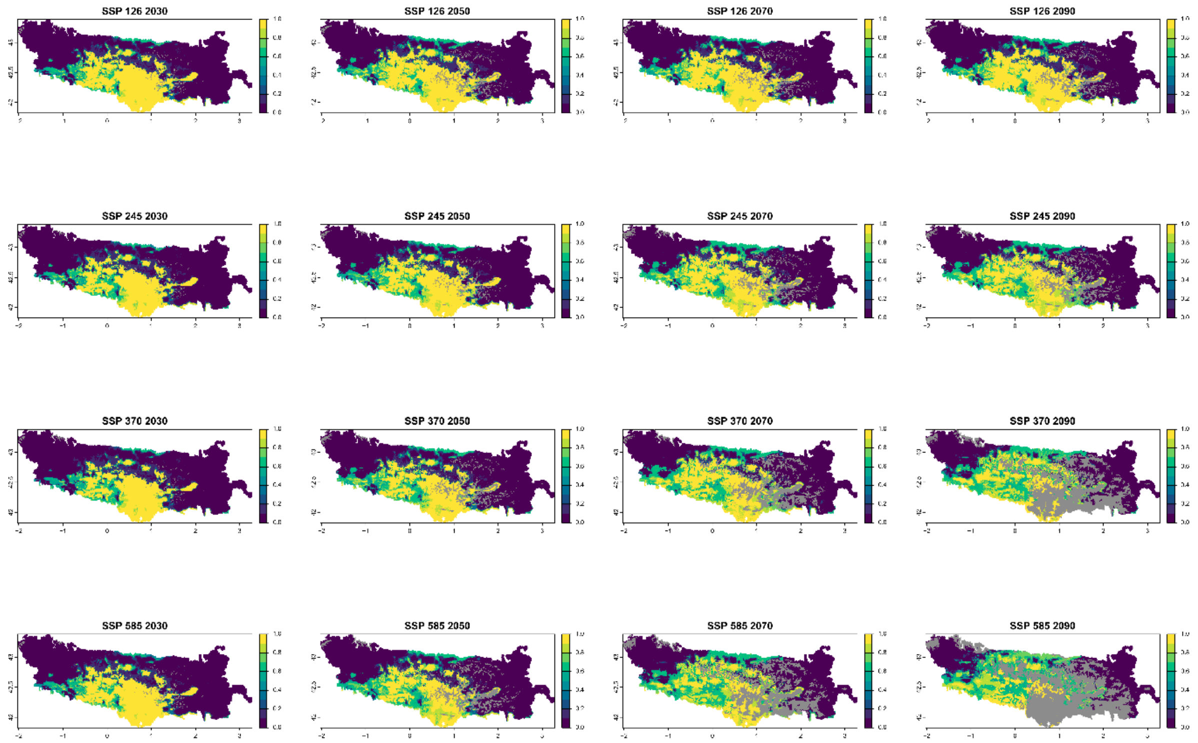


Legend: Non−analogous bioclimatic conditions (MESS < 0)

Current bioclimatic suitability in the Pyrenees for *Medicago hybrida*

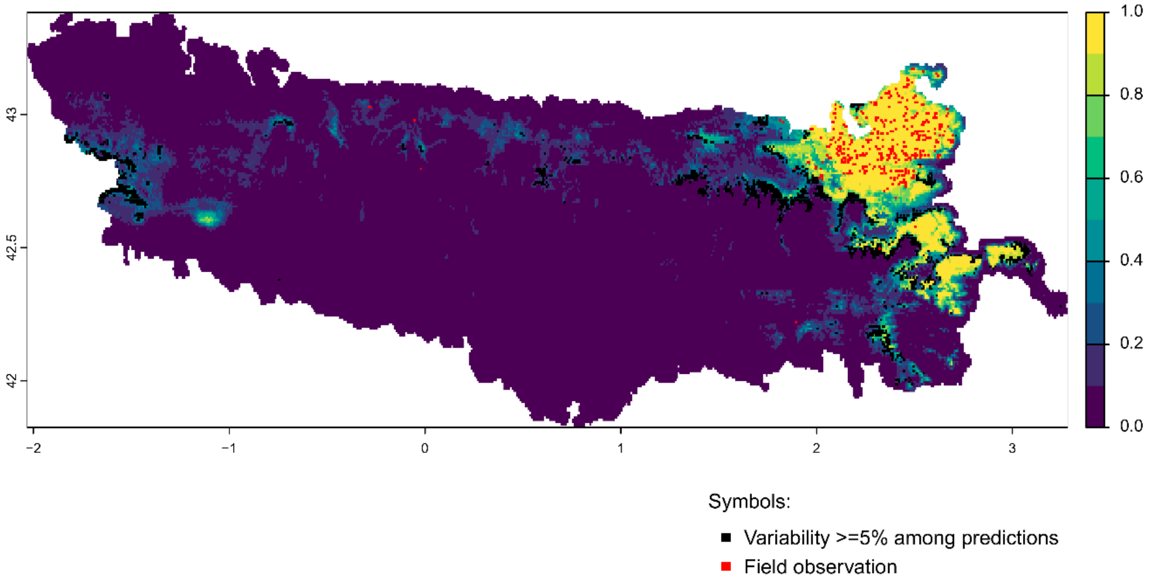


Future bioclimatic suitability in the Pyrenees by 2090 for *Medicago hybrida*

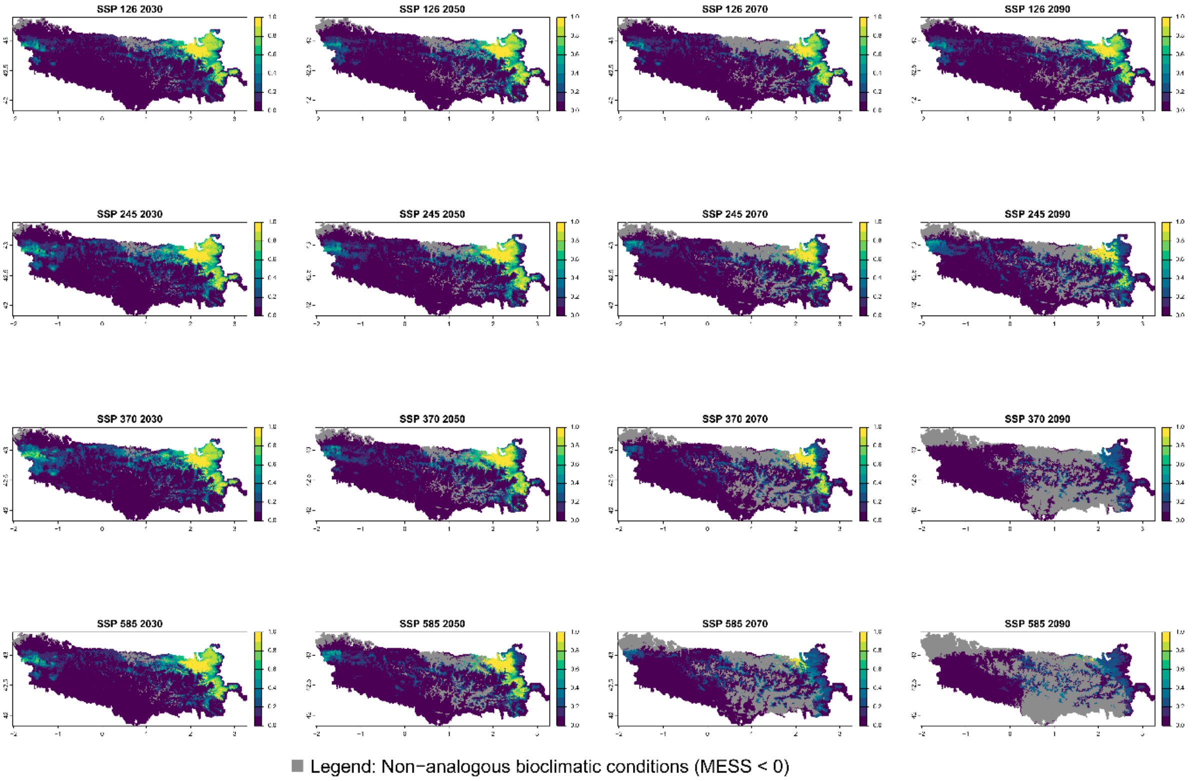

Current bioclimatic suitability in the Pyrenees for *Minuartia cerastiifolia*

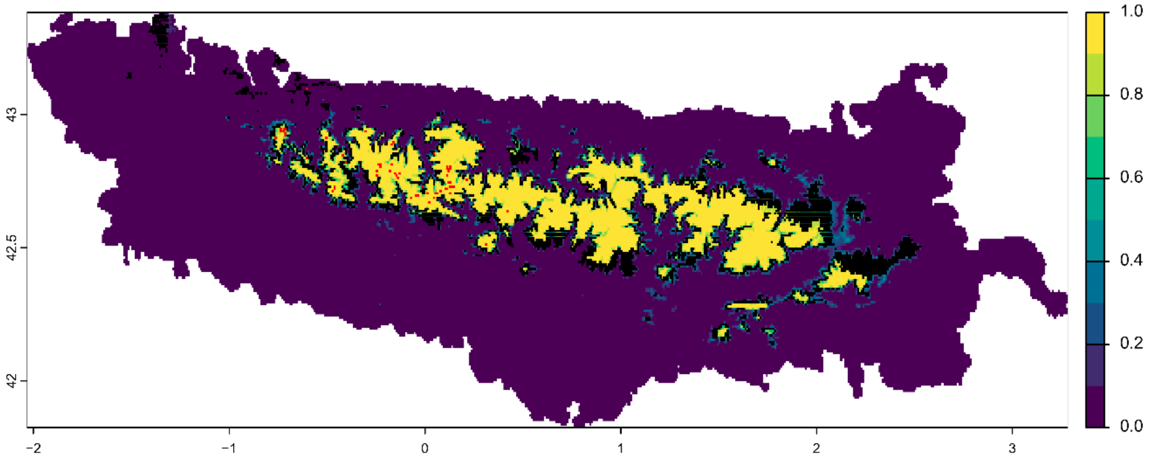


Symbols:

- ■ Variability >=5% among predictions
- ■ Field observation

Future bioclimatic suitability in the Pyrenees by 2090 for *Minuartia cerastiifolia*

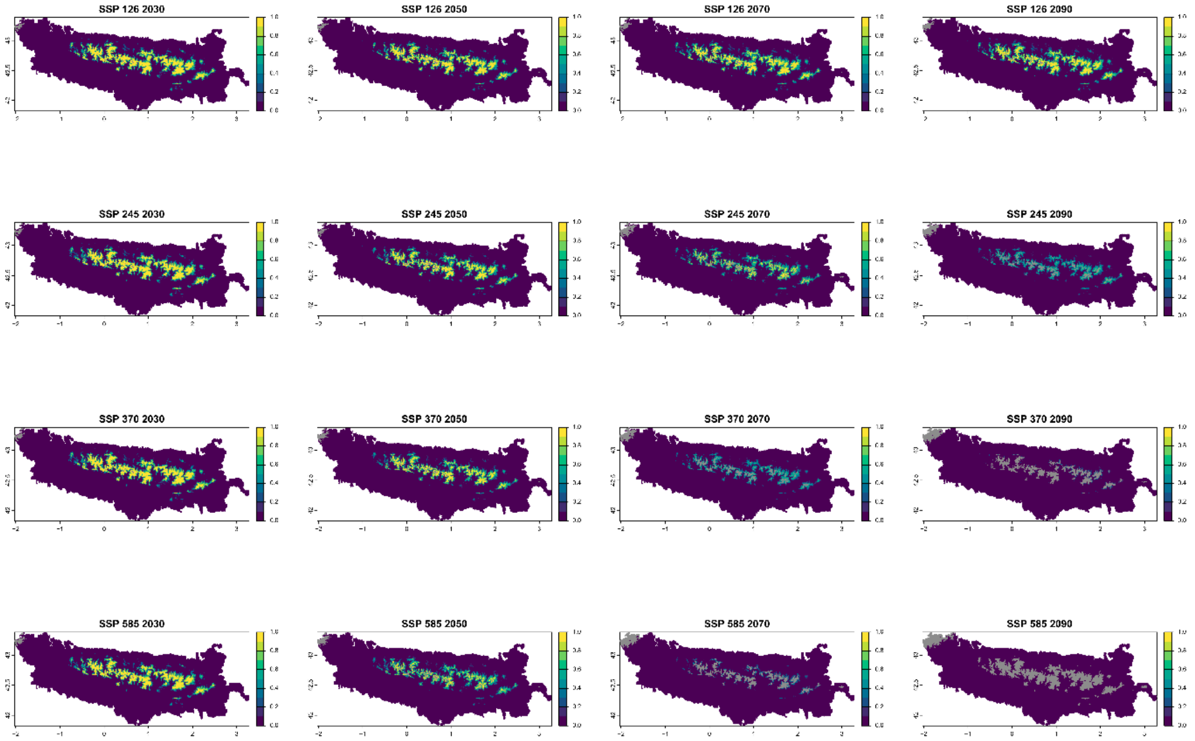


■ Legend: Non−analogous bioclimatic conditions (MESS < 0)

Current bioclimatic suitability in the Pyrenees for *Narcissus bicolor*

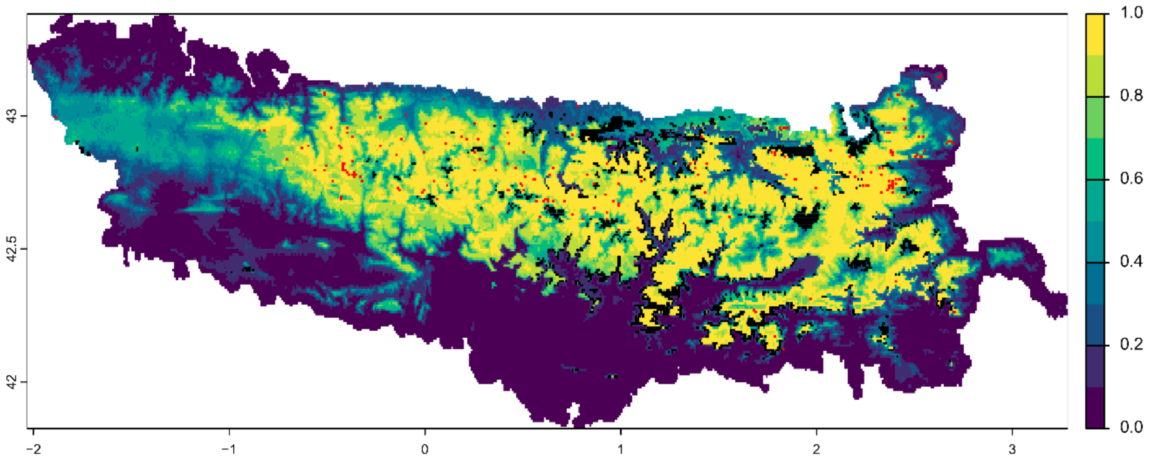


Symbols:

- Variability >=5% among predictions
- Field observation

Future bioclimatic suitability in the Pyrenees by 2090 for *Narcissus bicolor*

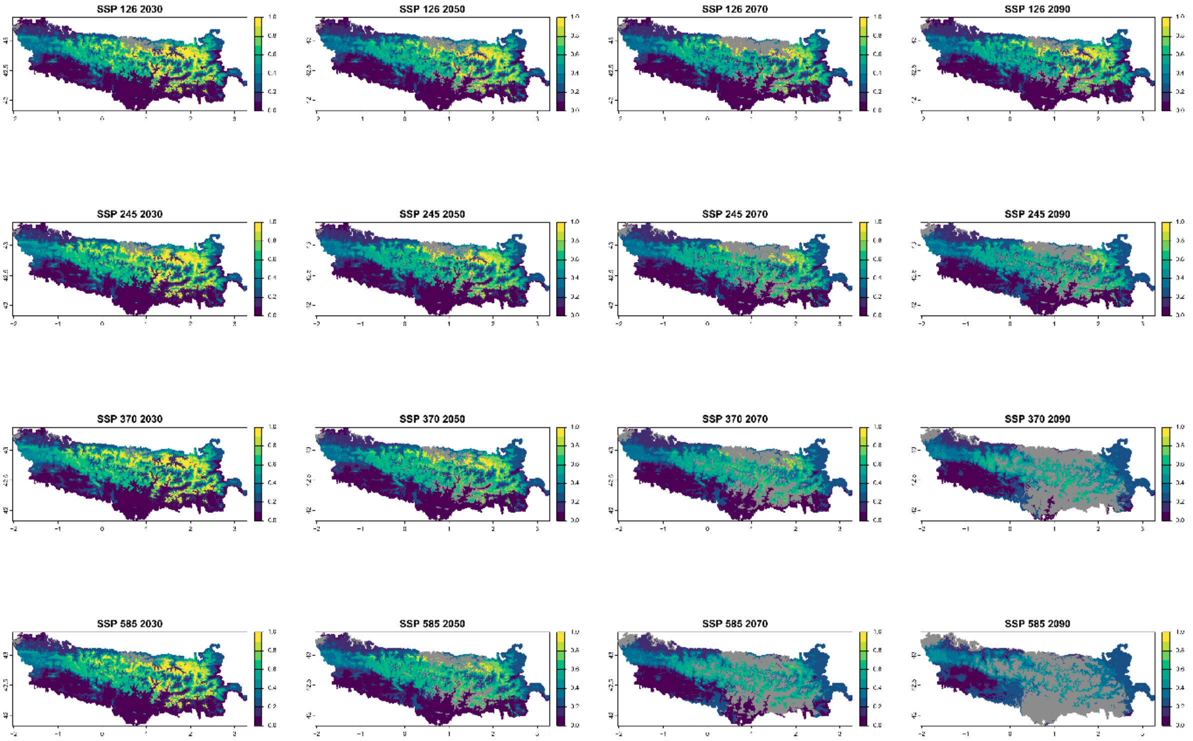


Legend: Non−analogous bioclimatic conditions (MESS < 0)

Current bioclimatic suitability in the Pyrenees for *Onobrychis pyrenaica*

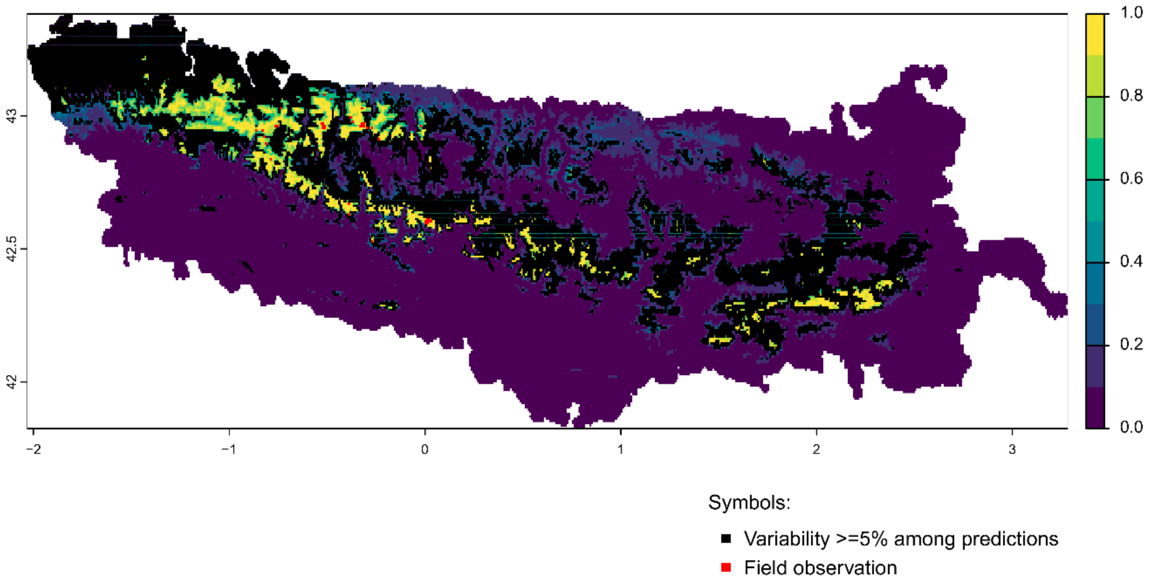


Future bioclimatic suitability in the Pyrenees by 2090 for *Onobrychis pyrenaica*

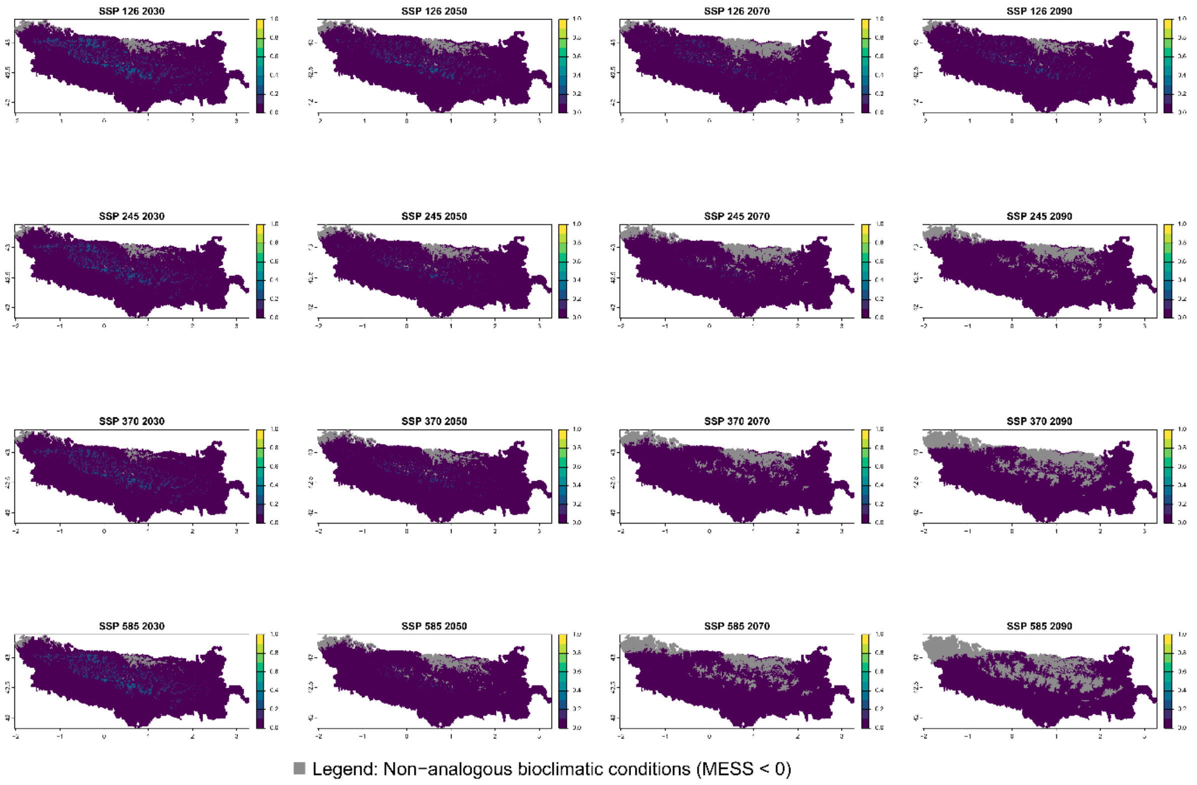

Current bioclimatic suitability in the Pyrenees for *Petrocoptis crassifolia*

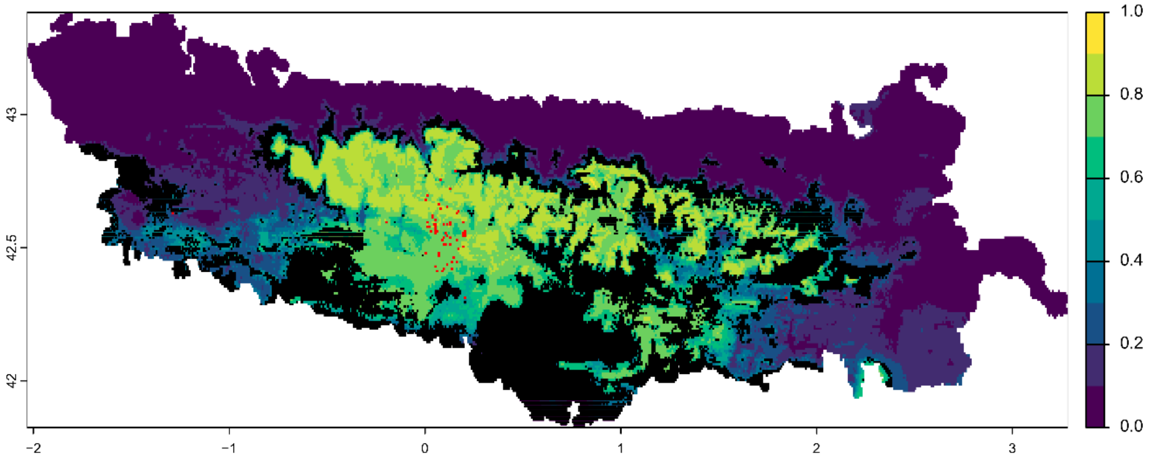


Symbols:

■ Variability >=5% among predictions

■ Field observation

Future bioclimatic suitability in the Pyrenees by 2090 for *Petrocoptis crassifolia*

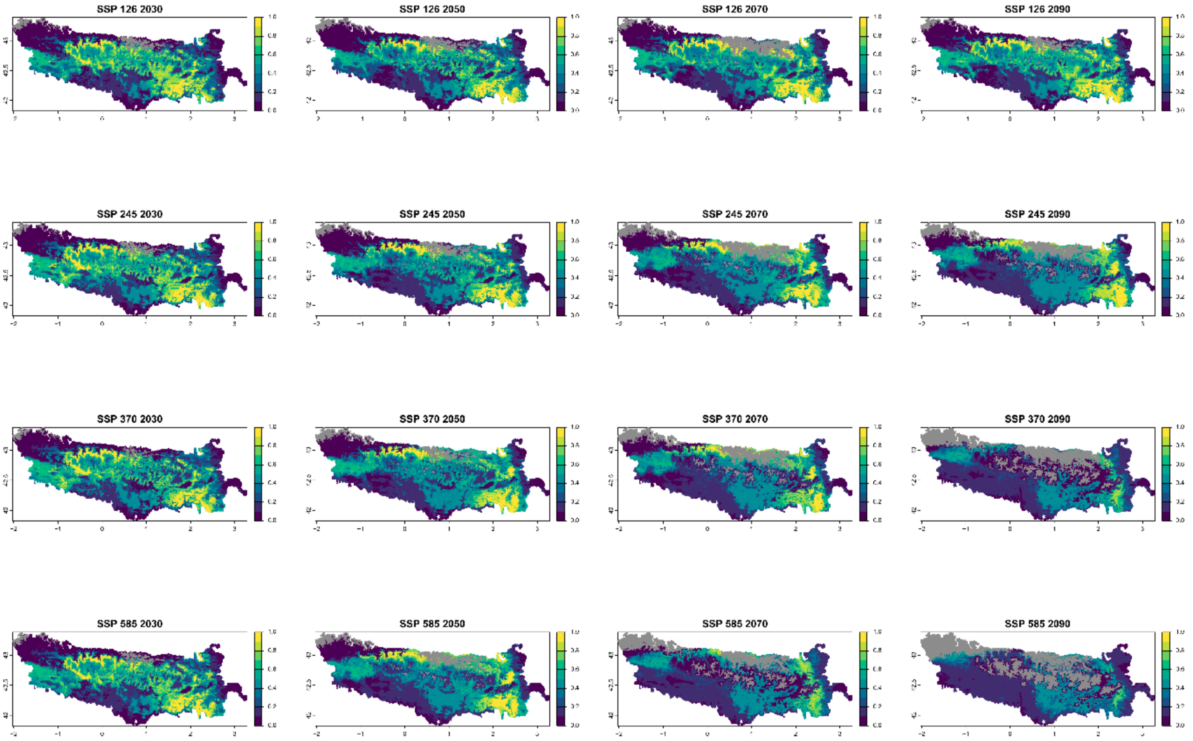


■ Legend: Non−analogous bioclimatic conditions (MESS < 0)

Current bioclimatic suitability in the Pyrenees for *Petrocoptis hispanica*

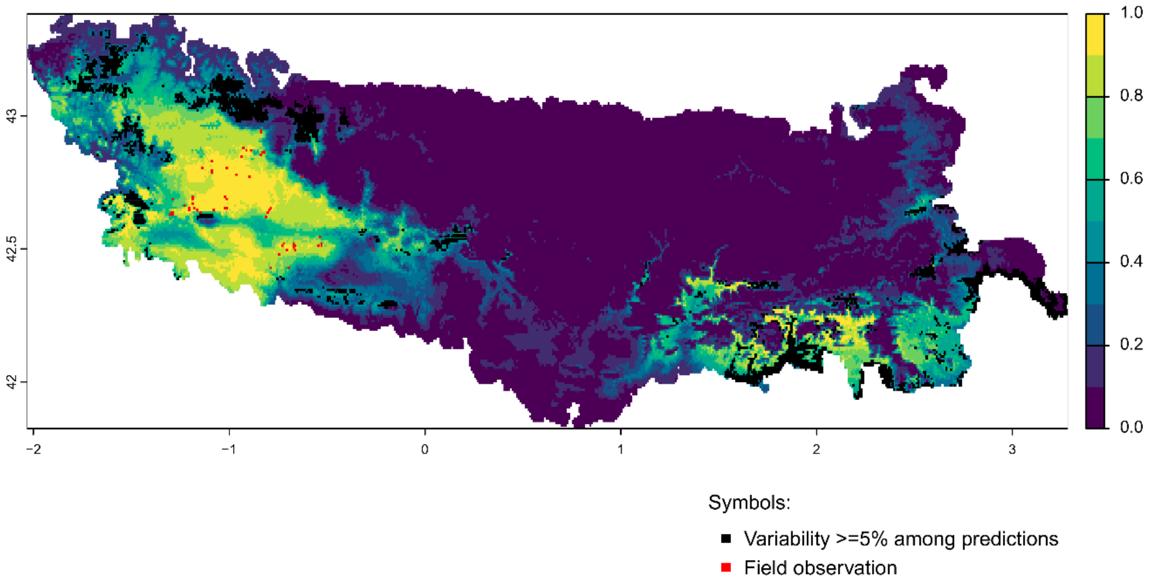


Future bioclimatic suitability in the Pyrenees by 2090 for *Petrocoptis hispanica*

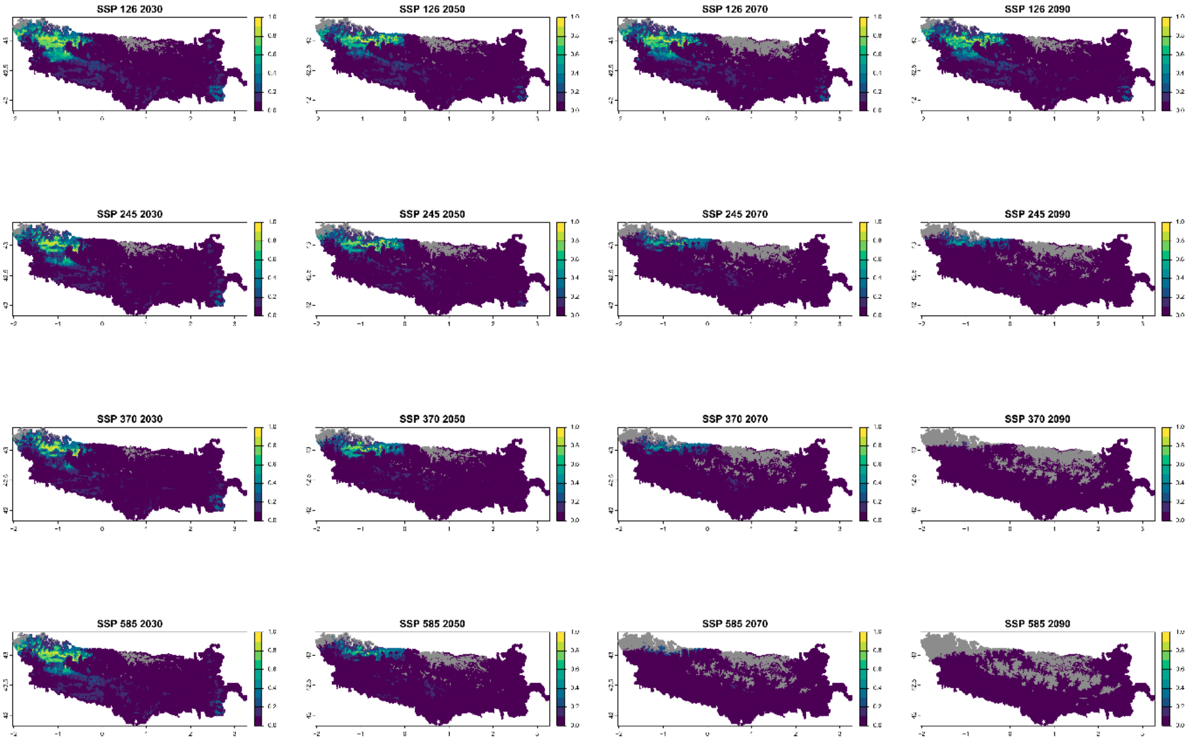


Legend: Non−analogous bioclimatic conditions (MESS < 0)

Current bioclimatic suitability in the Pyrenees for *Petrocoptis montsicciana*

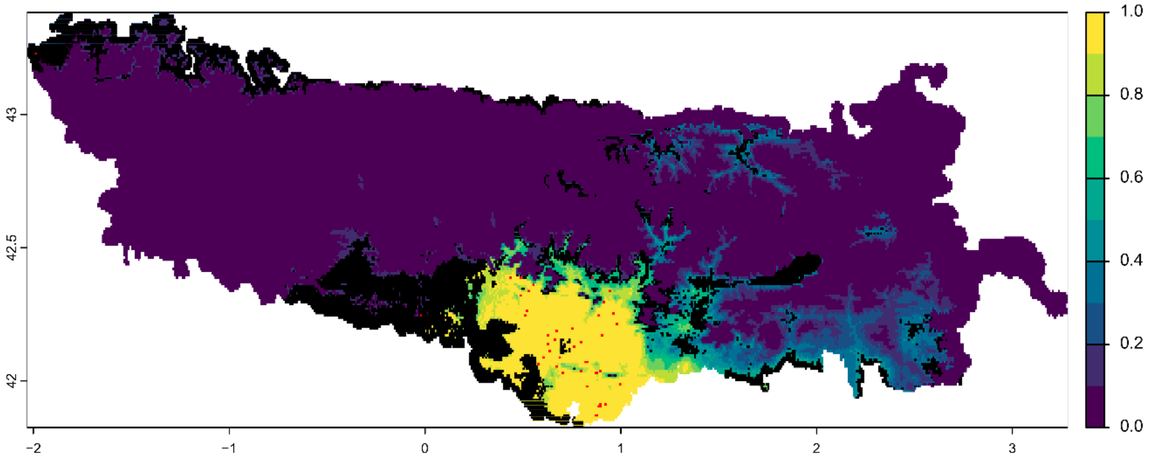


Symbols:

■ Variability >=5% among predictions

■ Field observation

Future bioclimatic suitability in the Pyrenees by 2090 for *Petrocoptis montsicciana*

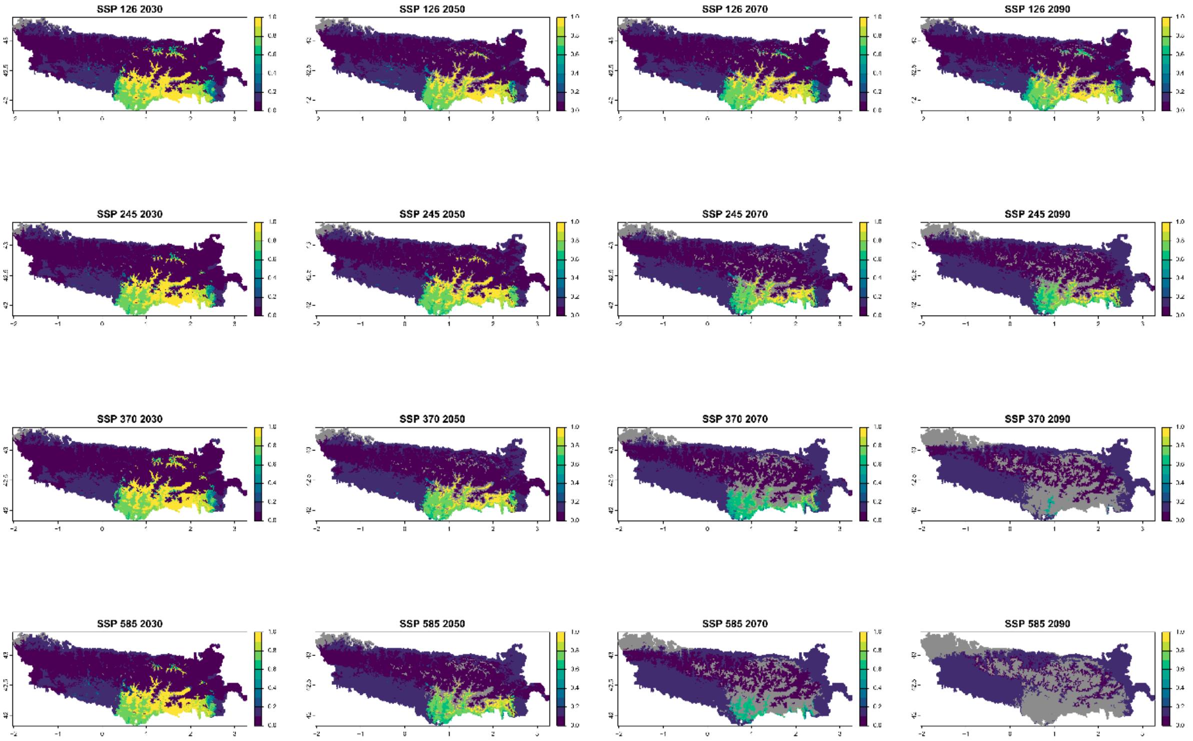


■ Legend: Non−analogous bioclimatic conditions (MESS < 0)

Current bioclimatic suitability in the Pyrenees for *Pinguicula longifolia*

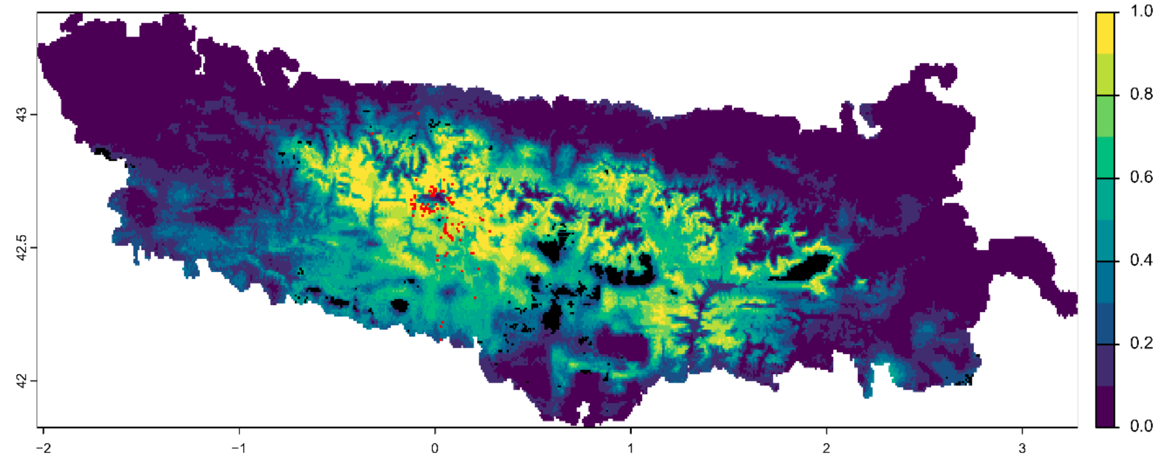


Symbols:

■ Variability >=5% among predictions

■ Field observation

Future bioclimatic suitability in the Pyrenees by 2090 for *Pinguicula longifolia*

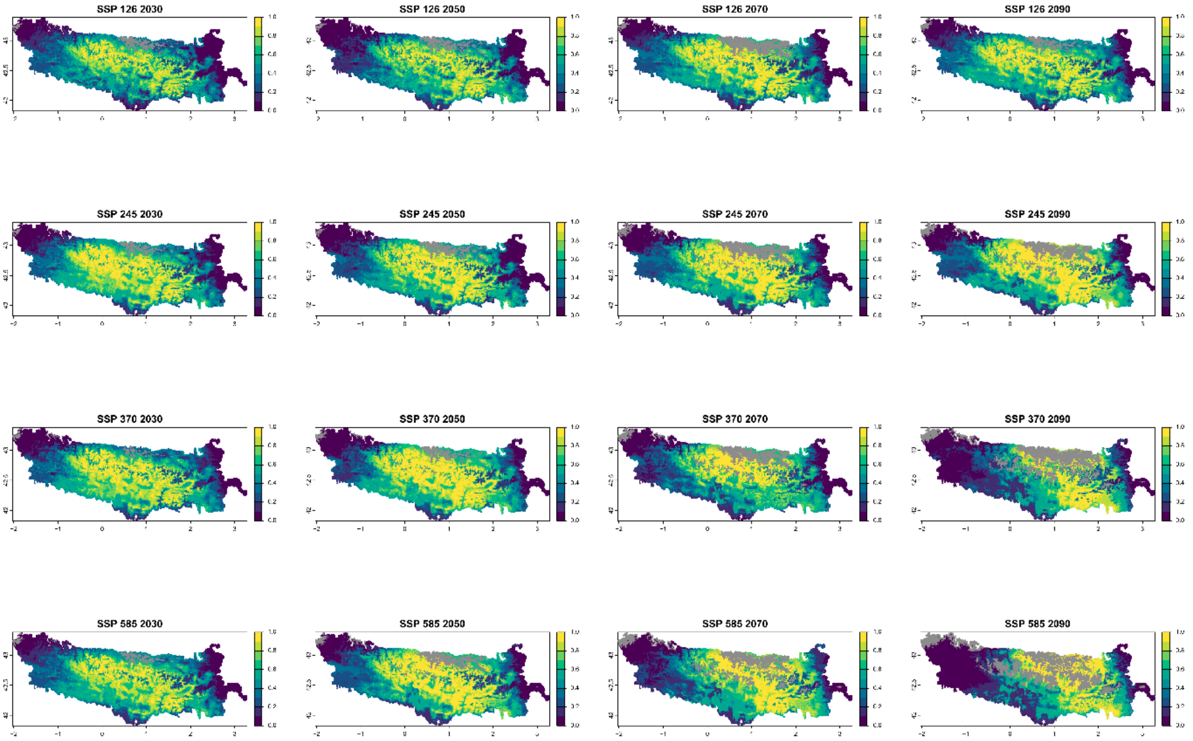


■ Legend: Non−analogous bioclimatic conditions (MESS < 0)

Current bioclimatic suitability in the Pyrenees for *Ramonda myconi*

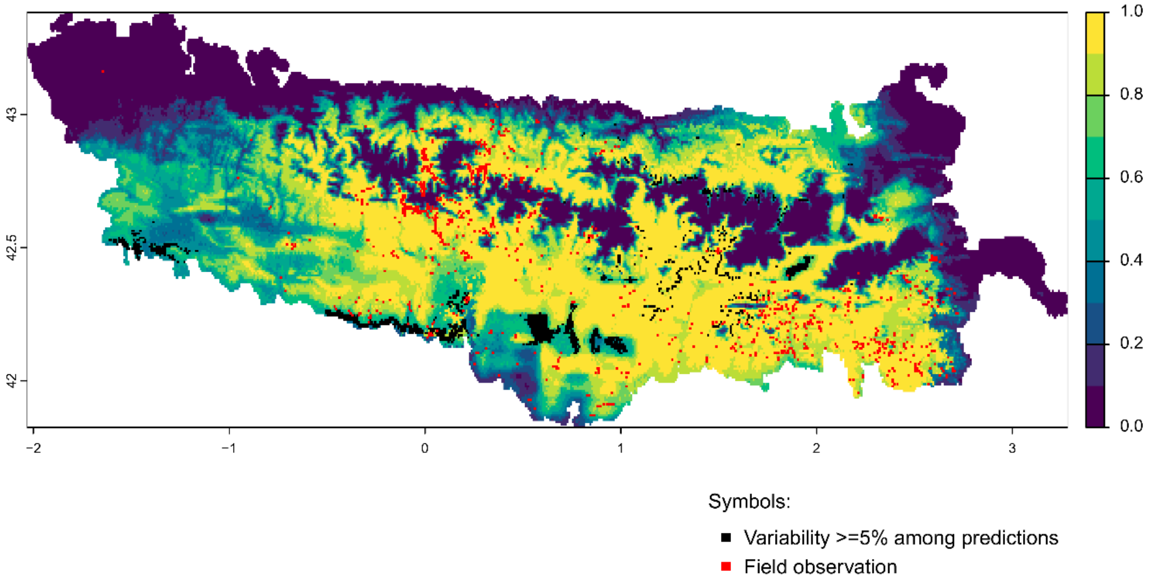


Future bioclimatic suitability in the Pyrenees by 2090 for *Ramonda myconi*

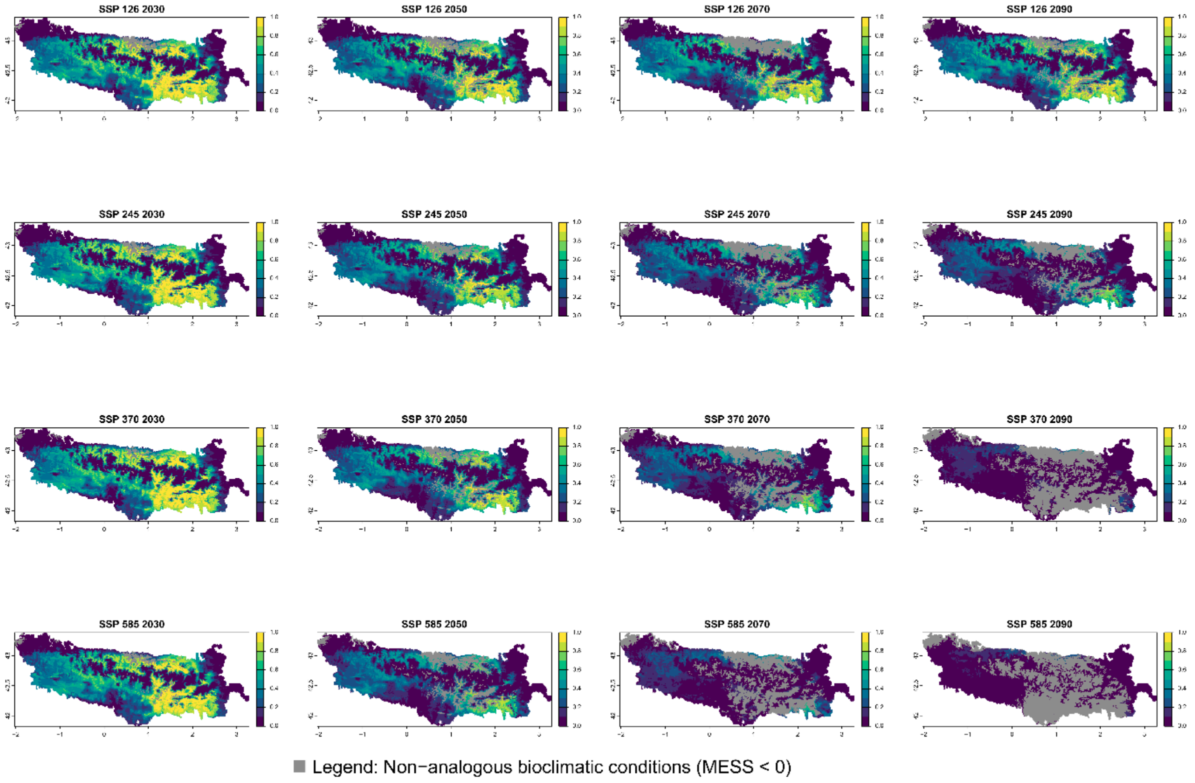

Current bioclimatic suitability in the Pyrenees for *Ranunculus pyrenaeus*

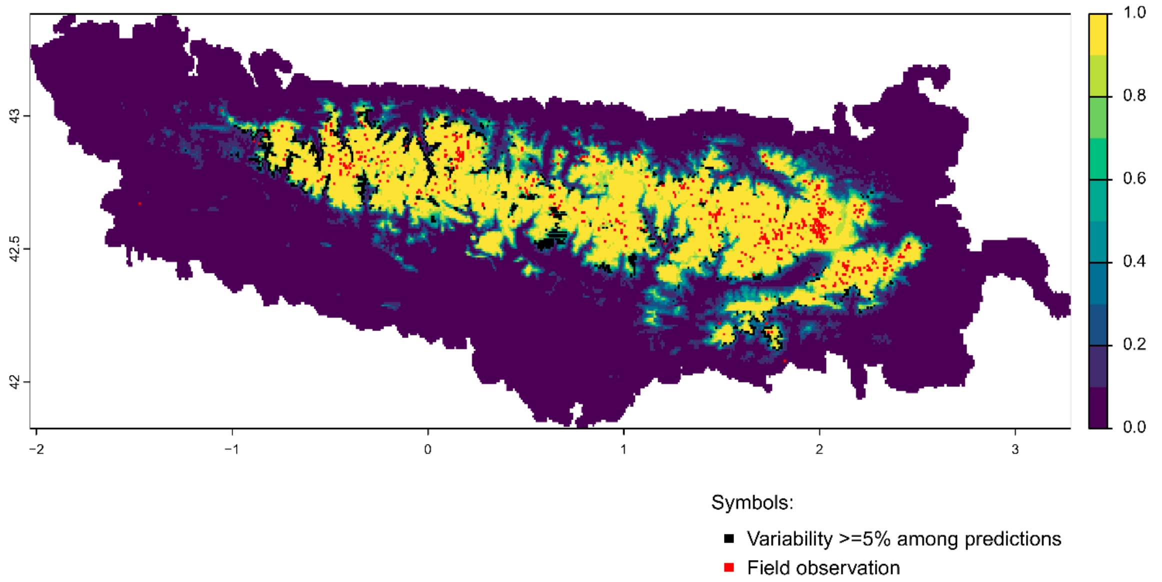


Future bioclimatic suitability in the Pyrenees by 2090 for *Ranunculus pyrenaeus*

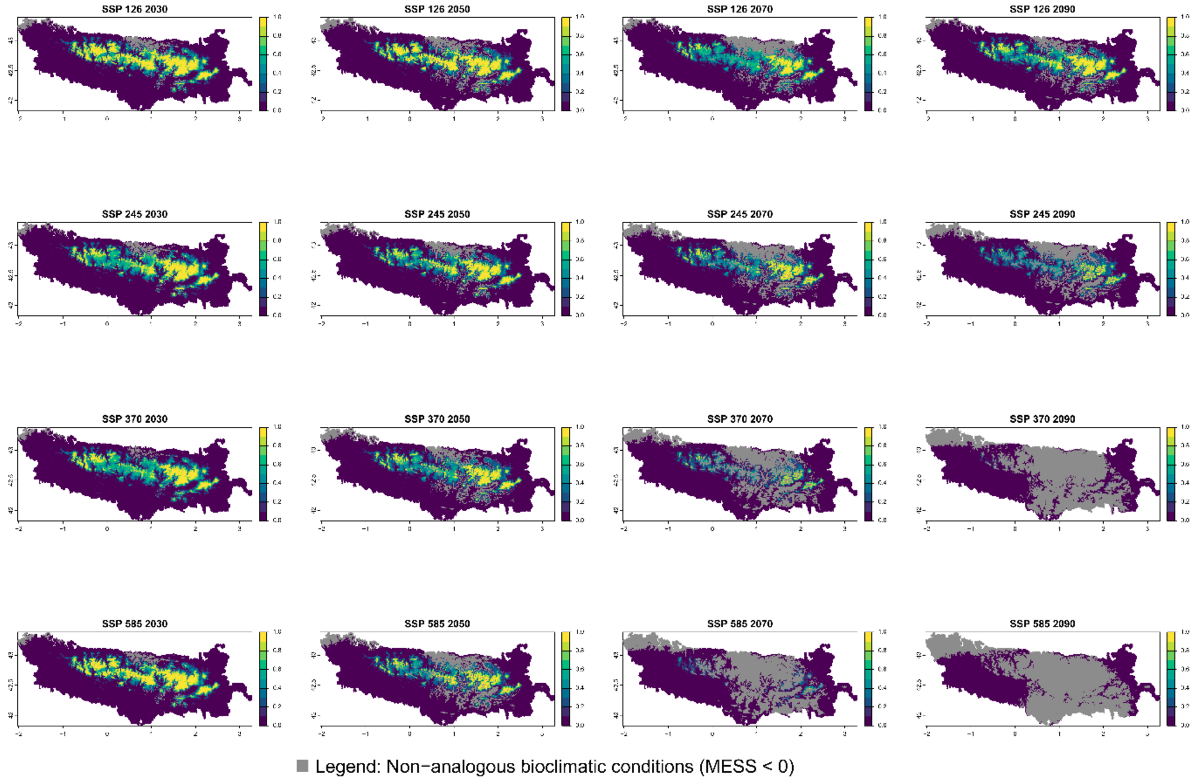

### Current bioclimatic suitability in the Pyrenees for *Ranunculus ruscinonensis*

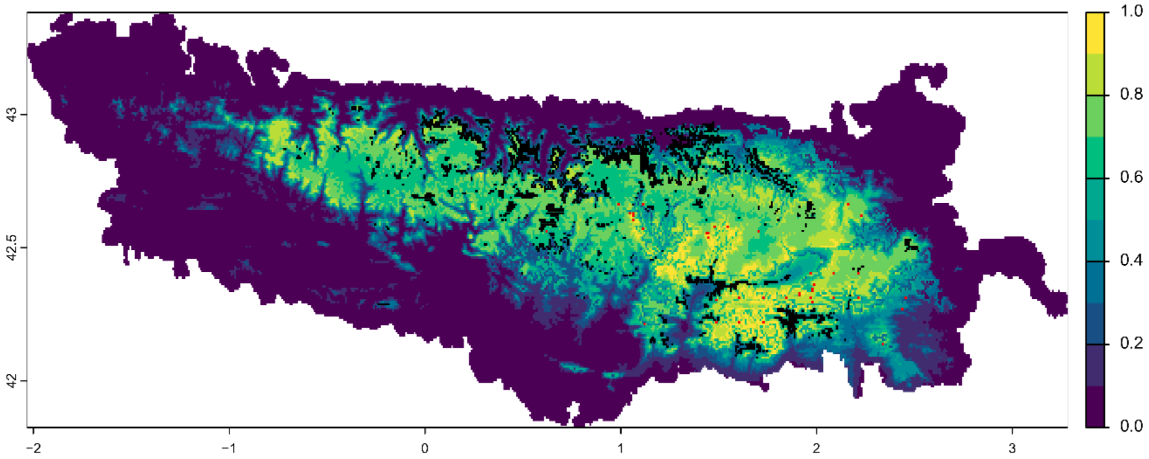


Symbols:

- ■ Variability >=5% among predictions
- ■ Field observation

Future bioclimatic suitability in the Pyrenees by 2090 for *Ranunculus ruscinonensis*

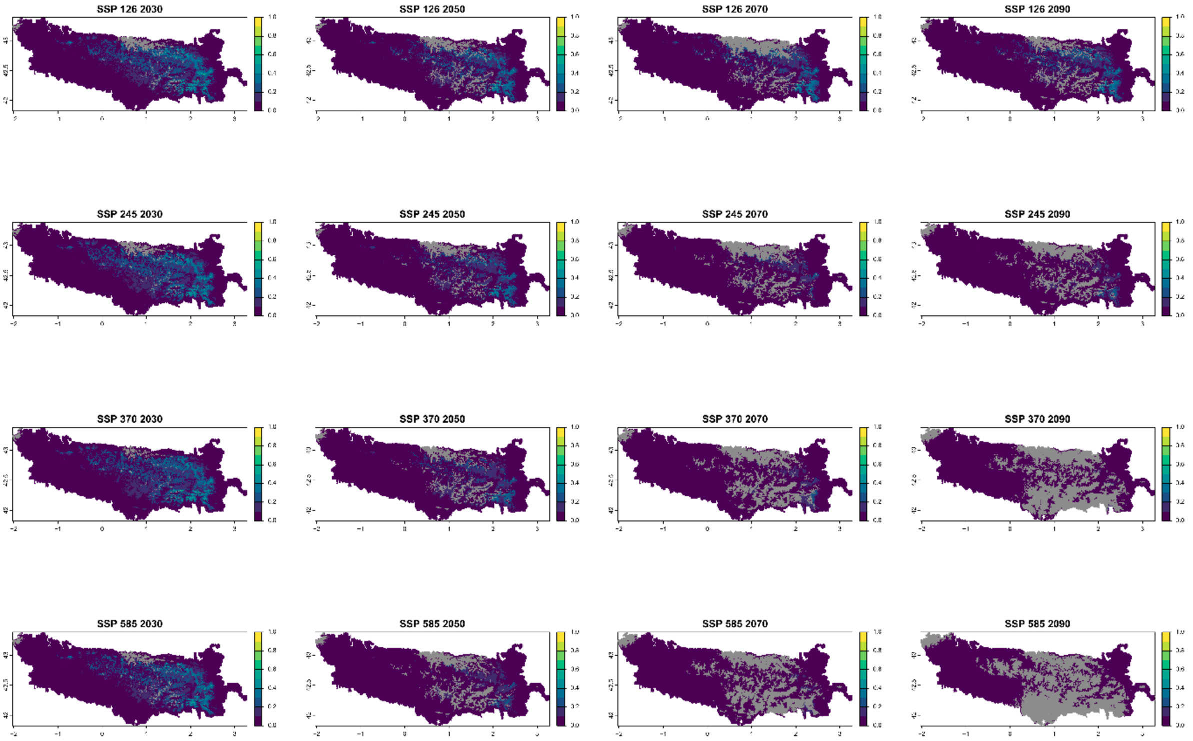


■ Legend: Non−analogous bioclimatic conditions (MESS < 0)

### Current bioclimatic suitability in the Pyrenees for *Rhaponticum centauroides*

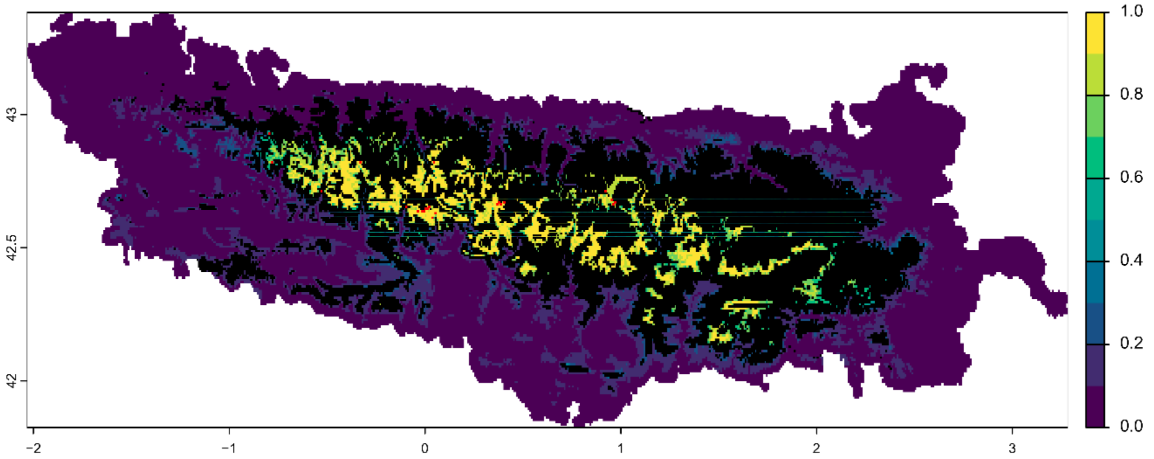


Symbols:

■ Variability >=5% among predictions

■ Field observation

#### Future bioclimatic suitability in the Pyrenees by 2090 for *Rhaponticum centauroides*

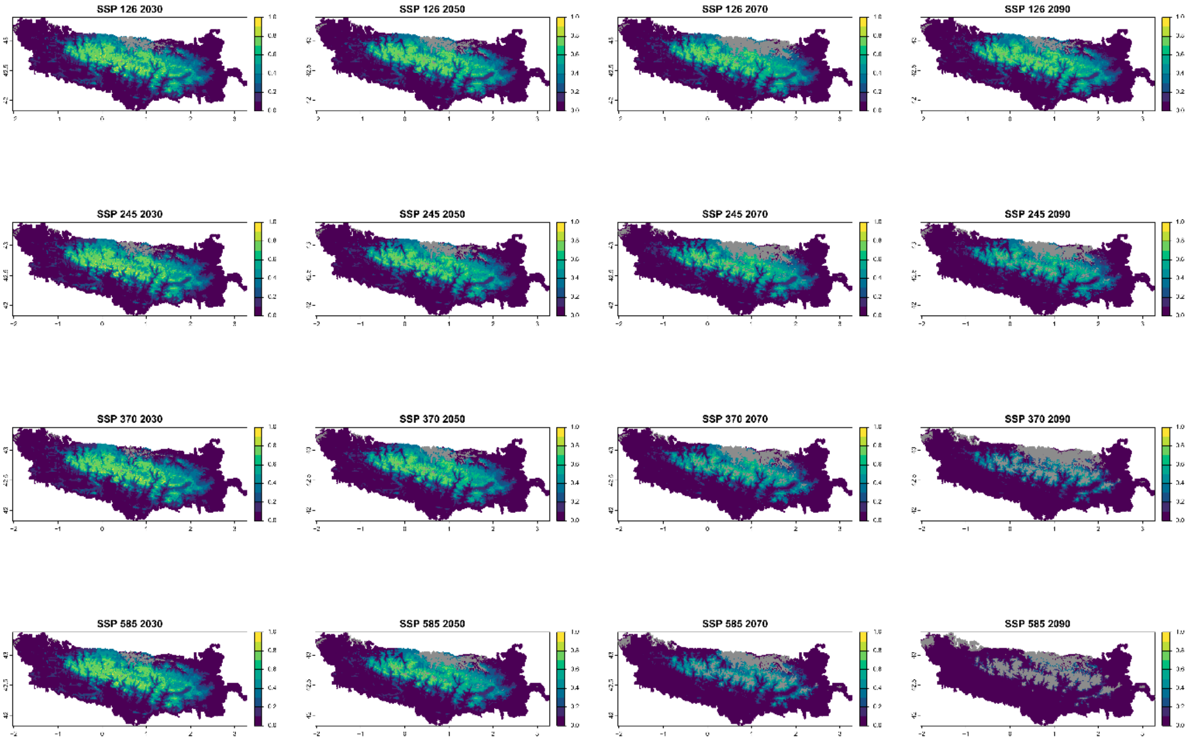


■ Legend: Non−analogous bioclimatic conditions (MESS < 0)

Current bioclimatic suitability in the Pyrenees for *Salix pyrenaica*

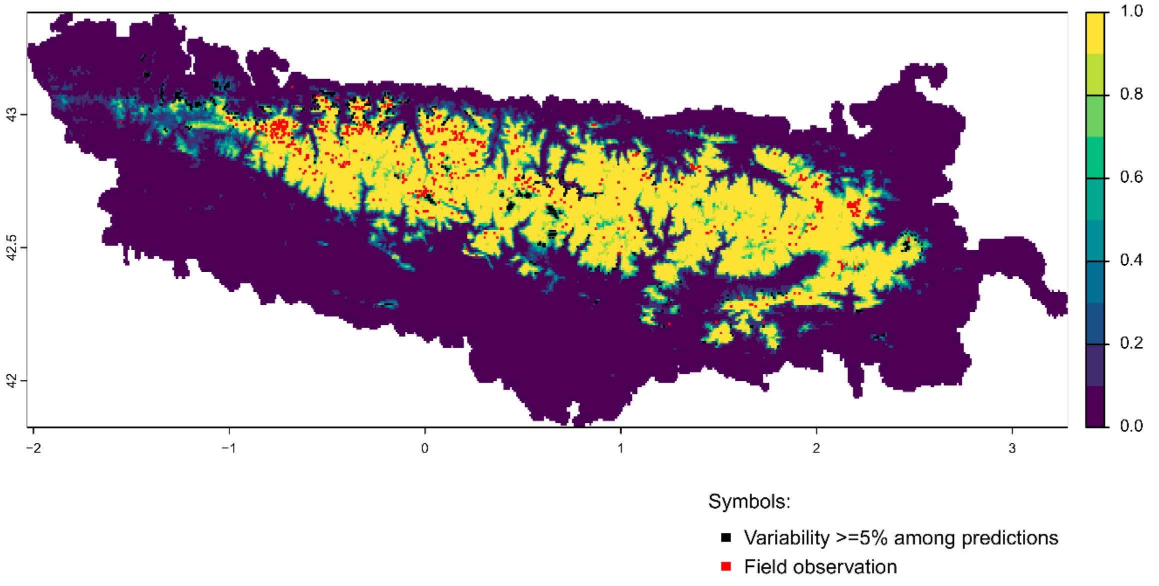


Future bioclimatic suitability in the Pyrenees by 2090 for *Salix pyrenaica*

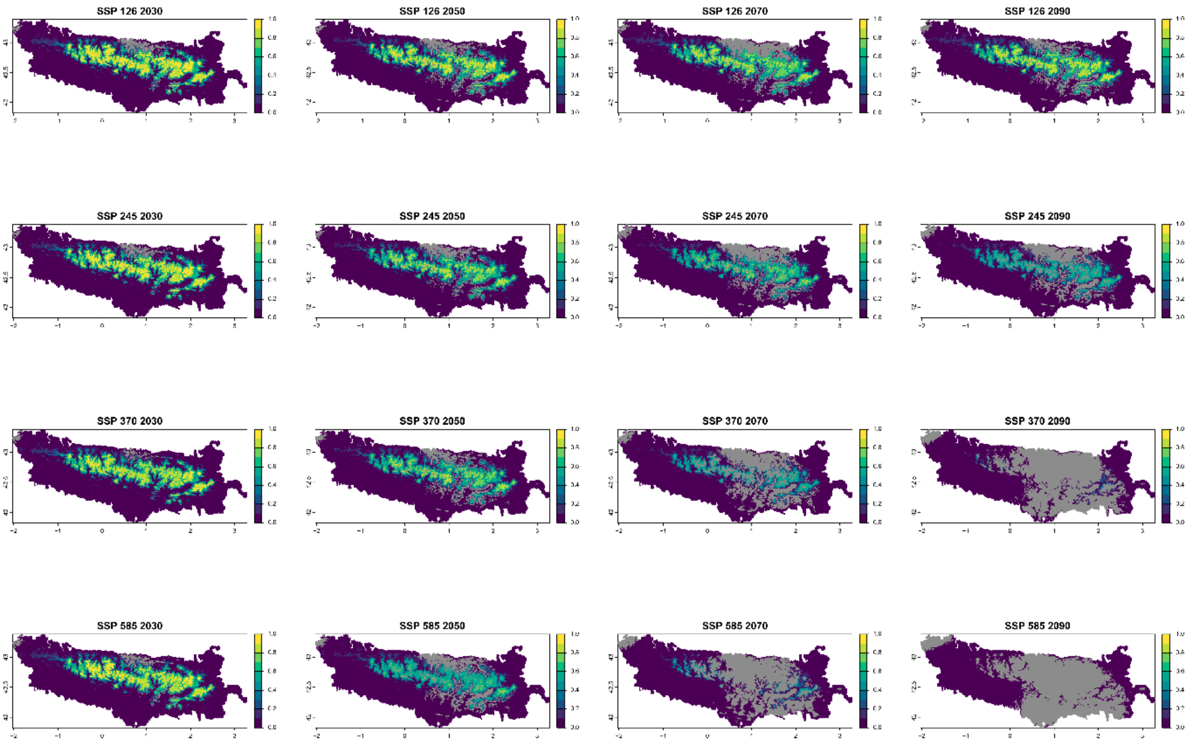


Legend: Non−analogous bioclimatic conditions (MESS < 0)

Current bioclimatic suitability in the Pyrenees for *Santolina benthamiana*

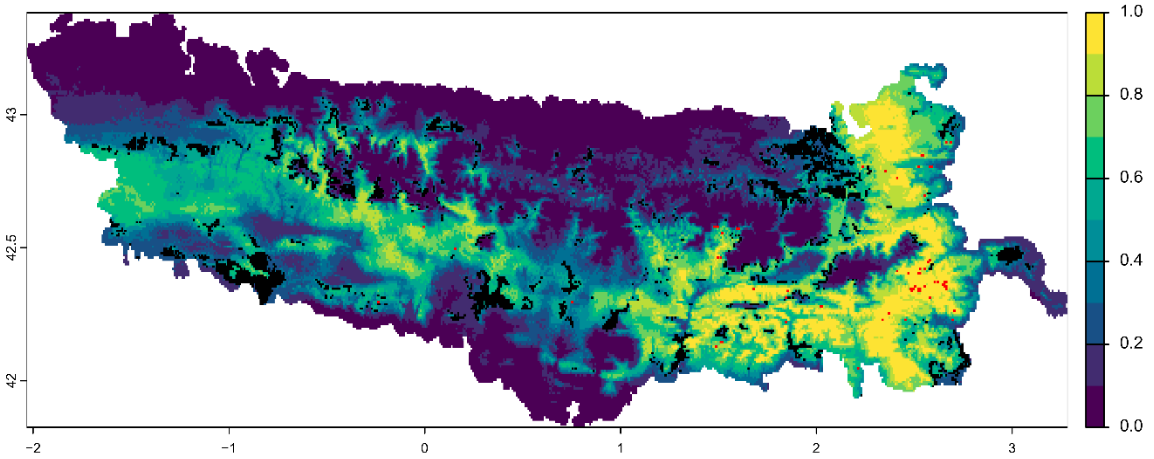


Symbols:

- Variability >=5% among predictions
- Field observation

Future bioclimatic suitability in the Pyrenees by 2090 for *Santolina benthamiana*

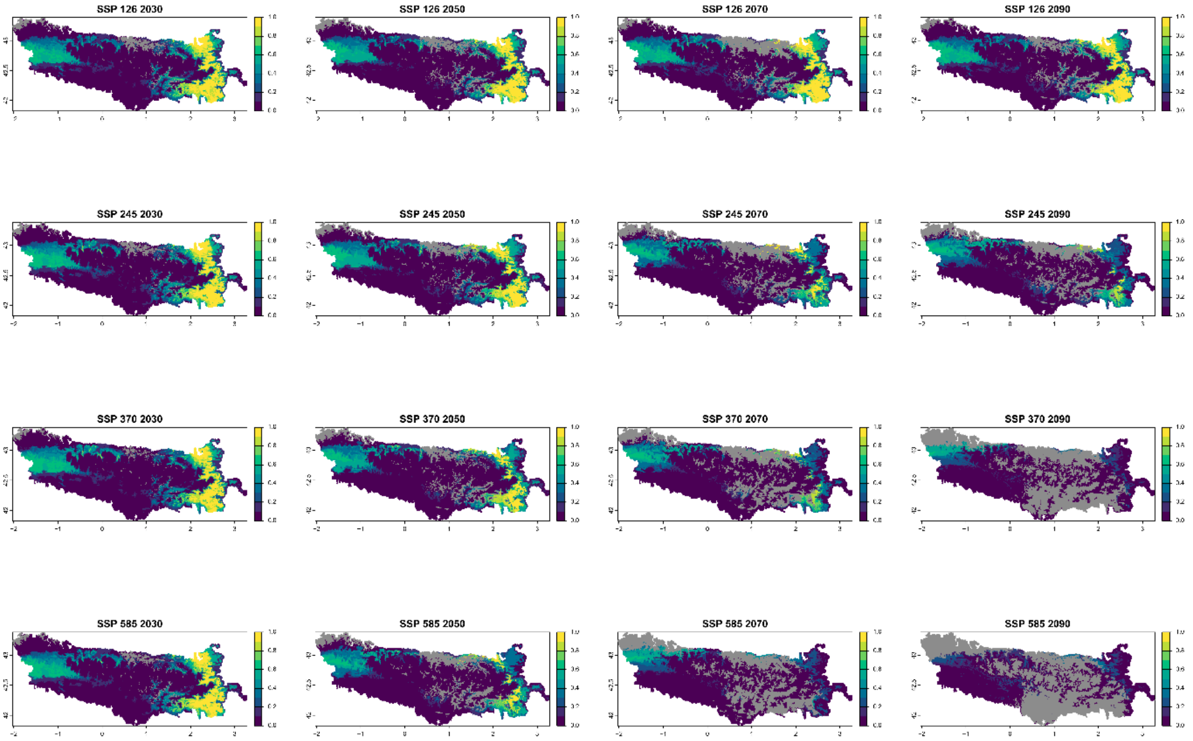


Legend: Non−analogous bioclimatic conditions (MESS < 0)

Current bioclimatic suitability in the Pyrenees for *Saponaria caespitosa*

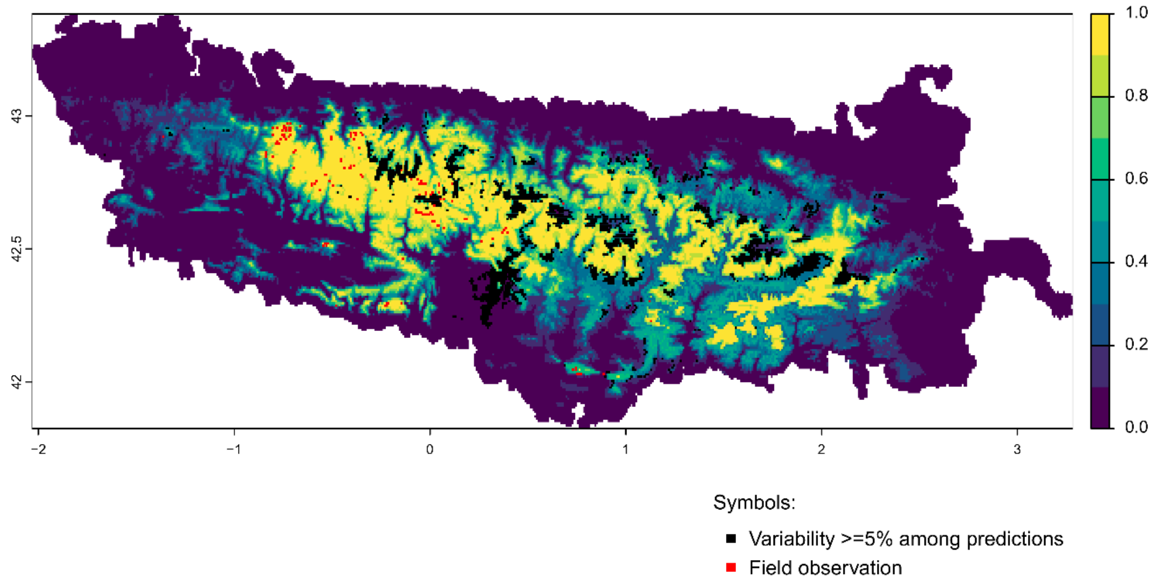


Future bioclimatic suitability in the Pyrenees by 2090 for *Saponaria caespitosa*

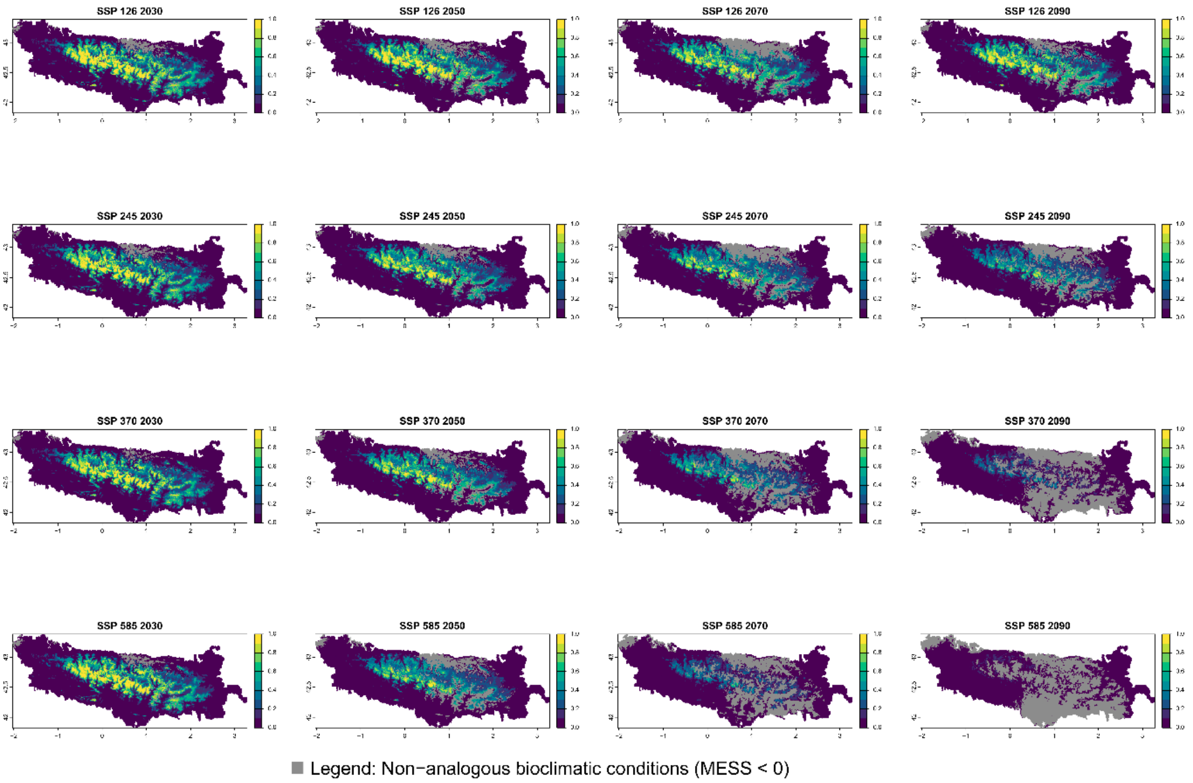

Current bioclimatic suitability in the Pyrenees for *Saxifraga aquatica*

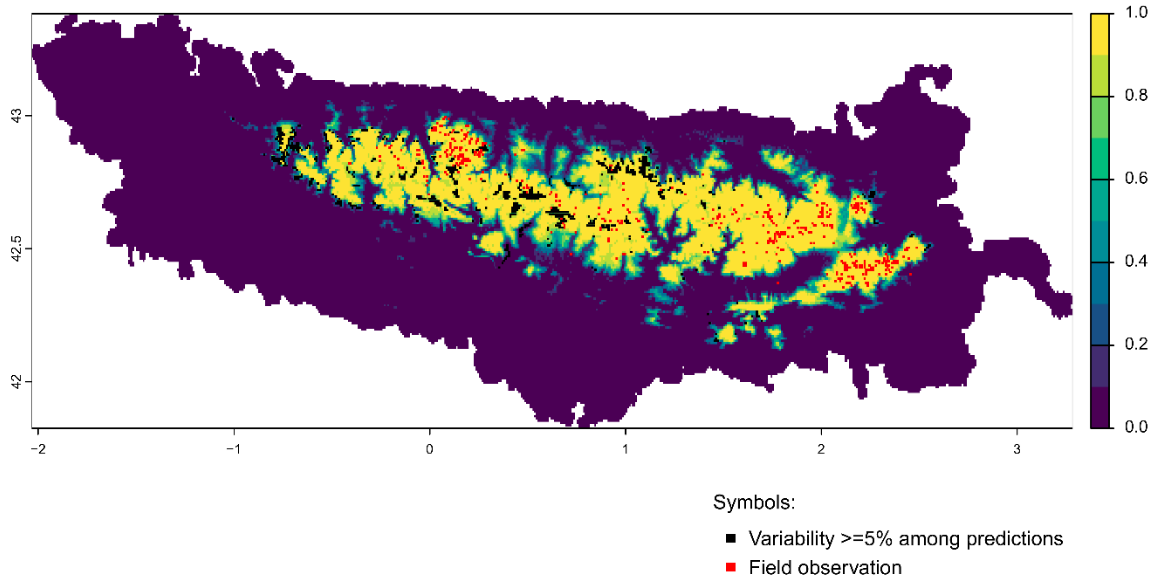


Future bioclimatic suitability in the Pyrenees by 2090 for *Saxifraga aquatica*

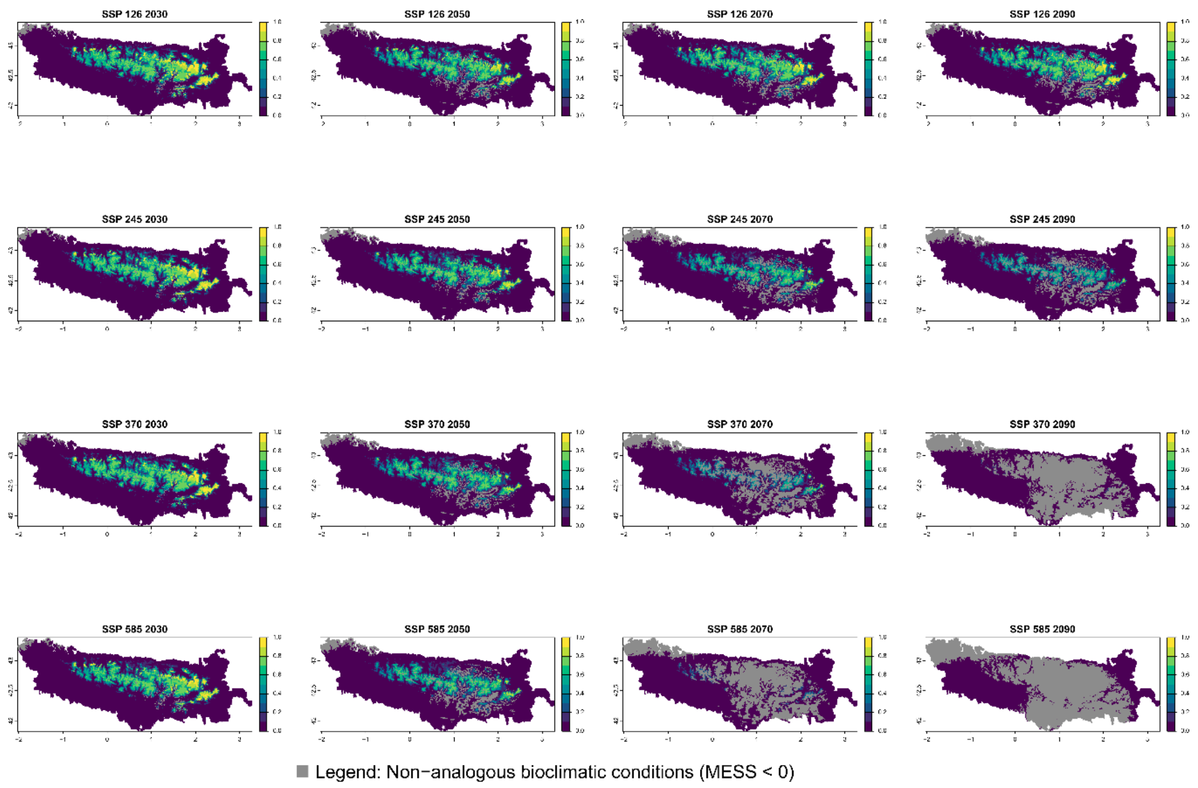

### Current bioclimatic suitability in the Pyrenees for *Saxifraga aretioides*

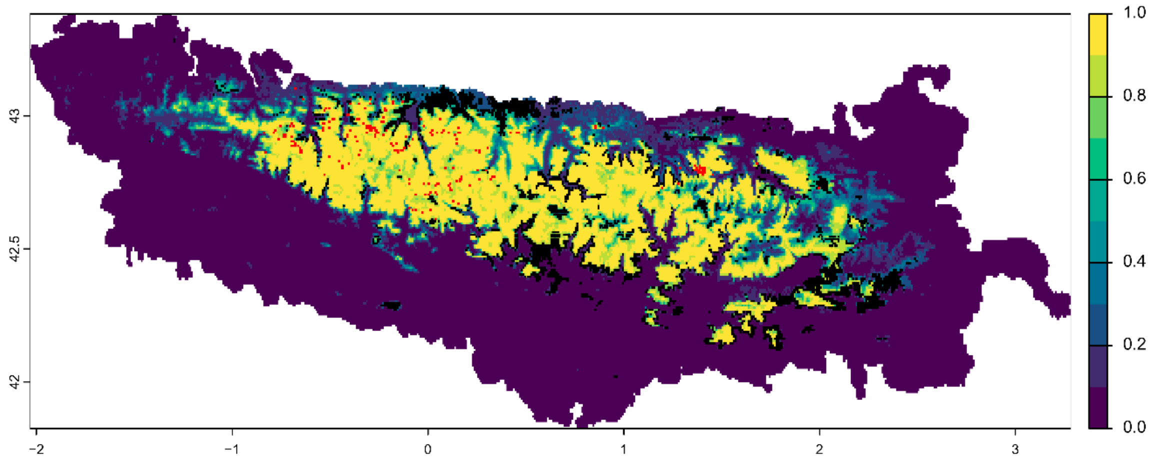


Symbols:

- ■ Variability >=5% among predictions
- ■ Field observation

Future bioclimatic suitability in the Pyrenees by 2090 for *Saxifraga aretioides*

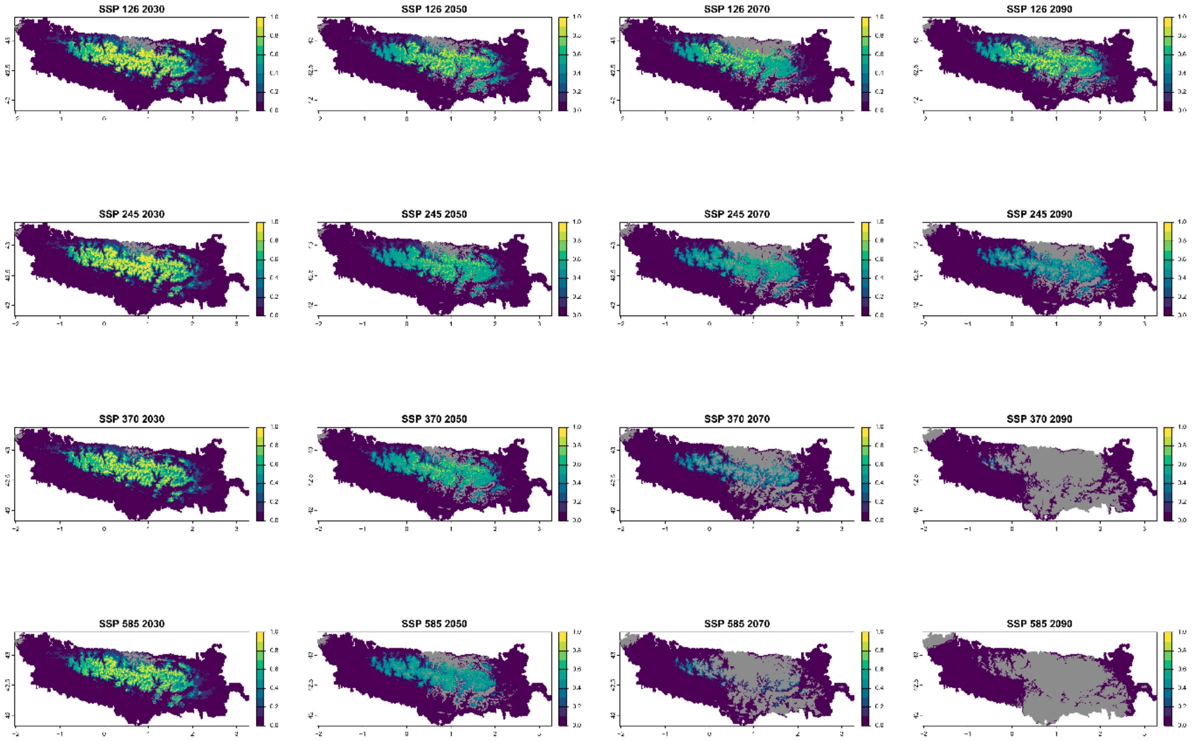


■ Legend: Non−analogous bioclimatic conditions (MESS < 0)

### Current bioclimatic suitability in the Pyrenees for *Saxifraga geranioides*

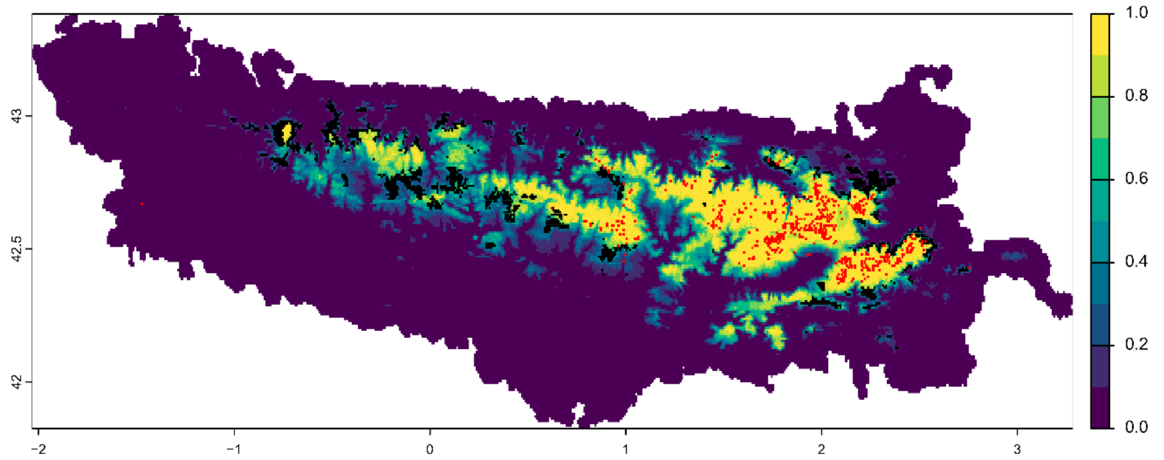


Symbols:

- ■ Variability >=5% among predictions
- ■ Field observation

Future bioclimatic suitability in the Pyrenees by 2090 for *Saxifraga geranioides*

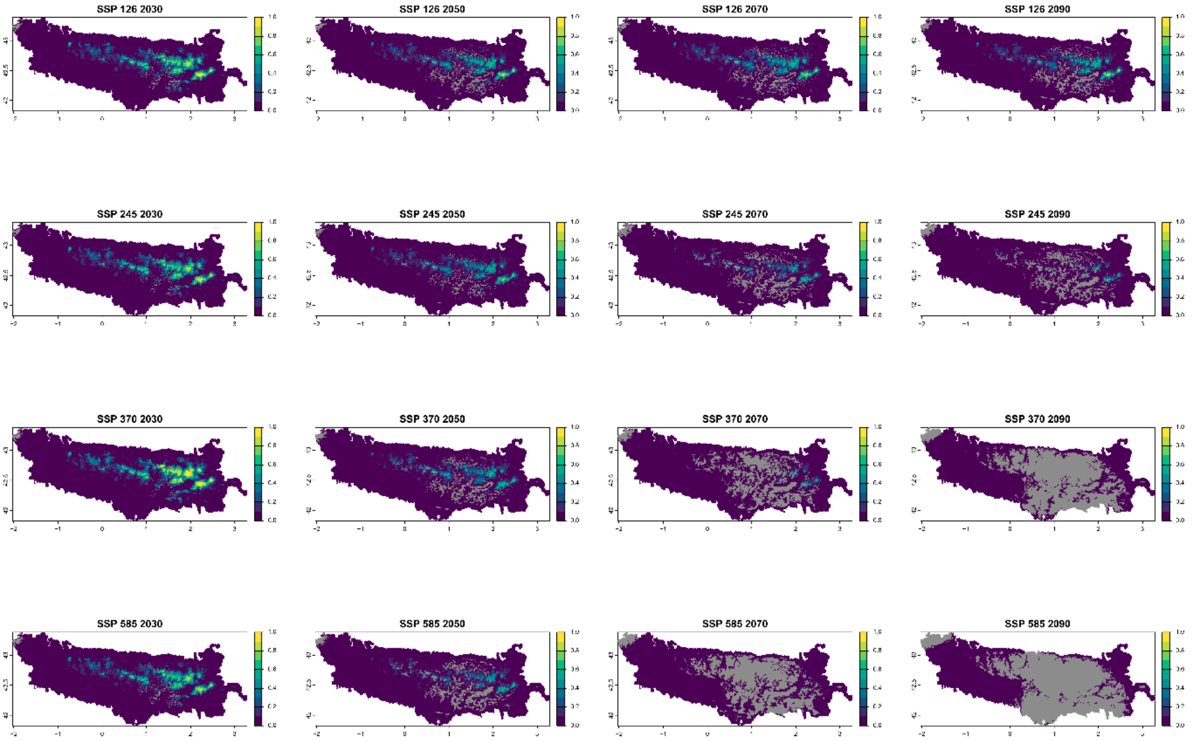


■ Legend: Non−analogous bioclimatic conditions (MESS < 0)

Current bioclimatic suitability in the Pyrenees for *Saxifraga hariotii*

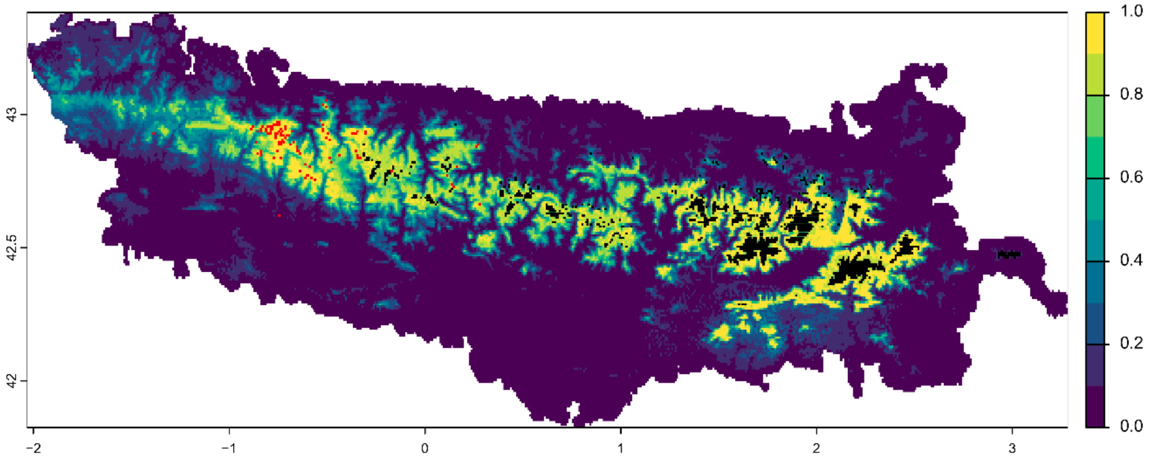


Symbols:

- ■ Variability >=5% among predictions
- ■ Field observation

Future bioclimatic suitability in the Pyrenees by 2090 for *Saxifraga hariotii*

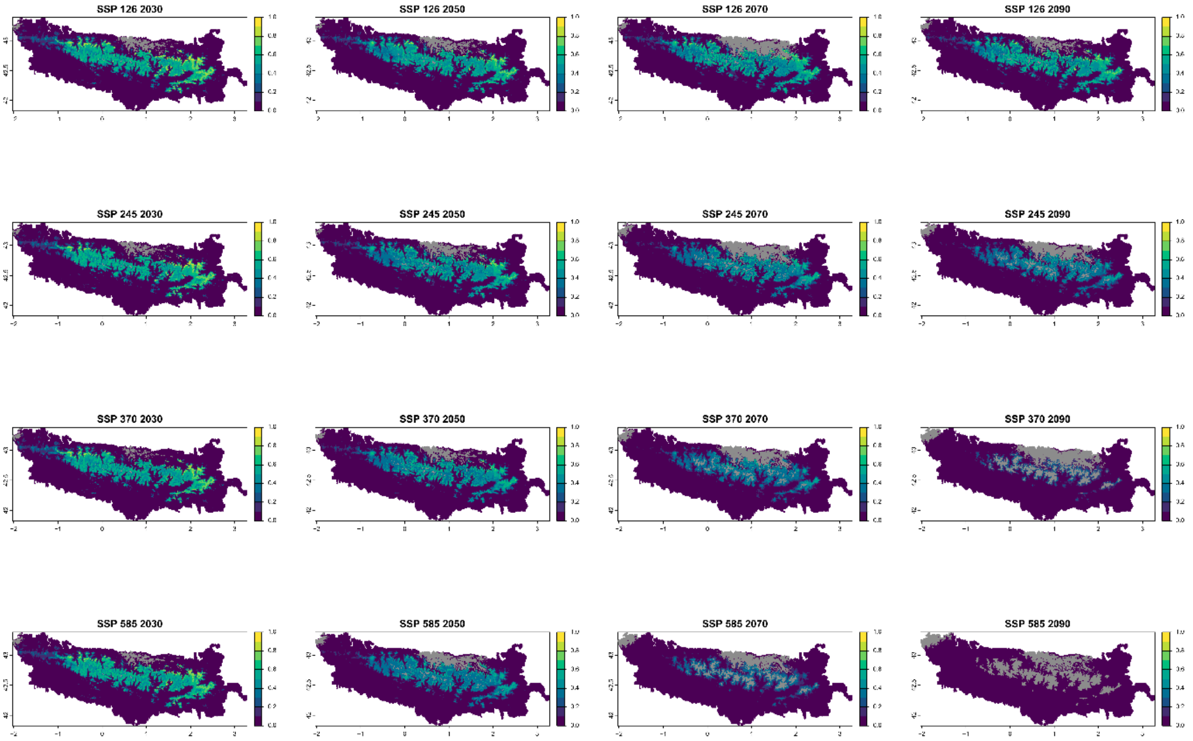


■ Legend: Non−analogous bioclimatic conditions (MESS < 0)

Current bioclimatic suitability in the Pyrenees for *Saxifraga intricata*

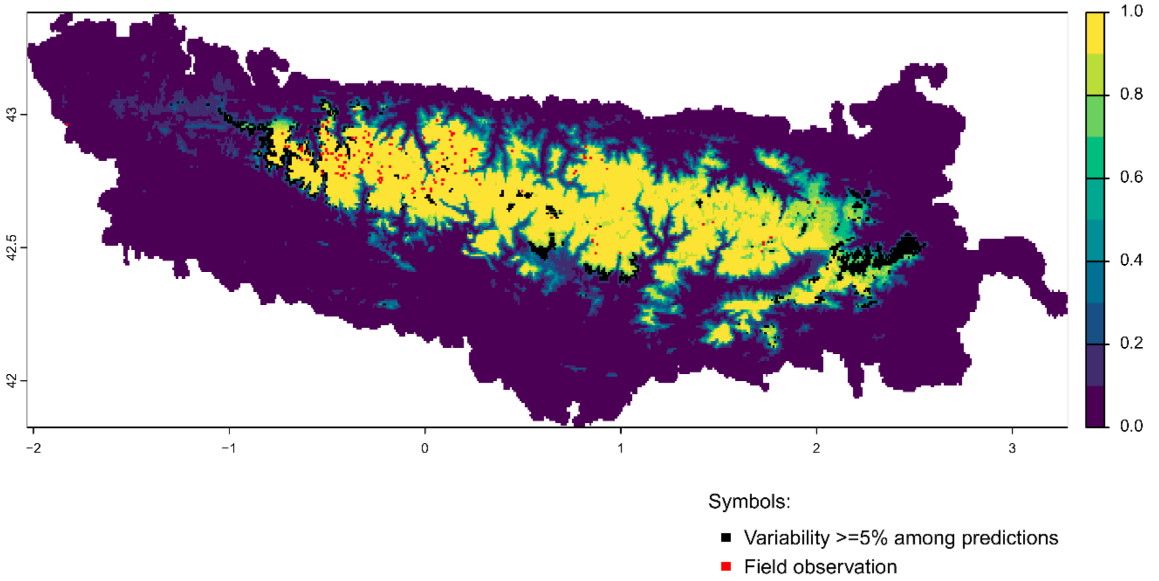


Future bioclimatic suitability in the Pyrenees by 2090 for *Saxifraga intricata*

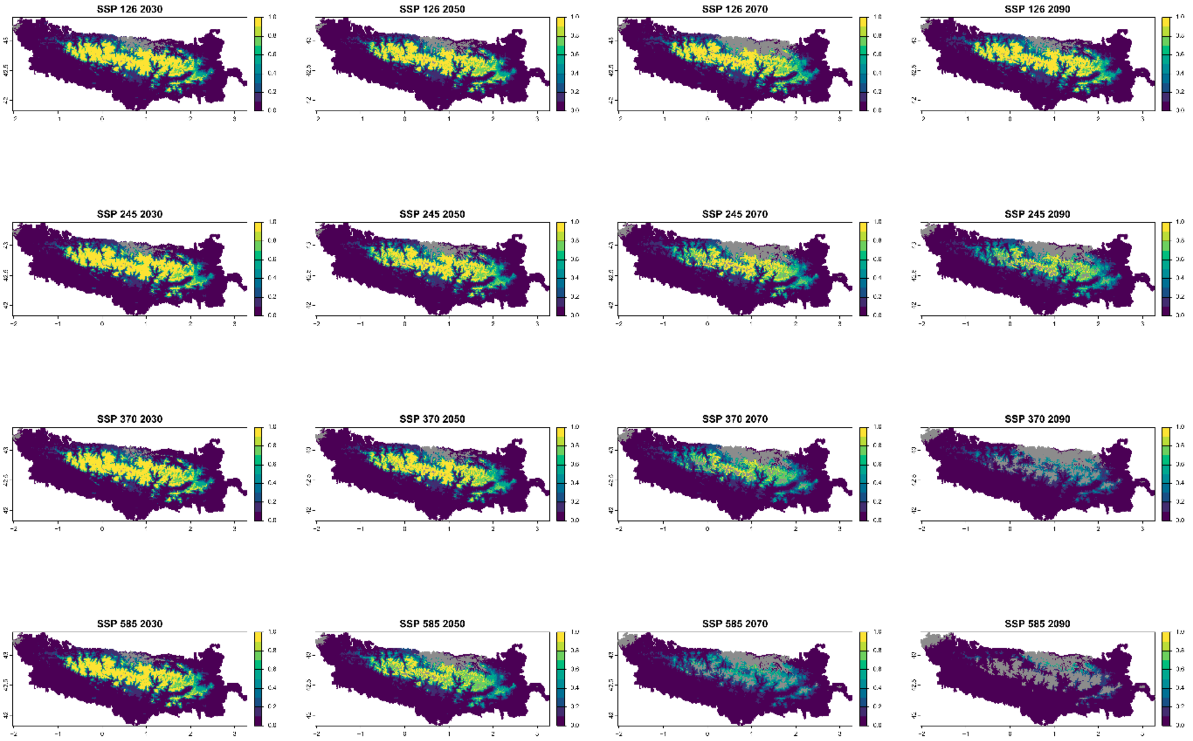


Legend: Non−analogous bioclimatic conditions (MESS < 0)

Current bioclimatic suitability in the Pyrenees for *Saxifraga media*

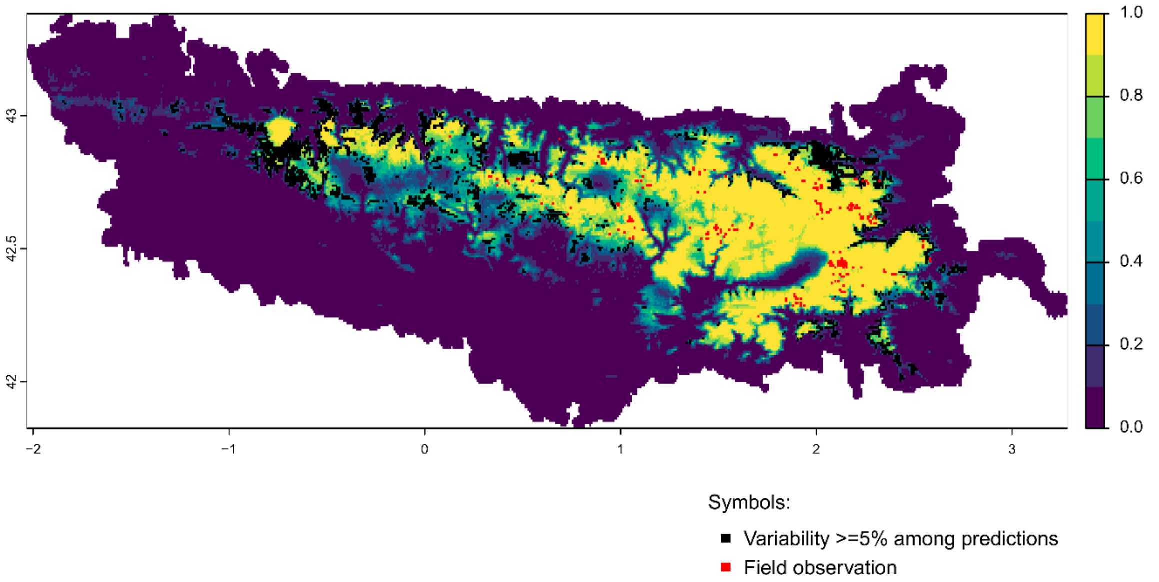


Future bioclimatic suitability in the Pyrenees by 2090 for *Saxifraga media*

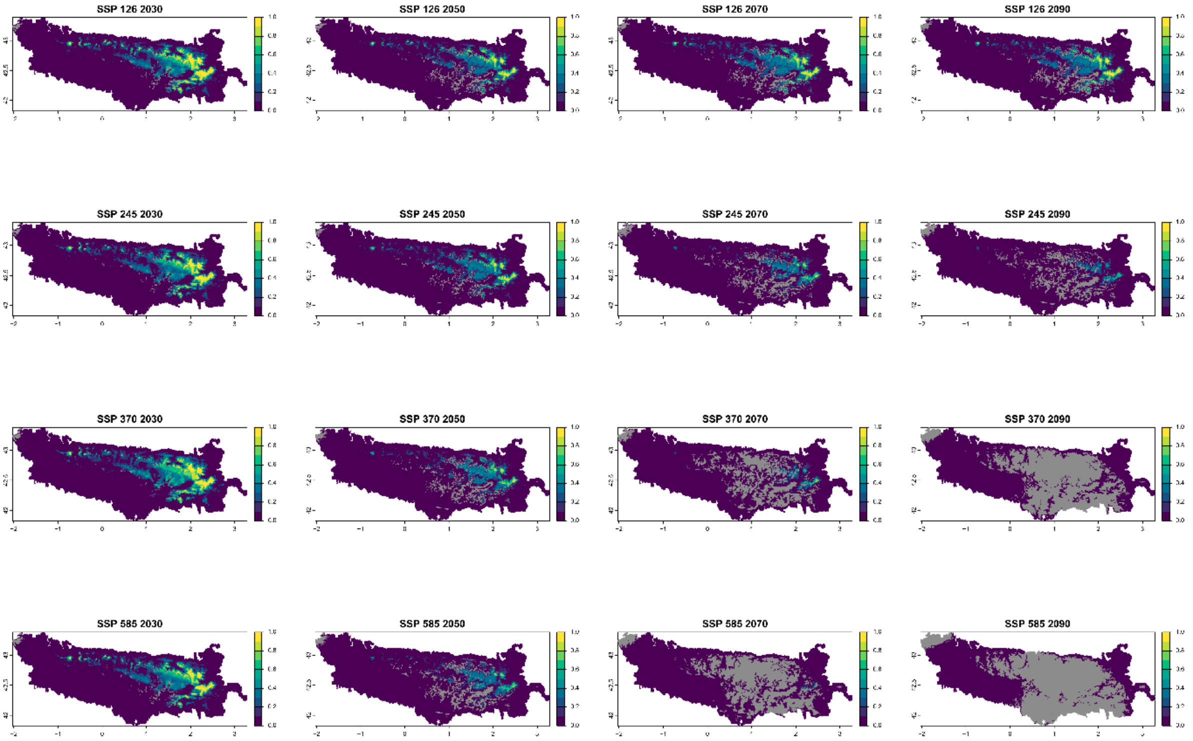


Legend: Non−analogous bioclimatic conditions (MESS < 0)

Current bioclimatic suitability in the Pyrenees for *Saxifraga umbrosa*

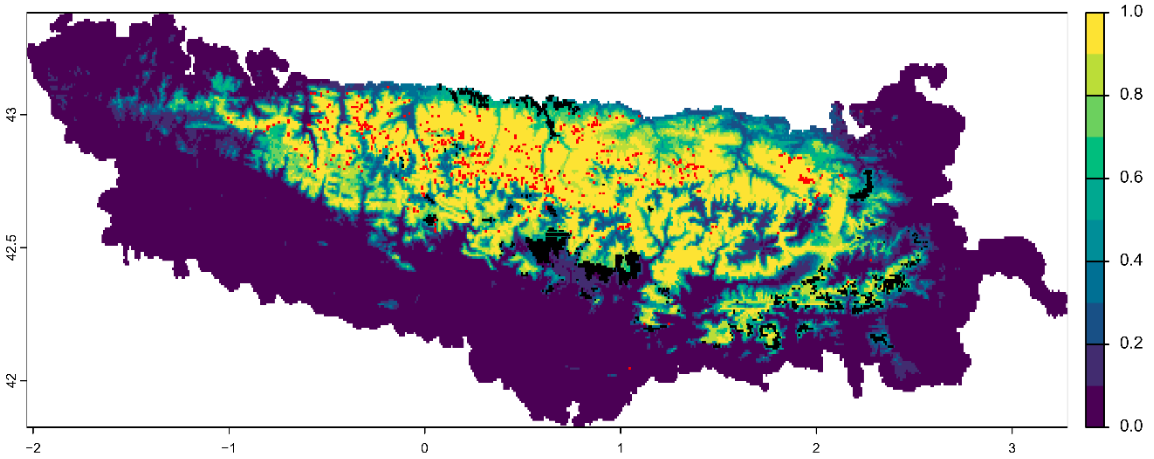


Symbols:

- ■ Variability >=5% among predictions
- ■ Field observation

Future bioclimatic suitability in the Pyrenees by 2090 for *Saxifraga umbrosa*

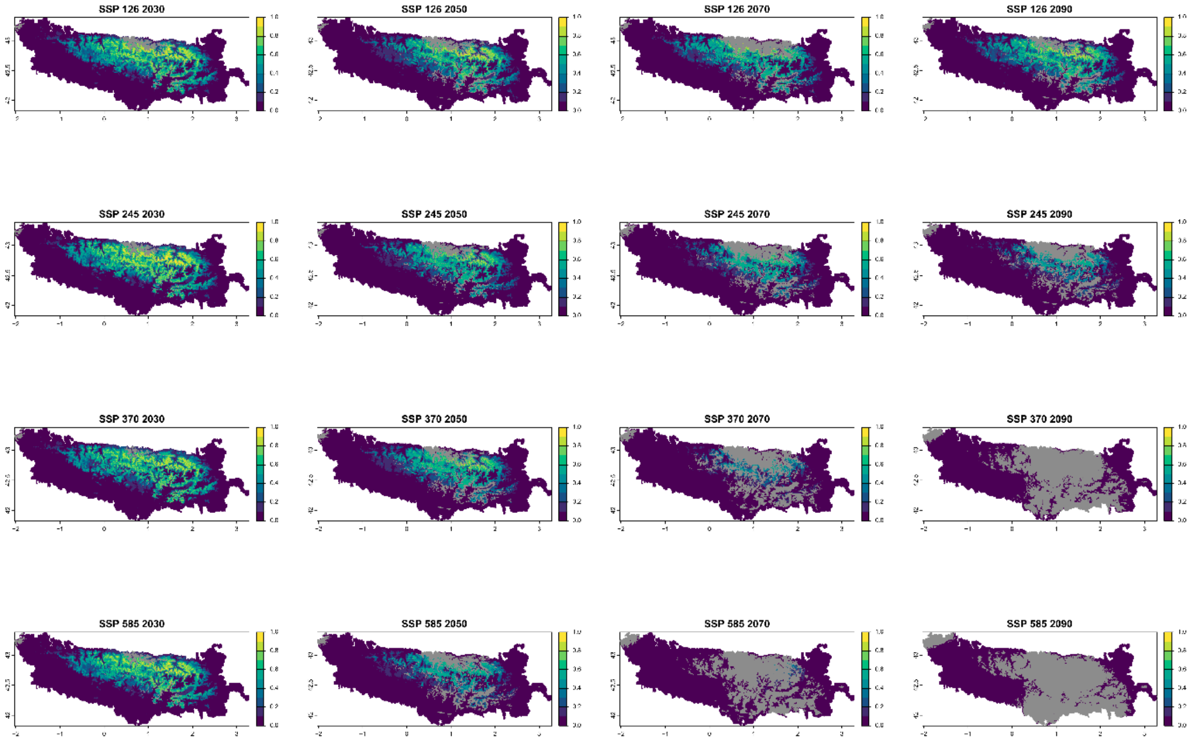


■ Legend: Non−analogous bioclimatic conditions (MESS < 0)

### Current bioclimatic suitability in the Pyrenees for *Scrophularia pyrenaica*

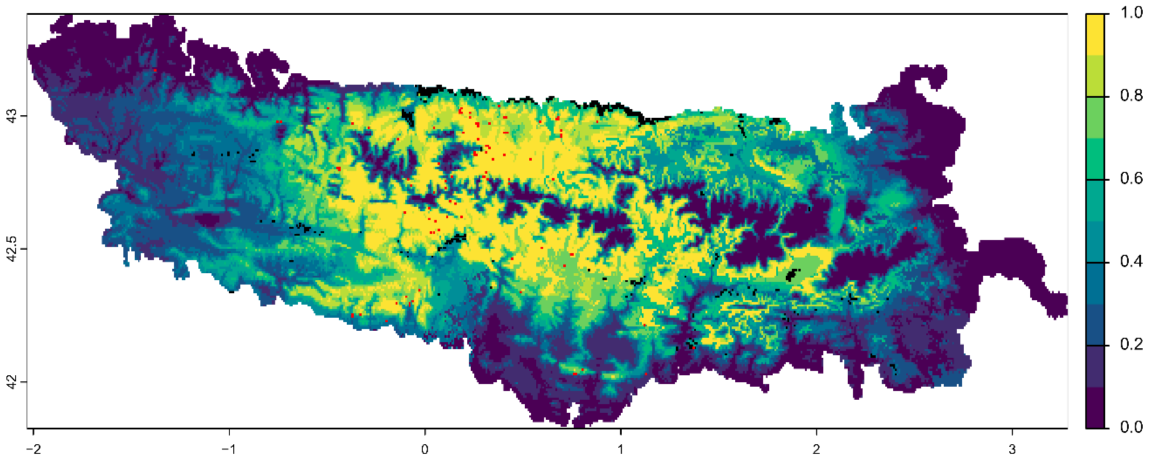


Symbols:

- Variability >=5% among predictions
- Field observation

#### Future bioclimatic suitability in the Pyrenees by 2090 for *Scrophularia pyrenaica*

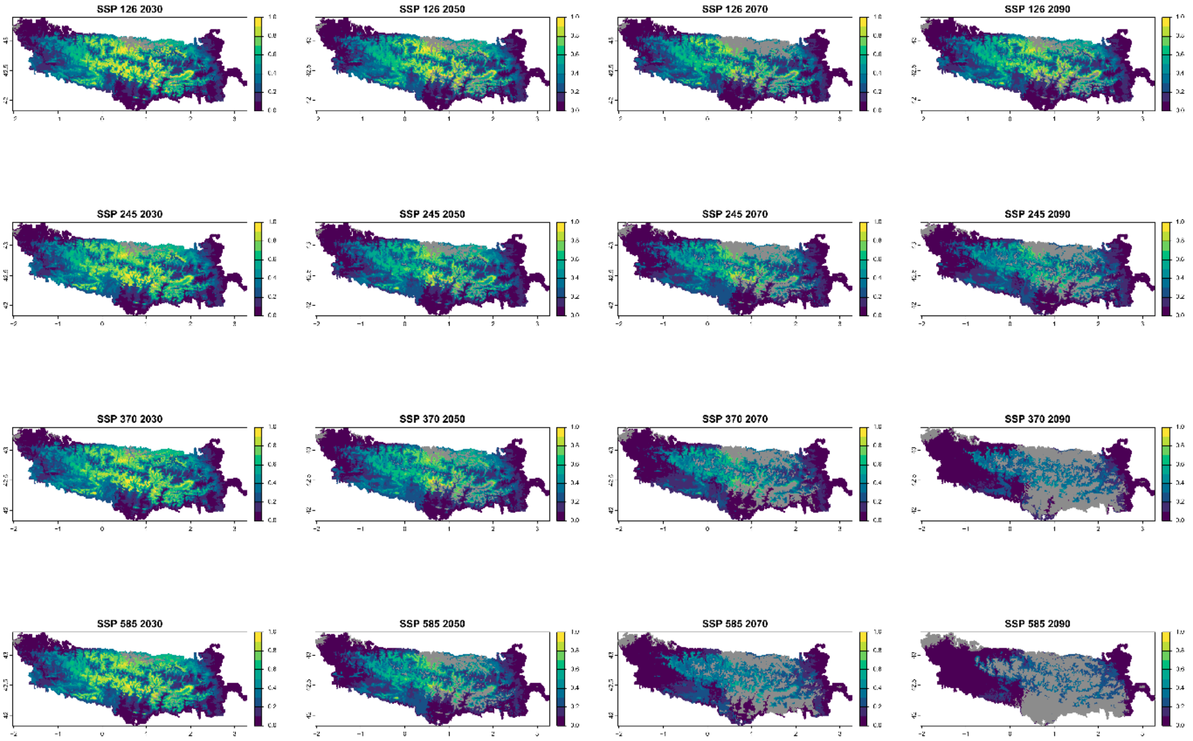


Legend: Non−analogous bioclimatic conditions (MESS < 0)

Current bioclimatic suitability in the Pyrenees for *Silene borderei*

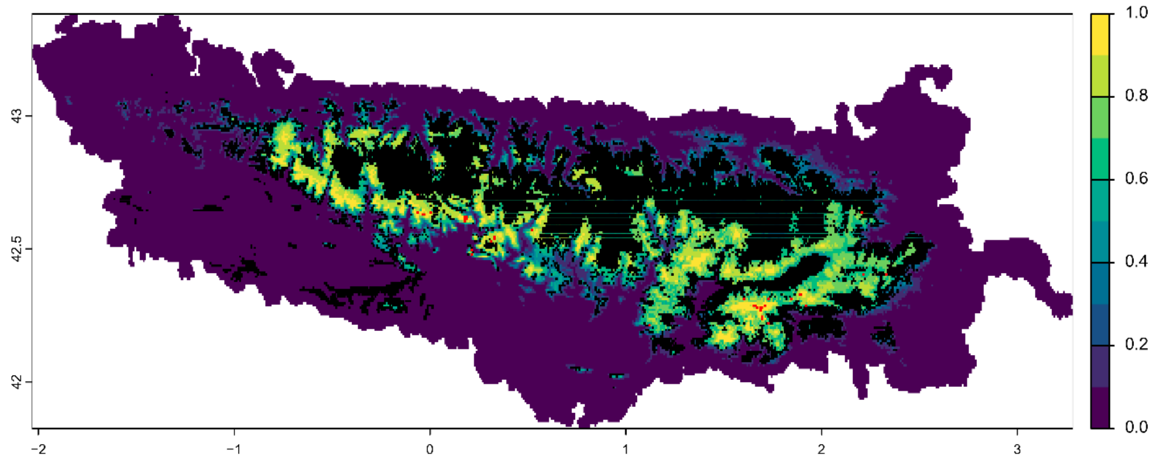


Symbols:

■ Variability >=5% among predictions

■ Field observation

Future bioclimatic suitability in the Pyrenees by 2090 for *Silene borderei*

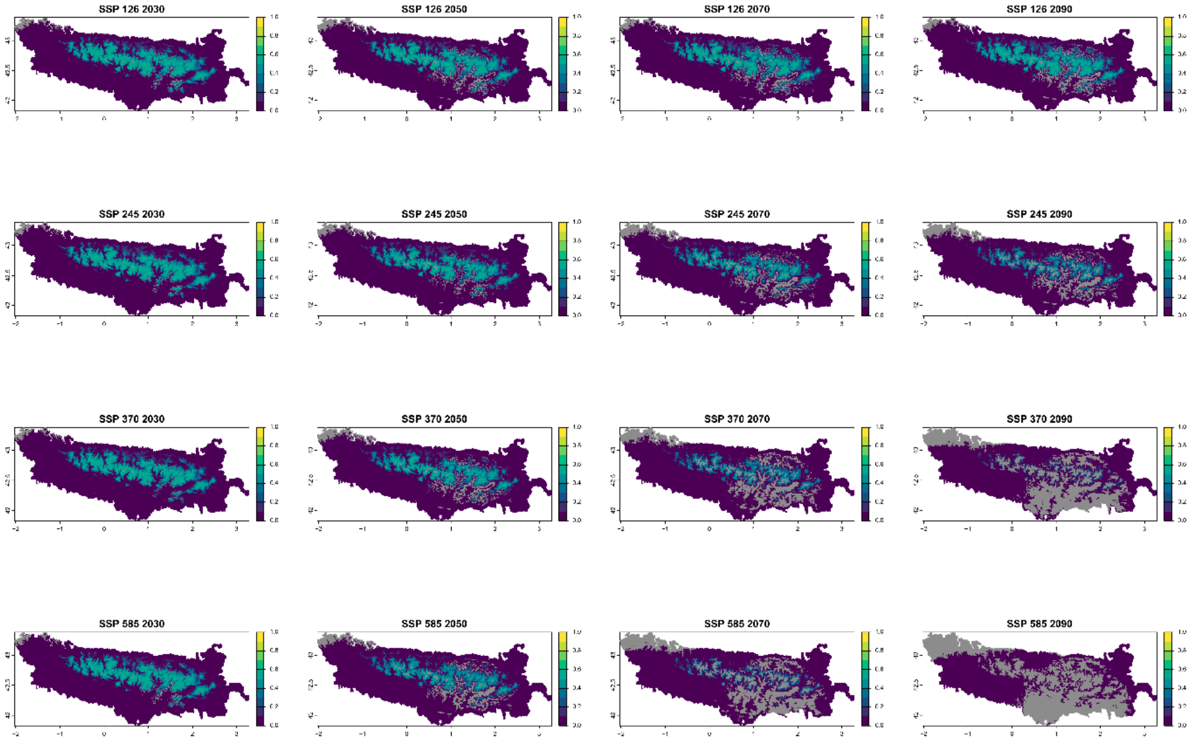


■ Legend: Non−analogous bioclimatic conditions (MESS < 0)

Current bioclimatic suitability in the Pyrenees for *Thalictrum macrocarpum*

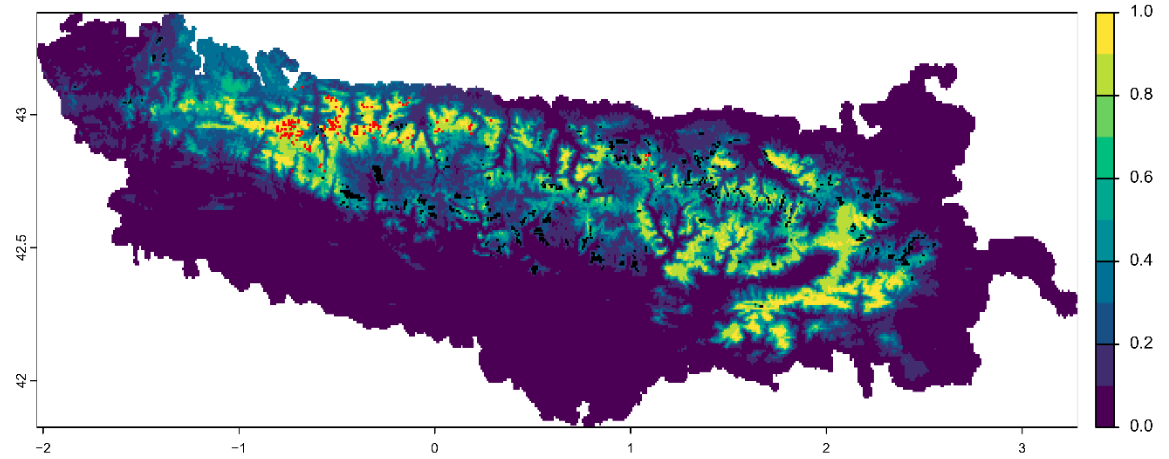


Symbols:

- ■ Variability >=5% among predictions
- ■ Field observation

Future bioclimatic suitability in the Pyrenees by 2090 for *Thalictrum macrocarpum*

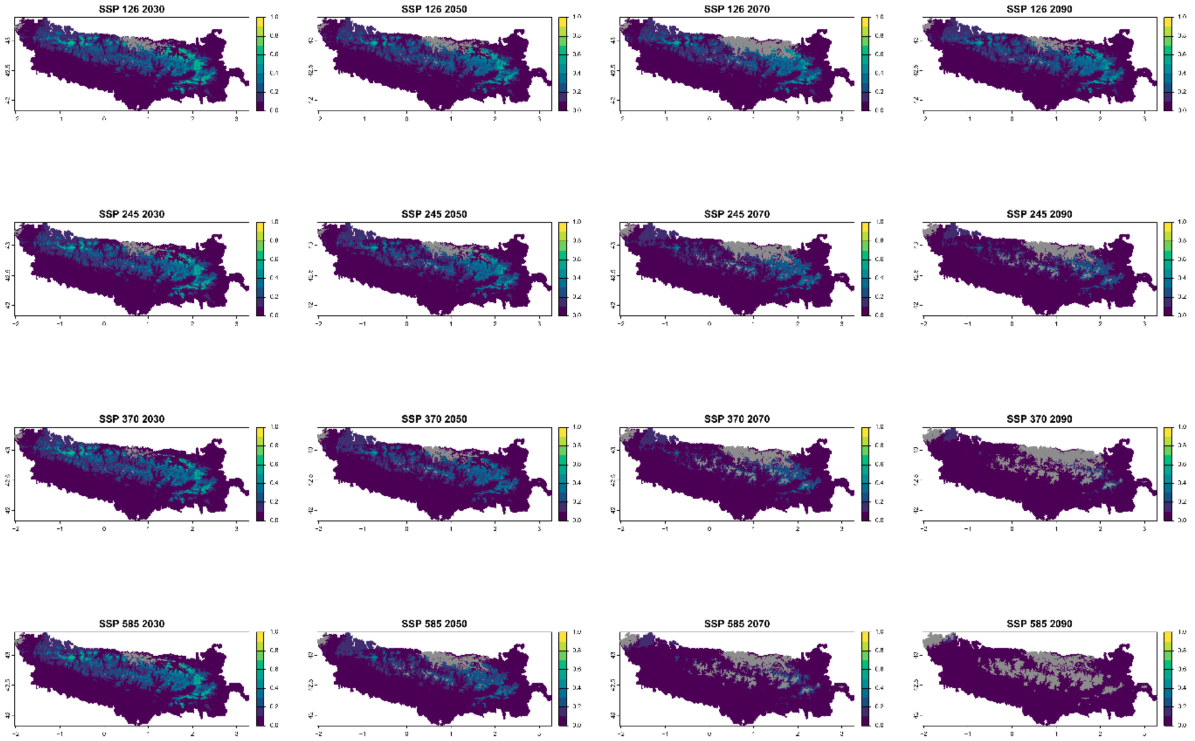


■ Legend: Non−analogous bioclimatic conditions (MESS < 0)

Current bioclimatic suitability in the Pyrenees for *Thymelaea calycina*

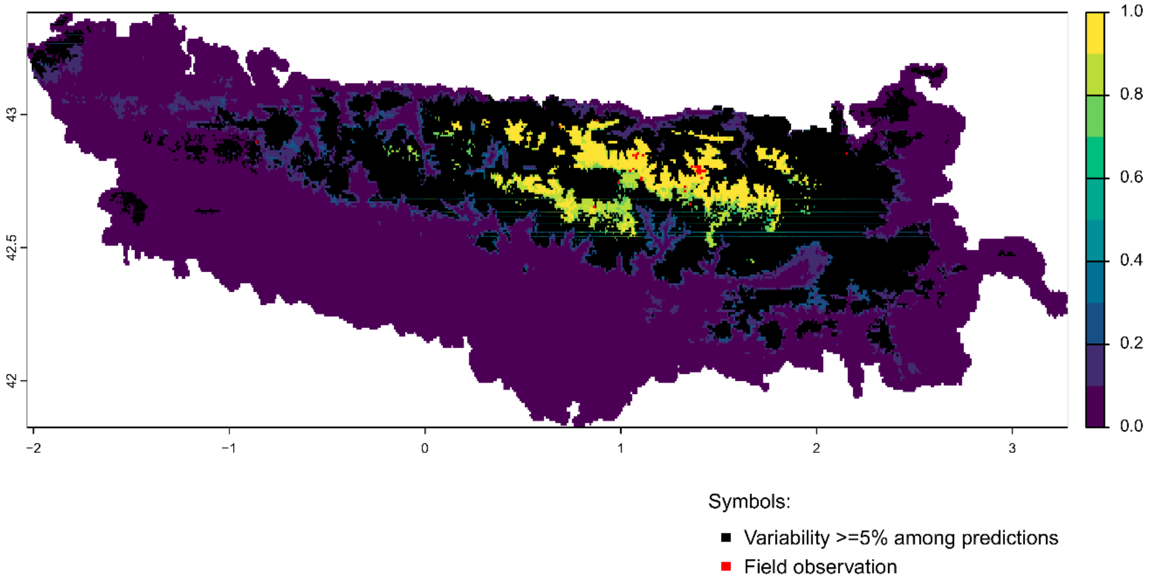


Future bioclimatic suitability in the Pyrenees by 2090 for *Thymelaea calycina*

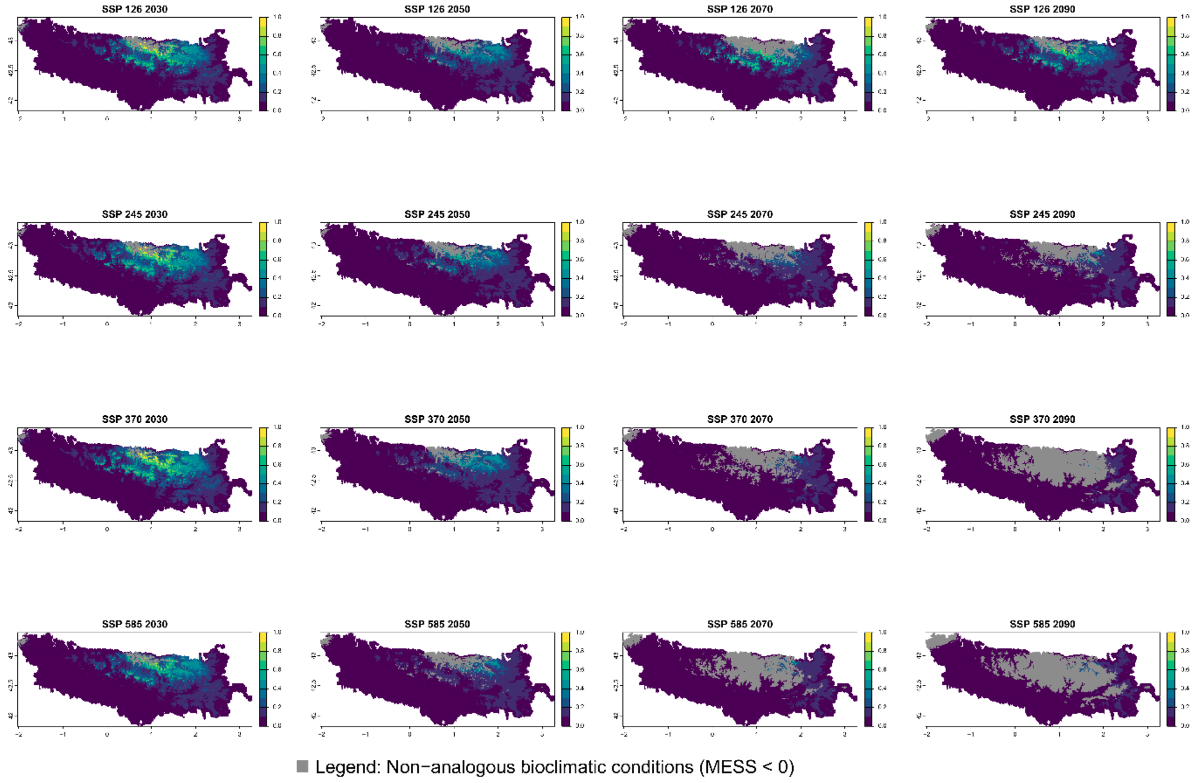

Current bioclimatic suitability in the Pyrenees for *Trisetum baregense*

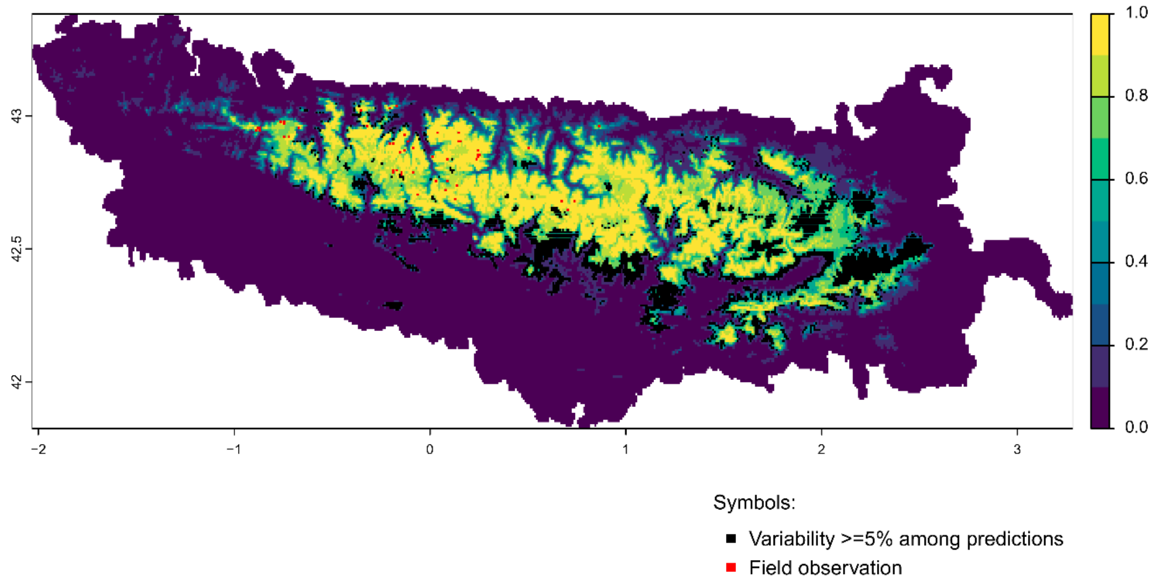


Future bioclimatic suitability in the Pyrenees by 2090 for *Trisetum baregense*

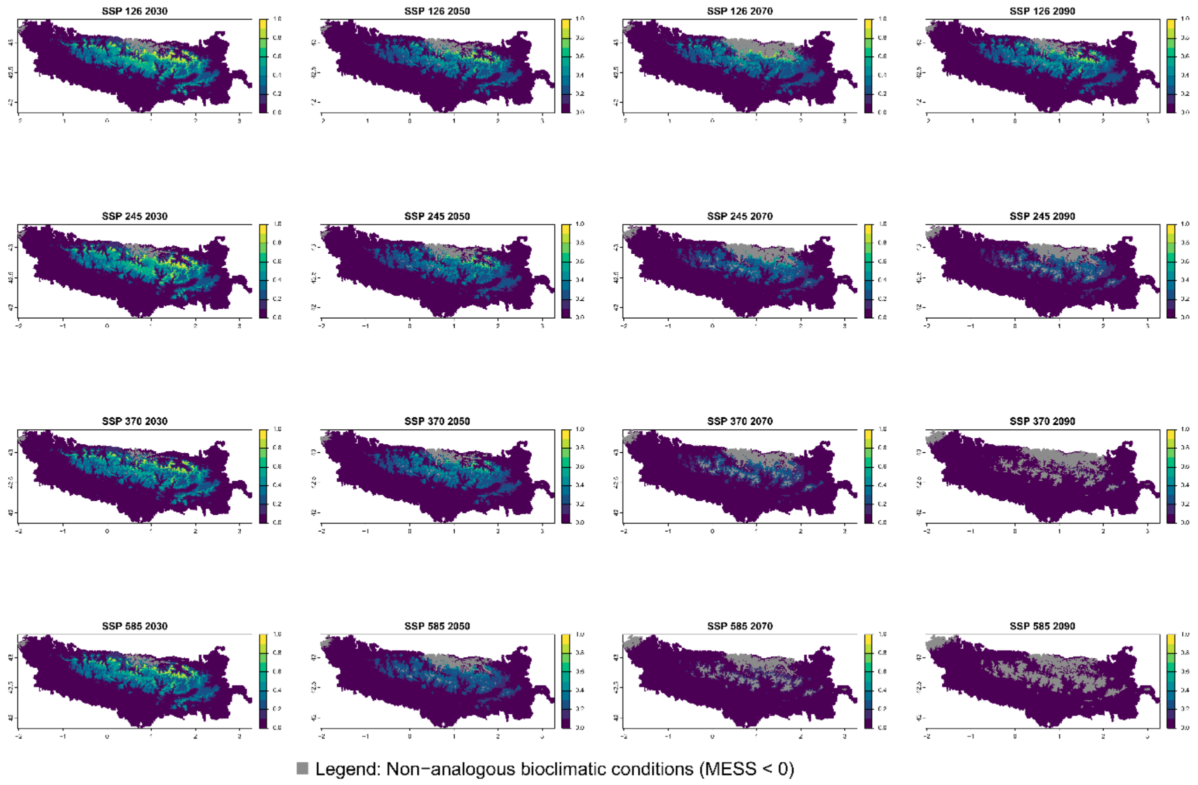

### Current bioclimatic suitability in the Pyrenees for *Viola diversifolia*

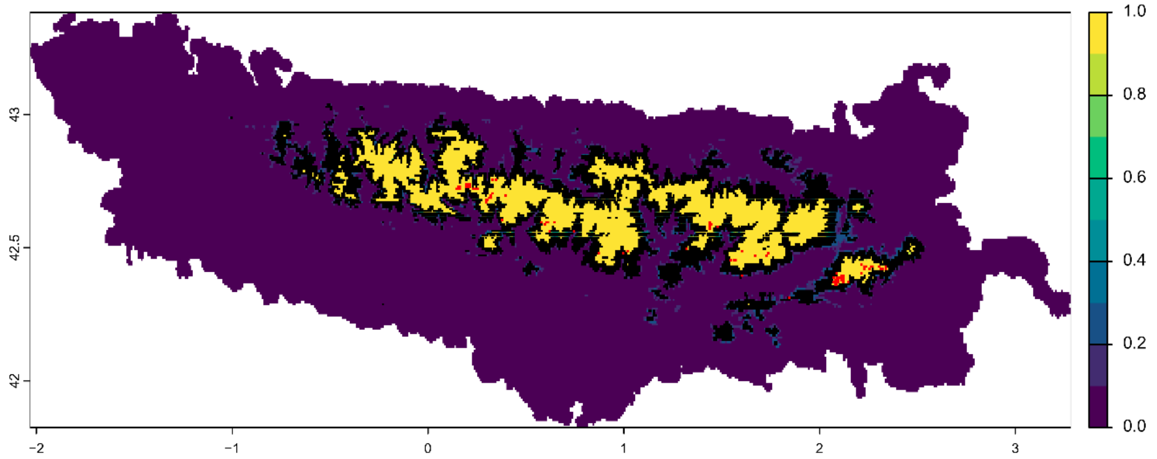


Symbols:

- ■ Variability >=5% among predictions
- ■ Field observation

### Future bioclimatic suitability in the Pyrenees by 2090 for *Viola diversifolia*

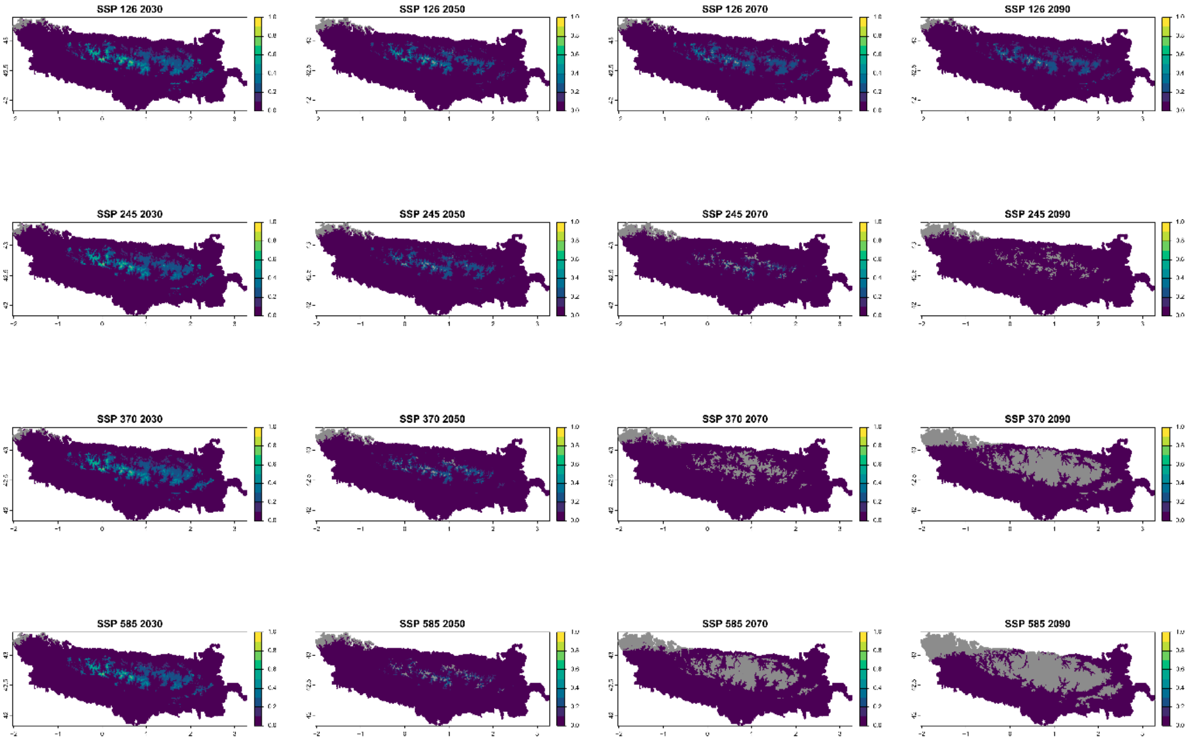


■ Legend: Non−analogous bioclimatic conditions (MESS < 0)

Current bioclimatic suitability in the Pyrenees for *Xatartia scabra*

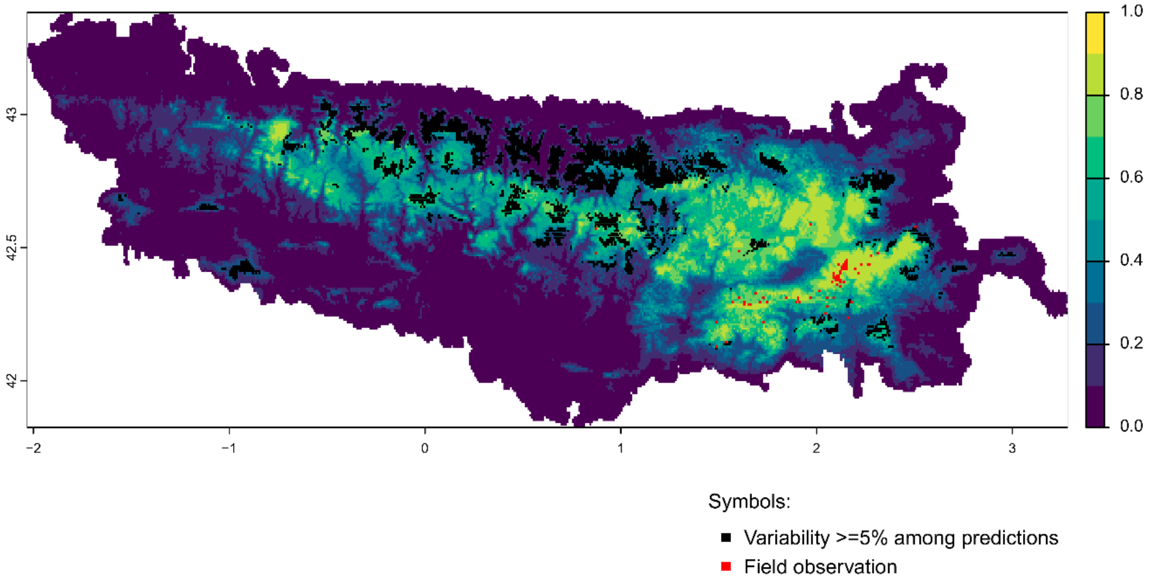


Future bioclimatic suitability in the Pyrenees by 2090 for *Xatartia scabra*

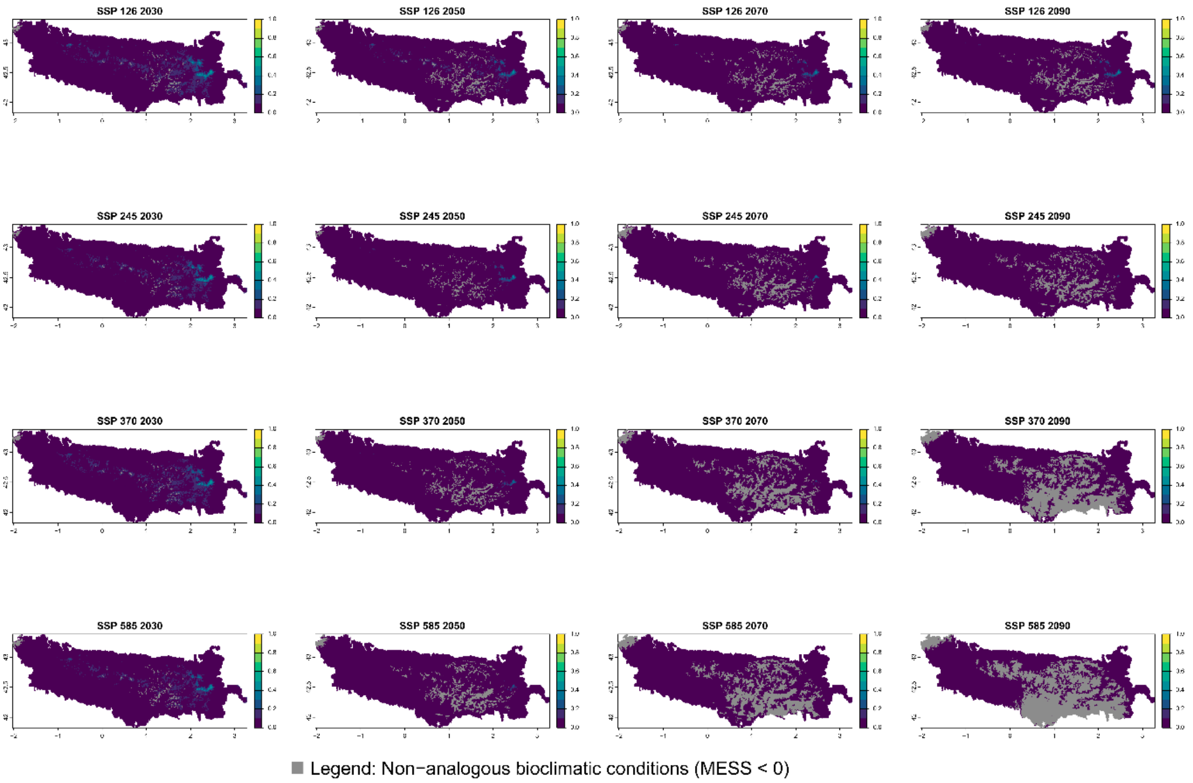
